# Epigenetic derepression of H3K9me3 mitigates Alzheimer-related pathology and improves cognition via immunomodulation and Vgf induction

**DOI:** 10.1101/2024.10.15.618518

**Authors:** Dieu-Trang Fuchs, Jean-Philippe Vit, Altan Rentsendorj, Julia Sheyn, Haoshen Shi, Yosef Koronyo, Oksana Chepurna, Miyah R. Davis, Jered W. Wilson, Lon S. Schneider, Stuart L. Graham, Vivek K. Gupta, Margaret Fahnestock, Tao Sun, Michael T. Kleinman, David Horne, Mehdi Mirzaei, Keith L. Black, Maya Koronyo-Hamaoui

## Abstract

We investigated the role of histone 3 lysine 9 trimethylation (H3K9me3), an epigenetic mechanism involved in the repression of synaptic plasticity and memory-related genes, within aging and Alzheimer’s disease (AD). Our study reveals that elevated cortical H3K9me3 strongly correlates with cognitive dysfunction in individuals with mild cognitive impairment (MCI) and AD. In old (18 months) and younger (14 months) APPSWE/PS1ΔE9 and 3xTg AD mouse models, inhibiting SUV39H1 methyltransferase with ETP69, substantially reduces cerebral H3K9me3 levels and attenuates amyloid-β burden, tau pathology, and gliosis. Administration of ETP69 further promotes dendritic spine formation, leading to rapid and sustained improvements in cognitive function. Proteomics analysis indicates that a significant proportion of dysregulated proteins in the brains of AD-model mice are reversed by ETP69. These proteins are enriched for synaptic plasticity and learning-related pathways. ETP69 exerts its effects through multiple neuroprotective mechanisms, including regulation of neuroinflammation, induction of both blood and cerebral-infiltrating monocytes involved in cerebral Aβ clearance. Moreover, ETP69 activates brain-derived neurotrophic factor (Bdnf) network, and particularly its downstream effector neurosecretory protein Vgf. These findings support the pharmacological inhibition of H3K9me3-mediated gene silencing to reverse AD-related pathology and cognitive decline.

## INTRODUCTION

Alzheimer’s disease (AD) is the world’s most prevalent type of senile dementia that is characterised by debilitating loss of memory and executive functions.^1^ The pathological hallmarks of AD, amyloid-β protein (Aβ) plaques and neurofibrillary tangles (NFTs) comprising hyperphosphorylated tau protein, are believed to induce neuroinflammation and vascular damage, ultimately leading to severe synaptic and neuronal loss.^2–4^ Besides known genetic risk factors, advanced age is a primary contributor to the development of AD.^5,6^

Epigenetic modifications of histones, namely methylation and acetylation, reshape chromatin structure and influence gene expression throughout the lifespan.^7,8^ Notably, these processes participate in the transcriptional regulation of synaptic plasticity–related genes such as brain-derived neurotrophic factor (*Bdnf*) and are involved in memory formation and consolidation.^9–12^ Epigenetic alterations and loss of chromatin dynamics occur in the ageing and neurodegenerative brain.^13–17^ The reversible nature of epigenetic modifications makes histone-modifying enzymes promising drug targets for improving brain functions in the presence of old age and neurological diseases.^18,19^

Methylation of position-specific lysine residues in histone tails is associated with either active or repressed chromatin.^20,21^ Dimethylation and trimethylation of the lysine 9 residue of histone 3 (H3K9me2/3) are two major epigenetic marks of transcriptional repression that have been linked to ageing and shown to increase in the brains of AD patients and AD-model mice.^22–24^ Notably, the elevation of H3K9me3, catalysed by the histone methyltransferase suppressor of variegation 3–9 homologue 1 (SUV39H1), is associated with reduced transcription of genes governing synaptic function, including *Bdnf.*^23^ In old wild-type (WT) mice, an early study suggested that selective inhibition of SUV39H1 by ETP69 (*also named* NT1721), a synthetic epidithiodiketopiperazine based on a natural product,^25^ reduce ageing-related memory deficits.^26^ However, whether and how SUV39H1 inhibition by ETP69 can attenuate Alzheimer’s-related pathology and cognitive decline in the context of AD, has never been investigated.

In this study, we explored the abundance of cerebral H3K9me3 in donor patients with diagnosis of mild cognitive impairment (MCI, due to AD) or AD dementia as compared to individuals with normal cognition. Then, we determined the relationship of brain H3K9me3 levels with severity of Alzheimer’s pathology and cognitive status. Further, the therapeutic potential and mode of action of ETP69 was investigated in old (18 months) and younger adult (14 months) WT, APP_SWE_/PS1_ΔE9_-transgenic (AD^+^) and APP_SWE_/tauP301L/PS1^tm1Mpm^ (3xTg AD^+^) model mice by performing histological, neuronal architecture analysis, proteome profiling, and multidomain behavioural testing. We found that targeting H3K9me3 epigenetic repression by ETP69 administration mitigated AD-related neuropathology, including paired-helical filaments (PHF) of tau and Aβ plaques, along with fully restoring dendritic spine formation, synaptic markers, and cognitive functions. The neuroprotective mechanisms of ETP69 involved modulation of innate immune responses as well as activation of the Bdnf signalling pathway and, especially, the neurosecretory protein Vgf.

## MATERIALS AND METHODS

### Mice

Double-transgenic B6.Cg-Tg (APP_SWE_, PSEN1_ΔE9_) 85Dbo/Mmjax AD (AD^+^) mice and their age-matched WT C57BL/6J littermates were obtained from the Mutant Mouse Resource and Research Center (MMRRC) at the Jackson Laboratory (RRID:MMRRC_034832-JAX), an NIH-funded strain repository, and were donated to the MMRRC by David Borchelt, Ph.D. (McKnight Brain Institute, University of Florida). Mice were then bred and maintained at Cedars-Sinai Medical Center. housed up to five per cage on a 12-h light/dark cycle and provided with food and water *ad libitum*. All animals used in this study had a congenic C57BL/6 background. Both male and female mice were used for all experiments and were assigned to experimental groups after balancing for age and genotype. Experiments were approved by the Institutional Animal Care and Use Committee (IACUC) of Cedars-Sinai Medical Center, Los Angeles, CA.

Female triple-transgenic B6;129-Tg (APP_SWE_,tauP301L)1Lfa *Psen1^tm1Mpm^* (3xTg AD^+^) mice (Jackson Laboratories, MMRRC_034830-JAX) were maintained at University of California, Irvine (UCI) under an approved IACUC protocol.

All experiments were performed according to the NIH Guidelines for the Care and Use of Laboratory Animals and the study was conducted in compliance with ARRIVE guidelines.

Animal group allocation, experimental procedures and data analyses were performed by different experimenters who were blinded to genotypes and treatments throughout this study.

### ETP69 treatment regimen

Lyophilized ETP69 was resuspended in 50% DMSO (in saline) at a concentration of 10 mg/ml. This ETP69 solution was administered to mice by i.p. injection at a dose of 10 mg/kg (ETP69-treated groups). DMSO (50%, in saline) was injected for use as a control (DMSO-injected groups). Five cohorts of WT and AD^+^ mice (*n*=153; 14 and 18 months old) were treated with either ETP69 or DMSO according to three different regimens (regimens S, R, and B). In 18-month-old mice (*n*=118), two cohorts of AD^+^ and WT mice received a single ETP69 dose (regimen S) 1 day before the start of behavioural testing (day 0) and were euthanized either on day 4 or day 15. A third group of AD^+^ and WT mice received repeated ETP69 doses (once weekly for 11 weeks) with the last administration on day 0 (regimen R). A fourth group of AD^+^ and WT mice received the first dose of ETP69 on day 0 and a booster dose on day 9, the day before memory retention testing involving the Barnes maze (regimen B). In 14-month-old mice (n=35), AD^+^ and WT mice received a single ETP69 dose (regimen S) on day 0 and were euthanized on day 4. Control AD^+^ and WT mice received DMSO according to the same regimens S, R, or B.

One cohort of 3xTg AD^+^ mice (*n*=11; 14 months old) received a single injection of ETP69 or DMSO on day 0 and were euthanized on day 7. Timelines of behavioural tests and experimental endpoints are presented in respective figures.

### Behavioural tests

All animals in this study underwent behavioural tests.

### Open field test

Spontaneous locomotor activity was assessed for 30 min using the Photobeam Activity System (www.sandiegoinstruments.com). Ambulatory and rearing activities were recorded, and distance, speed, and resting time were calculated.

### Barnes maze test

Mice were first trained to locate an escape box in a 20-hole circular table during a 4-min trial that was performed 3 times per day for 4 days (acquisition training phase), as previously described.^27^ Following a 2-day break, the memory retention of each mouse was evaluated on day 7 (retention phase). Memory extinction and learning of a new escape location were assessed on days 8–9 (reversal phase). The latency to find the escape box and the number of incorrect entries (errors) were recorded for each trial and averaged on each day for each mouse.

### Y-maze spontaneous alternation test

The Y-shaped apparatus used for this study consists of two arms equal in length and one longer arm. Mice were individually placed at the distal end of the long arm and allowed to move freely throughout the entire maze (all three arms) for 5 min under dim light. The sequence of arm entries and the total number of entries were recorded; the percentage of spontaneous alternations was calculated as follows:

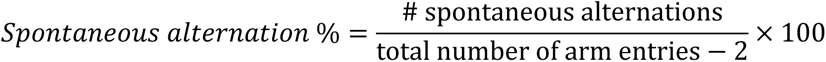

A spontaneous alternation is defined as the sequential visit to the three different arms without returning to a previously visited one.

### Visual-stimuli X-maze test

To assess spontaneous behaviour induced by color (under equal conditions) and contrast sensitivity, the mice were tested using our custom-made ViS4M, as previously described.^28,29^ For each mode, the mice were individually placed in the centre of the ViS4M and allowed to freely explore the maze for 5 min. The sequences of arm entries were manually documented according to the video recordings. The total number of entries, percentage of bidirectional transitions between arms, and percentage of alternations were all determined from the sequences of arm entries.

Chord diagrams were generated to visualize behavioural data from the Barnes maze and ViS4M tests using the free online resource Circos (mkweb.bcgsc.ca/tableviewer/) as previously described.^28,29^

### Context-specific fear conditioning test

During the acquisition phase, an animal was placed in a freezing behaviour-monitoring chamber (www.sandiegoinstruments.com) and allowed to habituate for 2 min before receiving a 0.2-mA electric foot shock for 1 s. The animal remained in the chamber for an additional 3 min and was then returned to its home cage. To assess context-specific fear, the animal was placed in the same chamber 24 h after the acquisition phase, and the freezing time (absence of movement for 3 s) over a 4-min session was recorded.

All behavioural tests were performed by an experimenter blinded to mouse genotypes and treatments.

### Brain collection and processing

After completion of the behavioural tests, on either day 4 or day 15, all mice were placed in a state of deep anaesthesia (50 mg/kg ketamine/xylazine) and transcardially perfused with ice-cold saline solution containing 0.5 mM EDTA. Mouse brains were collected and processed. Tissues from the brain regions shown in Figs. 2F, 3B, and 4B were either snap frozen and then stored at −80°C for protein extraction, fixed in 2.5% PFA overnight, and then cryoprotected in 60% sucrose for IHC or processed for Golgi-Cox staining.

**Fig. 1.**
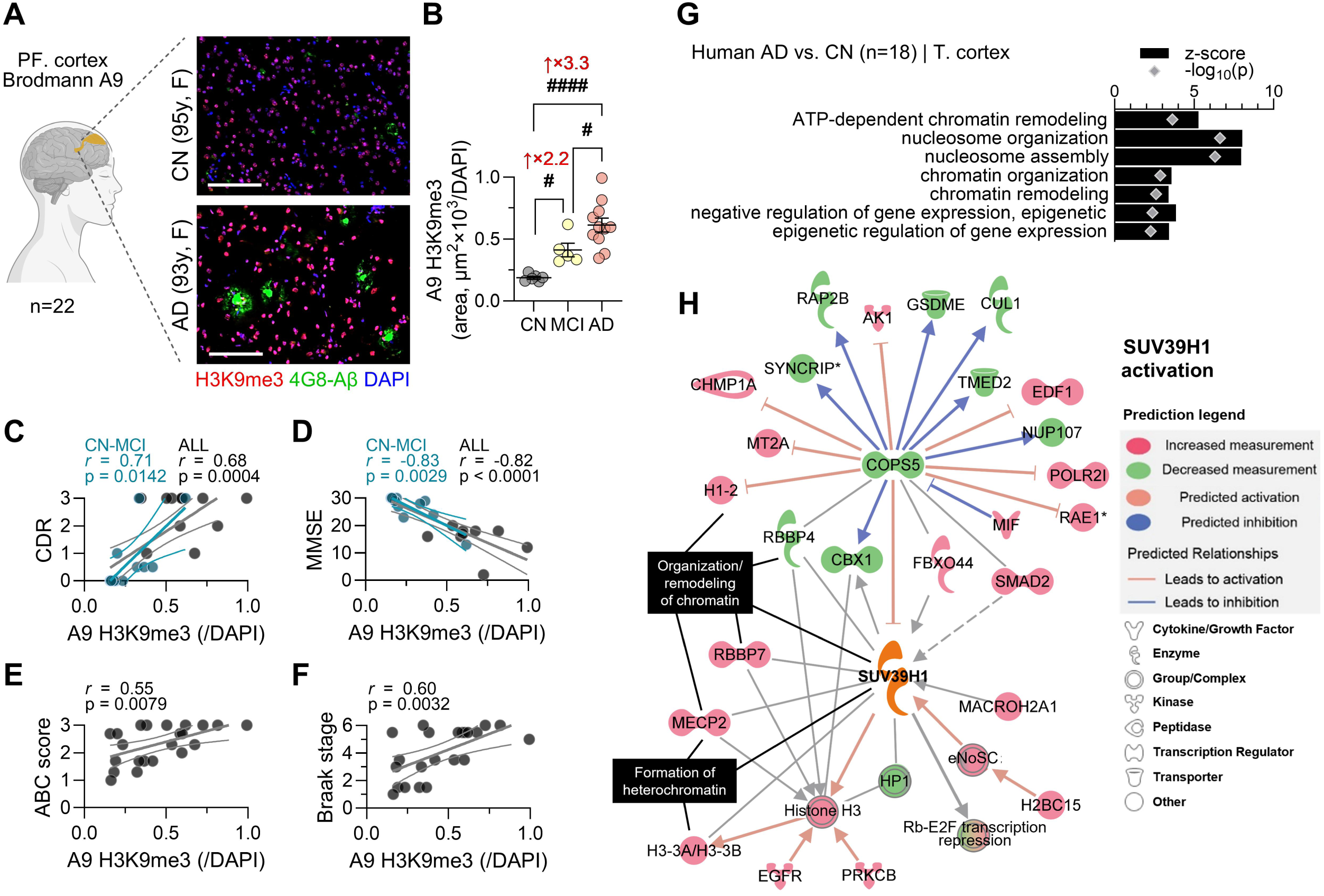
Increased cerebral H3K9me3 correlates with cognitive deficits and AD-related neuropathology. (**A**) Representative images of fluorescent immunostaining (scale bar: 100 μm) for H3K9me3 (red), 4G8 (green, Aβ plaques), and DAPI (blue, nuclei) in the dorsolateral prefrontal cortices (Brodmann area A9) of patients with AD or with mild cognitive impairment (MCI), and cognitively normal subjects (CN). (**B**) Quantification of H3K9me3 immunoreactivity corrected for DAPI nuclei counts (area, μm^2^×10^3^/DAPI). # *p* < 0.05 and #### *p* < 0.0001, by one-way ANOVA followed by Fisher’s LSD *post hoc* test. (**C–F**) Pearson’s correlation coefficients of A9 H3K9me3/DAPI levels with (C) clinical dementia rating (CDR), (D) mini mental state examination (MMSE), (E) ABC scores, and (F) Braak stages. Correlations limited to CN-MCI individuals are shown in blue for CDR and MMSE. (**G**) Gene Ontology analysis of upregulated DEPs in the temporal cortex of AD patients pertaining to chromatin organization and epigenetic regulation of gene expression. (**H**) Expression profiles of select first and second neighbours upstream and downstream of SUV39H1 predicting its activation in the brain of AD patients, according to IPA. All up and downregulated DEPs were included in this analysis.

**Fig. 2.**
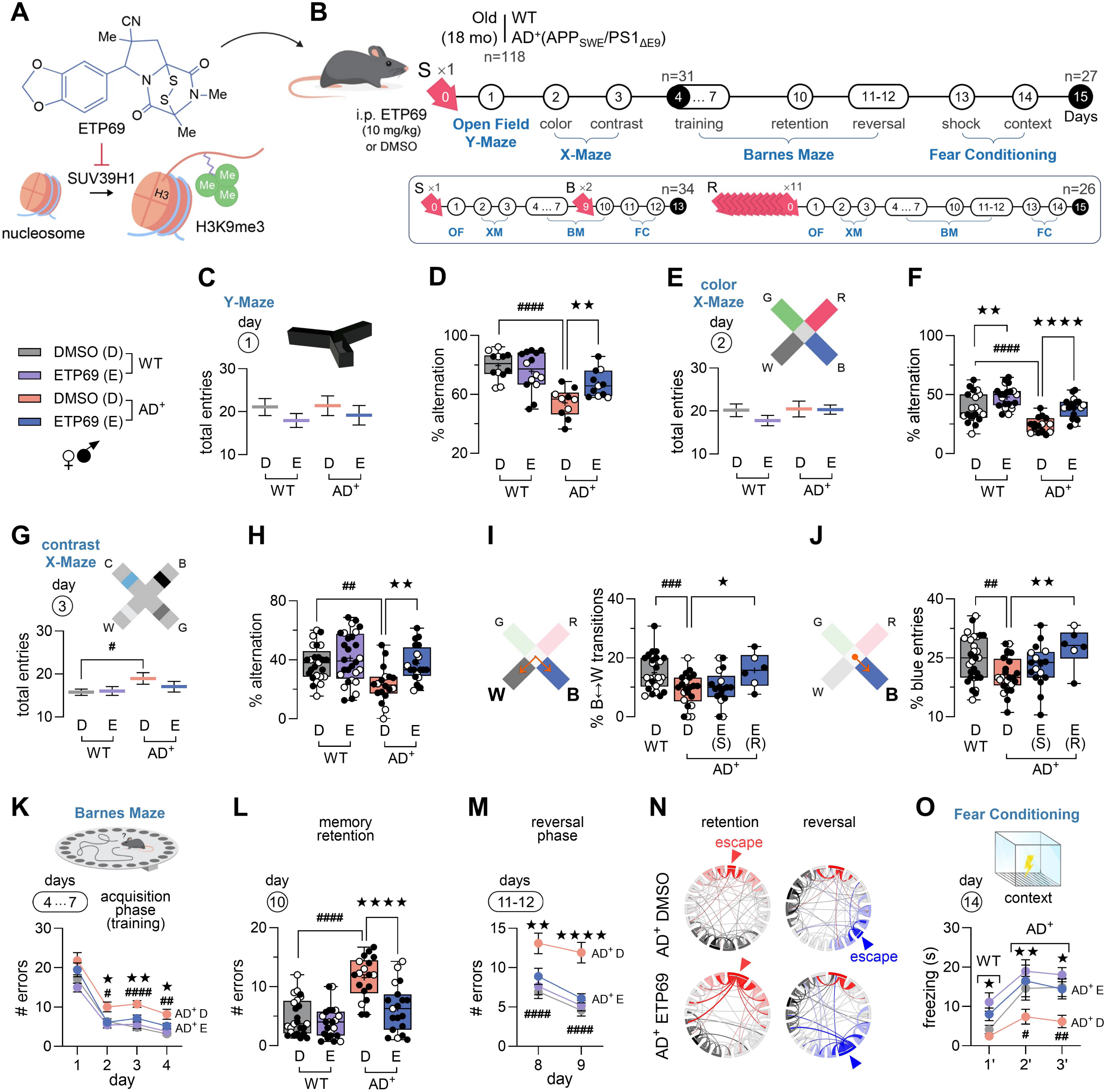
Short- and long-term effects of ETP69 on cognitive functions in old WT and AD^+^ mice. (**A**) Molecular structure of ETP69, an analogue of epidithiodiketopiperazine alkaloid chaetocin A and inhibitor of SUV39H1 responsible for trimethylation (me3) of H3K9. (**B**) Timeline of behavioural testing and regimens of i.p. ETP69 administration (10 mg/kg in 50% DMSO/saline) or 50% DMSO/saline in 18-month-old APP_SWE_/PS1_ΔE9_ (AD^+^) mice and WT littermates matched for age and sex. Two cohorts of AD^+^ and WT mice received a single ETP69 dose (regimen S) 1 day before the start of behavioural testing (day 0) and were euthanized either on day 4 or day 15. A third group of AD^+^ and WT mice received repeated ETP69 doses (once weekly for 11 weeks) with the last administration on day 0 (regimen R). A fourth group of AD^+^ and WT mice received the first dose of ETP69 on day 0 and a booster dose on day 9, the day before memory retention testing involving the Barnes maze (regimen B). Control AD^+^ and WT mice received DMSO according to the same regimens S, R, or B. Data from the 1^st^ week of testing for regimens S and B were combined since conditions were the same prior to the ETP69 boost. Regimen S: ETP69-AD^+^ mice (*n* = 18), control AD^+^ mice (*n* = 15), ETP69-WT mice (*n* = 22), and control WT mice (*n* = 21); regimen R: ETP69-AD^+^ mice (*n* = 6), control AD^+^ mice (*n* = 7), ETP69-WT mice (*n* = 6), and control WT mice (*n* = 7); and regimen B: ETP69-AD^+^ mice (*n* = 8), control AD^+^ mice (*n* = 7), ETP69-WT mice (*n* = 10), and control WT mice (*n* = 9). (**C–H**) Behavioural data for mice administered a single (S) injection. (**C**) Total number of entries for locomotor activity and (**D**) percentage of alternations for cognitive function in the Y-maze test. (**E**) Number of entries and (**F**) percentage of alternations in the color mode of the visual stimuli X-maze test. (**G**) Number of entries and (**H**) percentage of alternations in the contrast mode of the X-maze. (**I–J**) Behavioural data for mice administered a single (S) or repeated (R) injection. (**I**) Percentage of bidirectional transitions between the Blue and White arms and (**J**) percentage of entries into the Blue arm in the color mode of the X-maze test. (**K–O**) Combined data for mice subjected to regimens S, R, and B. (**K**) Error number during the acquisition phase, (**L**) on the retention test day, and (**M**) during reversal phases of the Barnes maze test, indicating memory and learning. (**N**) Chord diagrams representing average trajectory during the retention and reversal phases in ETP69-treated versus control AD^+^ mice. Arrowheads indicate the location of the escape box. (**O**) Freezing time in seconds during the contextual fear conditioning test. Circled number above each plot represents the day(s) when behavioural testing occurred according to the experimental timeline. D = DMSO-injected (control) and E = ETP69-treated animals. Individual data points are presented with group means ± SEMs. Box-and-whisker plots display median and lower and upper quartiles, together with filled and empty circles representing male and female mice, respectively. # *p* < 0.05, ## *p* < 0.01, and #### *p* < 0.0001: DMSO-AD^+^ versus DMSO-WT mice; * *p* < 0.05, ** *p* < 0.01, *** *p* < 0.001, and **** *p* < 0.0001: ETP69-mice versus DMSO-mice, by one-way or two-way ANOVA followed by Fisher’s LSD *post hoc* test.

**Fig. 3.**
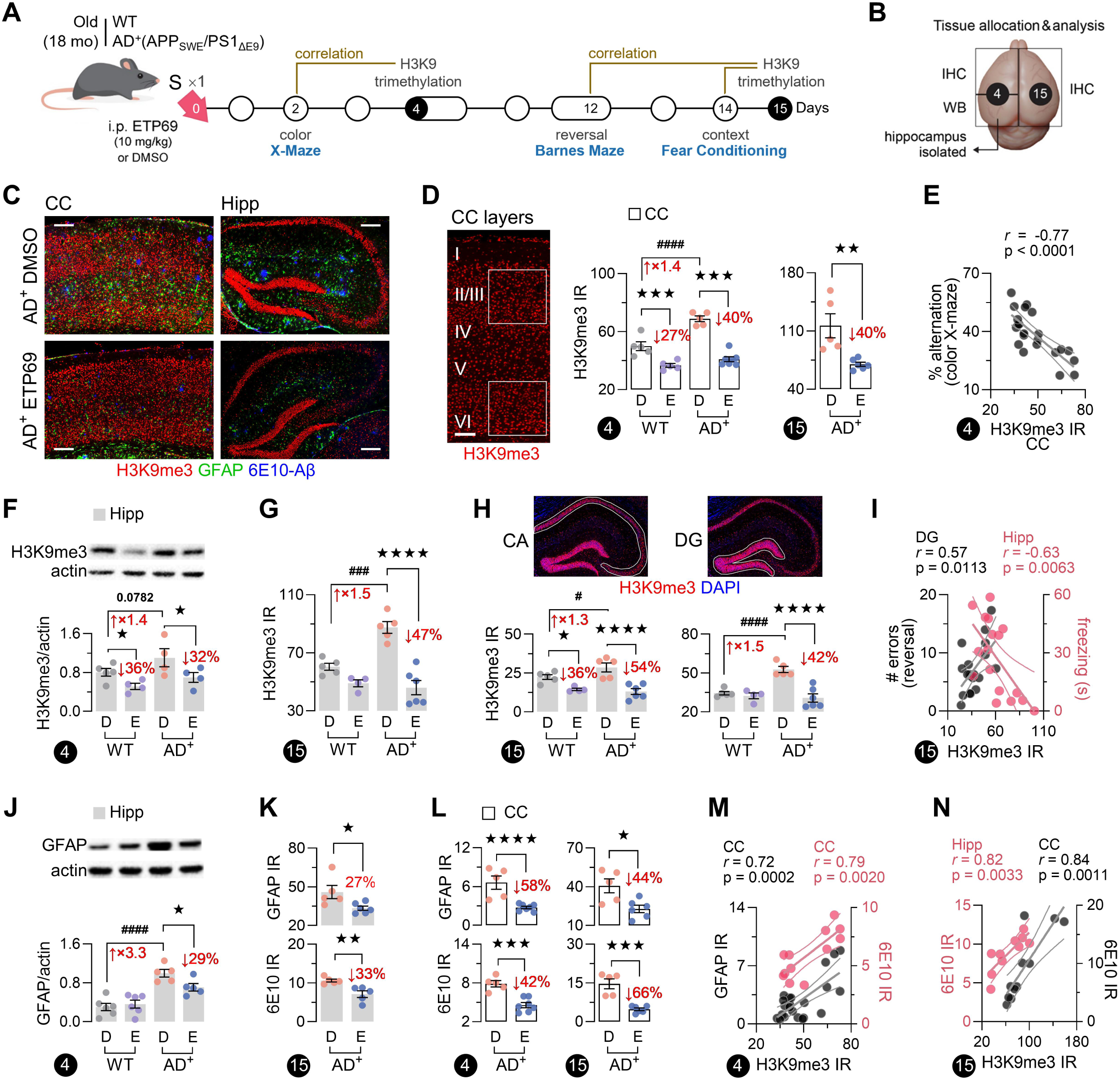
Rapid and sustained effects of a single ETP69 injection on cerebral H3K9me3 and AD-related pathology. (**A**) Simplified experimental timeline and correlations. (**B**) Brain tissue allocations for molecular analyses at the day 4 and day 15 end points. IHC, immunohistochemistry; WB, Western blotting. (**C**) Representative images of fluorescent immunostaining for H3K9me3 (red), GFAP (green, reactive astrocytes), and 6E10 (blue, Aβ plaques) in the cerebral cortices (CC) and hippocampi (Hipp) of 18-month-old APP_SWE_/PS1_ΔE9_ (AD^+^) mice administered ETP69 or DMSO (scale bar: 100 μm). (**D**) Image of H3K9me3 IR in the cortex showing areas used for measurements (layers II/III and VI) and quantification of cortical H3K9me3 IR area (μm^2^×10^3^). (**E**) Pearson’s coefficient between cortical H3K9me3 IR and performance in the color mode of the X-maze test. (**F**) Western blot analysis of H3K9me3 levels in the hippocampus. (**G**) Quantification of H3K9me3 IR area (μm^2^×10^3^) in the hippocampus. (**H**) Images of immunostaining for H3K9me3 (red) and DAPI (blue) in the hippocampus showing cornu ammonis (CA) and dentate gyrus (DG) areas used for measurements, and corresponding quantifications of H3K9me3 IR (% area). (**I**) Pearson’s correlations between hippocampal H3K9me3 IR and behavioural performance in the Barnes maze and contextual fear conditioning tests. (**J**) Western blot analysis of GFAP expression in the hippocampus. (**K–L**) Quantification of GFAP and 6E10 IR area (μm^2^×10^3^) in the (**K**) hippocampus and (**L**) cerebral cortex. (**M**) Pearson’s correlations of H3K9me3 IR with GFAP and 6E10 IR in the cerebral cortex on day 4 and (**N**) with 6E10 IR in the cortex and the hippocampus on day 15. Black circled numbers represent end-point days according to the experimental timeline. Individual data points are presented with group means ± SEMs. # *p* < 0.05, ### *p* < 0.001, and #### *p* < 0.0001: DMSO-AD^+^ mice versus DMSO-WT mice; * *p* < 0.05, ** *p* < 0.01, *** *p* < 0.001, and **** *p* < 0.0001: ETP69-mice versus DMSO-mice; by one-way ANOVA followed by Fisher’s LSD *post hoc* test or two-tailed unpaired Student’s t-test.

### Postmortem human brains

Human brain tissues were obtained from the Alzheimer’s Disease Research Center (ADRC) Neuropathology Core in the Department of Pathology (IRB protocol HS-042071) of Keck School of Medicine at the University of Southern California (USC, Los Angeles). USC-ADRC maintains human tissue collection protocols that are approved by institutional managerial committees and subject to oversight by the National Institutes of Health. Histological studies were performed under an approved IRB protocol at Cedars-Sinai Medical Center.

We examined brains from deceased patient donors with clinically and neuropathologically confirmed AD (*n*=11) or MCI (*n*=5) as well as brains from deceased individuals with normal cognition (CN [control], *n*=6). Donor information is provided in Supplementary Table S1. Fresh brain tissues (frontal cortex) were snap frozen and stored at −80°C. Portions of fresh-frozen brain tissues were fixed in 4% PFA for 16 h and then dehydrated in 30% sucrose/PBS. The brain tissues were coronally sectioned (30 μm thick) on a cryostat (Leica CM 3050_S) and mounted on slides coated with 3-aminopropyltriethoxysilane (#A3648 Sigma-Aldrich). The sections were then treated with target retrieval solution (pH 6.1; S1699, Dako) at 99°C for 40 min and washed with PBS before being used for immunohistochemistry (IHC). Additional details about the IHC analyses are provided below.

### Immunohistochemical analysis

Briefly, mouse or human brain cryosections (30 μm) were treated with serum-free protein blocking solution (X0909, Dako). The tissues were incubated with the following primary antibodies overnight at 4°C: for mouse tissues, anti-ANXA2 (1:200), anti-Aβ 1-16 clone 6E10 (1:200–1:500), anti-BDNF (1:200), anti-CD45 (1:25), anti-GFAP (1:500), anti-H3K9me3 (1:500), anti-Iba1 (1:250), anti-MT2A (1:100), anti-NeuN (1:1000), anti-PLTP (1:40), anti-PDS95 (1:6000), anti-Tau phospho-Ser396/404 (PHF-tau, 1:200) and anti-VGF (1:200); for human tissues, anti-Aβ 17– 24 clone 4G8 (1:200) and anti-H3K9me3 (1:500) (See Supplementary Table S2 for sources of these antibodies and further details). Tissue sections were then incubated with host-specific fluorophore-conjugated secondary antibodies for 1 h at room temperature and coverslipped using ProLong Gold with DAPI (Molecular Probes, Life Technologies). In some cases, slides were dipped in Thio-S solution for 1 min (to stain mature Aβ plaques) after the secondary antibody step and then washed in three baths of 70% ethanol (1 min each) before mounting with ProLong Gold DAPI. Negative controls were processed using the same protocol but without a primary antibody to assess nonspecific labelling.

### Golgi-Cox staining

Mouse brain hemispheres were processed using a Hito Golgi-Cox OptimStainTM Kit (#HTKNS1125, Hito) according to the manufacturer’s instructions. Briefly, brain samples were immersed in impregnation solution in the dark at room temperature for 2 weeks and then cryoprotected at 4°C. Next, the tissues were embedded in OCT compound, sectioned at a thickness of 100−200 μm, mounted on gelatine-coated slides (#HTHS0102, Hito) and allowed to dry overnight in a dark room. The sections were then incubated in a 20% ammonia solution, dehydrated in ascending concentrations of ethanol (50%, 75%, 95%, and 100%), cleared in xylene, and mounted with Permount mounting medium (#SP15-500, Fisher Scientific).

### Microscopy and quantification

Two to three brain tissue sections per animal were analysed, and 5 to 12 images per section were captured with the same settings and exposure time using a Carl Zeiss Axio Imager Z1 fluorescence microscope (Carl Zeiss MicroImaging, Inc.) equipped with ApoTome, AxioCam MRm, and AxioCam HRc cameras. The fluorescence signals were quantified using ImageJ software or Fiji. To assess levels of H3K9me3 in neuronal versus nonneuronal cells, the intensity of H3K9me3 IR in the nuclei (manually delimited with the polygon tool of ImageJ) of NeuN^+^ (average of 30 nuclei/animal), GFAP^+^ (average of 10 nuclei/animal), or Iba1^+^CD45^+^ (average of 20 nuclei/animal) cells was measured at high magnification. H3K9me3 IR in specific regions of the hippocampus (the cornu ammonis [CA] and dentate gyrus [DG] regions) was quantified within areas manually delimited with the polygon tool of ImageJ or Fiji.

In Golgi-Cox–stained brain sections, dendrite segments of pyramidal neurons in the CA1 region of the hippocampus were imaged using a Zeiss ApoTome microscope set at 63×. Dendritic spines were manually counted and classified as thin, mushroom, or stubby spines, as previously described.^30^. Image capture and quantification analysis were performed by different investigators.

### Western Blot analysis

Mouse brain tissues were lysed in 1× PBS/1% protease inhibitor/0.5% Triton X-100 and then centrifuged at 13500*g* at 4 °C for 20 min. Protein concentrations were determined using a BCA kit (#23227, Thermo Fisher) following the standard protocol. Equal amounts of total proteins were then separated on 4–20% Tris-glycine gels (#XP04205BOX, Invitrogen) and transferred to nitrocellulose membranes. After blocking in Tris-buffered saline with Tween 20 (TBST) (10 mmol/L Tris–HCl (pH 8.0), 150 mmol/L NaCl, and 0.1% Tween 20) containing 5% (w/v) bovine serum albumin (BSA), the membranes were incubated overnight at 4 °C with the following antigen-specific primary antibodies: anti-H3K9me3 (1:1000), anti-GFAP (1:1000), and anti−β-actin (1:1000) (See Supplementary Table S2 for sources of these antibodies). The membranes were then incubated with species-specific horseradish peroxidase−conjugated secondary antibodies (1:10000). The proteins were visualized by incubation with a chemiluminescence substrate kit (#34580, Thermo Fisher). Western blot images were collected (iBright Imaging System; Thermo Fisher), and the expression of target proteins was quantified using Image Studio Lite software version 5.2 (LI-COR Biosciences, Lincoln, NE) and normalized to the expression of β-actin.

### Complete Blood Count

Blood was collected from the vena cava while the animals were in a state of deep anaesthesia, just prior to transcardiac perfusion, and transferred to EDTA tubes (BD microtainer #365974). Blood was then processed and analyzed with the aid of a hematology analyzer according to manufacturer instructions (Horiba ABX Micros 60®).

### Preparation of brain samples for mass spectrometry

Tissues were homogenized in lysis buffer (100 mM TEA bromide [241059, Sigma], 1% SDC [D6750, Sigma], and 1× protease inhibitor cocktail set I [539131, Calbiochem]) by sonication (Qsonica Sonicator with an M-Tip probe) by performing two pulses of 15 s each at 4 Hz, with a 10-s pause between the two pulses. Insoluble materials were removed by centrifugation at 14,000 *rpm* for 15 min at 4°C. The concentration of protein in the supernatants was determined using the Bradford assay (Bio–Rad Laboratories). Solubilized proteins were reduced with 5 mM dithiothreitol, alkylated with 10 mM iodoacetamide, and digested with Lys-C (Wako, Japan) at a 1:100 enzyme-to-protein ratio at room temperature overnight and then with trypsin (Promega, Madison, WI) for at least 4 hours at 37°C. The resultant peptides were acidified with 1% trifluoroacetic acid and purified using styrene divinylbenzene reversed-phase sulfonate stage tips (Empore).

### Tandem Mass Tag (TMT) labelling

Three independent 10-plex TMT experiments were performed. Briefly, dried peptides from each sample were resuspended in 100 mM HEPES (pH 8.2), and the peptide concentration was measured using the MicroBCA protein assay kit. Sixty micrograms of peptides from each sample was subjected to TMT labelling with 0.8 mg of reagent per tube. Labelling was carried out at room temperature for 1 h with continuous vortexing. To quench any remaining TMT reagent and reverse tyrosine labelling, 8 μl of 5% hydroxylamine was added to each tube, and the samples were vortexed and incubated for 15 min at room temperature. For each of the 10-plex experiments, the ten labelled samples were combined and then dried by vacuum centrifugation.

Prior to high-pH reversed-phase fractionation, the digested and TMT-labelled peptide samples were cleaned using a reversed-phase C18 clean-up column (Sep-pak, Waters) and dried in a vacuum centrifuge. The peptide mixtures were resuspended in loading buffer (5 mM ammonia solution (pH 10.5)) and separated into a total of 96 fractions using an Agilent 1260 HPLC system equipped with a quaternary pump, a degasser, and a multi-wavelength detector (MWD) (set at wavelengths of 210, 214 and 280 nm). The peptides were separated on a 55-min linear gradient from 3% to 30% acetonitrile in 5 mM ammonia solution (pH 10.5) at a flow rate of 0.3 mL/min on an Agilent 300 Extend C18 column (3.5-μm particles, 2.1 mm ID and 150 mm long). The 96 fractions were finally consolidated into eight fractions. Each peptide fraction was dried by vacuum centrifugation, resuspended in 1% formic acid, and desalted again using styrene divinylbenzene reversed-phase sulfonate stage tips (3M-Empore).

### LC–ESI–MS/MS data acquisition

MS data were collected with an Orbitrap Lumos mass spectrometer coupled to a Proxeon NanoLC-1200 UHPLC. The 100-µm capillary column was packed with 35 cm of Accucore 150 resin (2.6 μm, 150 Å; Thermo Fisher Scientific). The scan sequence began with an MS1 spectrum (Orbitrap analysis, resolution of 60,000, 400−1600 Th, automatic gain control (AGC) target of 4 × 10^5^, maximum injection time of 50 ms). Data were acquired for 90 min per fraction. Analysis at the MS2 stage consisted of higher energy collision-induced dissociation (HCD), Orbitrap analysis with a resolution of 50,000, an AGC target of 1.25 × 10^5^, a normalized collision energy (NCE) of 37, a maximum injection time of 120 ms, and an isolation window of 0.5 Th. For data acquisition, including field asymmetric waveform ion mobility spectrometry (FAIMS), the dispersion voltage (DV) was set at 5000 V; the compensation voltages (CVs) were set at −40 V, −60 V, and −80 V; and the TopSpeed parameter was set at 1.5 s per CV.

### Database searching, peptide quantification

Spectra were converted to mzXML format using MSconvert. Databases were searched, including all entries from the Human UniProt Database (downloaded August 2019). This database was concatenated with one composed of all protein sequences for that database in the reverse order. Searches were performed using a 50-ppm precursor ion tolerance for total protein level profiling. The product ion tolerance was set to 0.2 Da. These wide mass tolerance windows were chosen to maximize sensitivity in conjunction with COMET searches and linear discriminant analysis.

TMT tags on lysine residues and peptide N-termini (+229.163 Da for TMT) as well as carbamidomethylation of cysteine residues (+57.021 Da) were set as static modifications, while oxidation of methionine residues (+15.995 Da) was set as a variable modification. Peptide-spectrum matches (PSMs) were adjusted to a false discovery rate (FDR) of 1%. PSM filtering was performed using linear discriminant analysis, and the matches were assembled further to a final protein-level FDR of 1% using the picked FDR method. An isolation purity of at least 0.7 (70%) in the MS1 isolation window was used for the samples. For each protein, the filtered peptide TMT signal-to-noise (SN) values were summed to obtain protein quantifications. To control for total protein loading within each TMT experiment, the summed protein quantities in each channel were adjusted to be equal within the experiment.

Protein levels were quantified by summing reporter ion counts across all PSMs. Reporter ion intensities were adjusted to correct for the isotopic impurities of the different TMT reagents according to manufacturer specifications. Finally, the abundance of each protein was scaled such that the summed SN ratio for that protein across all channels was equal to 100, thereby generating a relative abundance (RA) measurement. The following two criteria were applied to identify significantly differentially regulated proteins: 1) at least three measurements per group, and 2) a *p* value equal to or lower than 0.05.

### Functional network and computational analysis

Heatmaps of detectable protein hierarchies were created, and a principal component analysis (PCA) was performed by using ClustVis (https://biit.cs.ut.ee/clustvis/).^31^ Volcano plots were prepared using GraphPad Prism 9.3.1. software. Pie charts of protein classifications were created using PANTHER (http://pantherdb.org). Gene Ontology enrichment analysis was performed using the MouseMine database (www.mousemine.org). The data were analysed using Ingenuity Pathway Analysis (IPA, Qiagen; https://digitalinsights.qiagen.com). Differentially expressed proteins (according to FCs and P values) were incorporated into a canonical pathway and upstream regulator analyses and were used to generate diagrams in GraphPad Prism.

### Statistical analyses

The data were analysed using GraphPad Prism 9.3.1. software. One-way or two-way analysis of variance (ANOVA) followed by Fisher’s least significant difference (LSD) *post hoc* test was used for multiple comparisons. Comparisons between two groups were analysed using an unpaired two-tailed Student’s t-test. The threshold for statistical significance was set at 0.05. The statistical association between two or more variables was determined using Pearson’s correlation (*r*) (GraphPad Prism 9.3.1.). Pearson’s coefficient indicated the direction and strength of the linear relationship between two variables. The results are shown as the means ± standard errors of the mean (SEMs). Degrees of significance between groups are indicated as follows: \**p*<0.05, \*\**p*<0.01, ***p<0.001, and \*\*\*\**p*<0.0001.

## RESULTS

### Excessive cerebral H3K9me3 links to cognitive deficits and neuropathology in AD patients

Levels of brain H3K9me3 were investigated in patients with a clinical and post-mortem neuropathological diagnosis of AD dementia (*n*=11, mean age 86.5 ± 2.8 years, 6 females/5 males) or MCI (due to AD; *n*=5, mean age 88.6 ± 1.5 years, 3 females/2 males), and compared to age- and sex-matched cognitively normal individuals (CN; *n*=6, mean age 91.3 ± 2.8 years, 4 females/2 males; summary and individual human donor information in Table 1 and Supplementary Table S1). Histological analysis of H3K9me3 immunoreactive (IR) area corrected for nuclei count in the dorsolateral prefrontal cortex (Brodmann A9), linked to decision-making and memory and impacted in AD, revealed substantial 3.3- and 2.2-fold increases in AD and MCI patients, respectively, compared to CN controls (*p* < 0.0001 and *p* = 0.0207; Fig. 1, A and B). A significant 1.6-fold increase of cortical H3K9me3 raw IR area was also observed when comparing all AD patients, including MCI, to CN subjects (*p* = 0.0068; Supplementary Fig. S1A). Higher cortical H3K9me3/DAPI levels strongly correlated with cognitive deficits (lower MMSE and higher CDR), with the strongest correlations observed in CN and early-stage AD patients (CN-MCI cases) for CDR (*r* = 0.71, *p* = 0.0142; Fig. 1C) and MMSE (*r* = -0.83, *p* = 0.0029; Fig. 1D).

**Table 1.** Demographic and brain neuropathology data of human subjects.

|  | Diagnosis |  |  | One-Way ANOVA |  |
| --- | --- | --- | --- | --- | --- |
|  | CN | MCI | AD | F-value | P-value |
| **Subjects** | 6 | 5 | 11 | N/A | N/A |
|  | 4F/2M | 3F/2M | 6F/5M |  |  |
| **Age** | 91.3 ± 2.8 | 88.6 ± 1.5 | 86.5 ± 2.8 | 0.7611 | 0.4809 |
| **Race** | 2H/4W | 1B/4W | 3A+B/8W | N/A | N/A |
|  | 33/67% | 20/80% | 27/73% |  |  |
| **PMI** | 9.7 ± 2.5 | 6.8 ± 2.9 | 10.8 ± 1.5 | 0.8492 | 0.4434 |
| **MMSE** | 28.2 ± 1.1 | 23.5 ± 3.7 | 15.2 ± 1.8 | 11.64 | **0.0008** |
| **CDR** | 0.17 ± 0.17 | 1.50 ± 0.61 | 2.45 ± 0.25 | 12.87 | **0.0003** |
| **Braak stage** | 2.7 ± 0.7 | 3.0 ± 0.7 | 5.2 ± 0.3 | 9.134 | **0.0017** |
| **Stage I-II** | 3 | 2 | 0 |  |  |
| **Stage III-IV** | 2 | 2 | 2 |  |  |
| **Stage V-VI** | 1 | 1 | 9 |  |  |
| **ABC score** | 1.93 ± 0.29 | 1.88 ± 0.29 | 2.73 ± 0.11 | 6.281 | **0.0081** |
| **Aβ score (A9)** | 1.94 ± 0.42 | 1.90 ± 0.55 | 4.00 ± 0.49 | 6.188 | **0.0090** |
| **NFT score (A9)** | 0.08 ± 0.08 | 0.90 ± 0.55 | 2.00 ± 0.59 | 3.358 | 0.0576 |
| **Atrophy score (A9)** | 1.50 ± 0.50 | 1.60 ± 0.60 | 2.64 ± 0.53 | 1.380 | 0.2757 |

Where applicable, data are presented as mean ± SEM. CN—cognitively normal, MCI—mild cognitive impairment, AD—Alzheimer’s disease. F—female, M—male. Age at death in years. H—Hispanic, W—White, B—Black, A—Asian. PMI—post-mortem interval in hours. MMSE—mini mental state examination. CDR—clinical dementia rating: 0—normal cognition, 1—mild dementia, 2—moderate dementia, 3—severe dementia. ABC score: A—Aβ plaque score modified from Thal, B—NFT stage modified from Braak, C—Neuritic plaque score modified from CERAD. Aβ, NFT and atrophy severity scores: 0—none, 1—sparse, 3—moderate, 5—frequent. A9—Brodmann area A9, dorsolateral prefrontal cortex. N/A—not applicable.

Cortical H3K9me3/DAPI levels significantly correlated with AD progression, ABC scores and Braak stage (*r* = 0.55, *p* = 0.0079 and *r* = 0.60, *p* = 0.0032, respectively; Fig. 1, E and F). Interestingly, cerebral Aβ-plaque burden correlated with H3K9me3 levels (*r* = 0.66, *p* = 0.0012; Supplementary Fig. S1B) and NFT severity more strongly correlated with H3K9me3/DAPI (*r* = 0.75, *p* = 0.0001; Supplementary Fig. S1C). Atrophy scores correlated moderately with H3K9me3/DAPI, especially in the later stages of AD (Supplementary Fig. S1D), when neurodegeneration is more prevalent (extended correlation data of A9 H3K9me3 or H3K9me3/DAPI levels with AD-related cognitive status and neuropathological parameters are summarized in Supplementary Fig S1E heatmaps). Overall, these results suggest that an increase in cortical H3K9 hyper-trimethylation coincides with the severity of AD pathology and cognitive dysfunction.

Next, we examined epigenetic pathways identified by proteomics analysis on mass spectrometry (MS) in another human cohort,^32^ including confirmed AD patients (*n*=10, mean age 90.0 ± 4.8 years, 7 females/3 males) and age- and sex-matched cognitively normal individuals (CN; *n*=8, mean age 91.3 ± 3.6 years, 5 females/3 males). Gene ontology analysis of upregulated differentially expressed proteins (DEPs) in AD versus CN showed the enrichment of proteins involved in chromatin organization and remodelling, nucleosome assembly, and epigenetic regulation of gene expression (Fig. 1G). These proteins included many histones, especially histone H1, which functions as a linker between two nucleosome units, promoting its stabilization and the high-order packaging of chromatin in the nucleus, and histone H3-3, which accumulates in aged human brain (Supplementary Fig. S1, F and G).^33,34^ The first and second neighbour proteins upstream and downstream of SUV39H1, the histone methyltransferase responsible for the trimethylation of H3K9, was further analysed using Ingenuity Pathway Analysis (IPA) on the entire dataset of up- and down-regulated DEPs. The expression profile of all the DEPs in our dataset, with direct or indirect relationship with SUV39H1, predicted its activation in AD brains (Fig. 1H), consistent with our findings of excessive cortical H3K9me3 in MCI and AD patients.

### Targeting H3K9me3 with ETP69 rapidly improves spatial memory in old AD mice

In the following set of experiments, we explored whether reduction of H3K9me3 levels via ETP69-mediated SUV39H1 inhibition would ameliorate cognitive function in 18-month-old transgenic APP_SWE_/PS1_ΔE9_ (AD^+^ mice) and age- and sex-matched WT littermates. Mice were treated with intraperitoneal (i.p.) injections of ETP69 (10 mg/kg dissolved in dimethyl sulfoxide [DMSO]) according to different regimens, as detailed in Figure 2, A and B (*n*=118 mice). Control AD^+^ and WT mice received DMSO according to the respective regimens. All animals underwent a series of behavioural tests to assess locomotor, hippocampus-based spatial learning and memory, and visual functions (Fig. 2, and Supplementary Fig. S2 and S3), and to determine rapid and prolonged ETP69 effects as early as 1 day and up to 14 days post–ETP69 injection (Fig. 2B).

The effects of ETP69 treatment were first evaluated within days 1–3 by performing the open field, Y-maze, and color- and contrast-mode visual X-maze tests (Fig. 2, C to J, and Supplementary Fig. S2). The locomotor activity in AD^+^ and WT mice was largely unaffected by ETP69 as assessed by distance travelled, resting time, average speed, and rearing in the open field test and by arm entries in the Y-maze and X-maze tests. Notably, the short-term working memory deficits detected in old AD^+^ compared to WT mice were reversed by ETP69 treatment, as observed by the increased percentage of spontaneous alternations (Y-maze: *p* = 0.0093; color X-maze: p < 0.0001; contrast X-maze: *p* = 0.0020; Fig. 2, D, F and H; data for repeated ETP69 administration in Supplementary Fig. S2, G and I). These data indicate spatial memory restoration by ETP69 in old AD^+^ mice. A single dose of ETP69 increased spontaneous alternations in the color-mode X-maze in old WT mice (*p* = 0.0027; Fig. 2F), demonstrating that ETP69 can improve cognitive function in normal ageing.

We next measured arm transitions and entries in the color-mode X-maze test to assess the potential effect of ETP69 on visual function. Specifically, we recently found that reduced bidirectional Blue↔White transitions and Blue arm entries reflected color vision dysfunction in AD^+^ mice.^28,29^ Indeed, this study shows that AD^+^ mice made fewer Blue↔White transitions (*p* = 0.0009; Fig. 2I and Supplementary Fig. S2, J and K) and Blue arm entries (*p* = 0.0059; Fig. 2J) than WT mice. Of note, both bidirectional Blue↔White transitions and Blue arm entries were reversed in AD^+^ mice to the levels of WT mice by repeated ETP69 injections (*p* = 0.0383 and *p* = 0.0035, respectively; Fig. 2, I and J).

### ETP69 leads to prolonged cognitive improvement in old WT and AD mice

We proceeded to examine the potential lasting effects of ETP69 treatment from days 4 to 14 on additional aspects of learning and memory functions by using the Barnes maze and contextual fear conditioning tests (Fig. 2, K to O; separate data under S, R, and B regimens in Supplementary Fig. S3). In the Barnes maze test, after the spatial learning and memory acquisition phase, long-term spatial reference memory (memory retention) is evaluated after a two-day hiatus, while memory suppression and learning are assessed during the reversal phase. As expected, control AD^+^ mice made significantly more errors than WT mice prior to finding the escape box, during the 4-day training phase (RM two-way ANOVA, F(1, 30) = 27.40, p < 0.0001; Fig. 2K), the memory retention test (p < 0.0001; Fig. 2L), and the 2-day reversal phase (RM two-way ANOVA, F(1, 23) = 34.07, *p* < 0.0001; Fig. 2M). Importantly, ETP69 substantially reduced the errors made by AD^+^ mice (Training: RM two-way ANOVA, F(1, 27) = 10.23, *p* = 0.0035; Retention: p < 0.0001; Reversal: RM two-way ANOVA, F(1, 20) = 16.19, *p* = 0.0007; Fig. 2, K to M), essentially restoring the animals’ learning and memory functions comparable with non-transgenic WT mice. Chord diagrams illustrate movement frequency between paired boxes of the Barnes maze, enabling the visualization of all mice trajectories relative to the escape box location indicated by arrowheads (Fig. 2N). ETP69-treated AD^+^ mice exhibited focused search efforts near the escape box (red ribbons on memory retention day and blue ribbons on 2^nd^ day of reversal phase), as opposed to predominant random exploratory behaviour of control AD^+^ mice. In the contextual fear conditioning test, we evaluated associative learning and memory functions in the chamber (context) after aversive foot-shock stimulus. Both old WT and AD^+^ mice displayed improvement in associative memory in response to ETP69, as shown by their significantly longer freezing times (WT ETP69 vs. DMSO: *p* = 0.0119 in the 1^st^ minute; AD^+^ ETP69 vs. DMSO: RM two-way ANOVA, F(1, 26) = 11.71, *p* = 0.0021; Fig. 2O).

Altogether, performances in our array of behavioural tests indicated that even a single injection of ETP69 is sufficient to provide rapid (after 1 day) and prolonged (lasting for 14 days) improvement of hippocampus-based spatial and associative learning and memory functions in healthy and Alzheimer-like mice of advanced age.

### Reduction of cerebral H3K9me3 by ETP69 ameliorates AD-like neuropathology

Following behavioural testing, the rapid (4 days) and lasting (15 days) effects of a single injection of ETP69 (Fig. 3, A) on H3K9me3 levels and AD-like neuropathology were investigated in the brains of old AD^+^ and WT mice using Western blotting (WB) and immunohistochemistry (IHC) (Fig. 3, B to N and Supplementary Fig. S4). In control AD^+^ mice compared to WT littermates, significant 1.3–1.5-fold increases of H3K9me3 levels were detected in the densely populated II/III and VI layers of the cerebral cortex (p < 0.0001; Fig. 3, C and D) and in the hippocampus, including the cornu ammonis (CA1 to C3) and the dentate gyrus (DG) (*p* = 0.0782 - p < 0.0001; Fig. 3, F to H). ETP69 dramatically reduced H3K9me3 levels in old AD^+^ mice in the cerebral cortex (40% by IHC, p < 0.0001; Fig. 3D) and the hippocampus (32% by WB, *p* = 0.0454; Fig. 3F), 4 days post-injection, and maintained its effect up to 15 days (CC: 40% by IHC, *p* = 0.0069, and Hipp: 47%, CA: 54% and DG: 42% by IHC, p < 0.0001) (Fig. 3, D, G and H, and Supplementary Fig. S4, A and C). Notably, ETP69 treatment of old WT and AD^+^ mice reduced cortical and hippocampal H3K9me3 below the levels observed in old control WT mice (Fig. 3, D, G and H), and hypothetically to the levels of younger WT mice. H3K9me3 reduction by ETP69 in old WT ranged from 27% in the cerebral cortex (*p* = 0.0004; Fig. 3D) to 36% in the hippocampus (Hipp by WB, *p* = 0.0286 and CA by IHC, *p* = 0.0185; Fig. 3, F and H). As seen in patients with AD, our results indicated a strong association between elevated cerebral H3K9me3 and cognitive deficits in AD- model mice. The cognitive performance of mice significantly correlated with cortical H3K9me3 levels on day 4, as shown in the color mode (performed on day 2, *r* = −0.77, p < 0.0001; Fig. 3E) and contrast mode (day 3, *r* = −0.48, *p* = 0.0416; Supplementary Fig. S4B) of the X-maze test, with cortical H3K9me3 levels on day 15 in the Barnes maze test (day 10, *r* = 0.77, *p* = 0.0053; fig. S4E), and with hippocampal H3K9me3 levels on day 15 in the Barnes maze test (day 12, *r* = 0.57, *p* = 0.0113 and day 10, *r* = 0.49, *p* = 0.0299; Fig. 3I and Supplementary Fig. S4E) and the contextual fear conditioning test (day 14, *r* = −0.63, *p* = 0.0063; Fig. 3I).

We next sought to determine whether ETP69 treatment affected two features of AD neuropathology: the formation of Aβ plaques and reactive astrogliosis. In hippocampal homogenates of old control AD^+^ versus WT mice, there was a substantial increase (3.3-fold by WB, p < 0.0001) of the marker of reactive astrocytes, glial fibrillary acidic protein (GFAP), which was significantly decreased (29% by WB, *p* = 0.0223), 4 days after ETP69 treatment of AD^+^ mice (Fig. 3J). ETP69 maintained its protective effect on GFAP^+^ astrogliosis (27% by IHC, *p* = 0.0318) and reduced 6E10^+^ Aβ-plaque burden (33% by IHC, *p* = 0.0066) in the hippocampus of ETP69-treated versus control AD^+^ mice, 15 days after injection (Fig. 3K). In cortical regions of AD^+^ mice, the effects of ETP69 on astrogliosis and Aβ load were up to twice greater than in the hippocampus with significant reduction in GFAP and 6E10 IR both 4 days (GFAP: 58%, *p* < 0.0001 and 6E10: 42%, *p* = 0.0004) and 15 days (GFAP: 44%, *p* = 0.0133 and 6E10: 66%, *p* = 0.0004) post-ETP69 treatment (Fig. 3L). Further, we found that elevated cortical H3K9me3 directly correlated with increased cortical GFAP^+^ astrogliosis (day 4: *r* = 0.72, *p* = 0.0002) and Aβ plaque burden (day 4: *r* = 0.79, *p* = 0.0020 and day 15: *r* = 0.84, *p* = 0.0011) (Fig. 3, M and N). Similarly, increased hippocampal H3K9me3 strongly correlated with hippocampal Aβ accumulation (day 15: *r* = 0.82, *p* = 0.0033) (Fig. 3N). Altogether, these findings indicate that a single dose of ETP69 is sufficient to sustainably reduce cerebral H3K9me3 along with mitigating astrogliosis and Aβ plaque burden in old advanced-stage AD^+^ mice.

### ETP69 enhances cortical and hippocampal dendritic spine density in old WT and AD mice

Loss of synapses and dendritic spine abnormalities are key pathological features of AD that strongly predict cognitive decline.^35–37^ To determine whether ETP69 affects dendritic spine formation, we applied Golgi-Cox neuronal architecture staining on the cerebral cortices and hippocampi of old AD^+^ and WT mice from the B regimen cohort (Fig. 4, A to F, and Supplementary Fig. S5, A and B). Based on high-magnification photographs, we classified dendritic spines as thin (filopodia-like and long-thin), stubby, or mushroom spines according to size and shape (Fig. 4, G and H). The density (number of spines/μm of dendrite) of thin spines, representing newly formed synapses, was significantly reduced in old control AD^+^ versus WT mice, especially in the cortex (28%, *p* = 0.0004; Fig. 4I). Notably, ETP69 treatment significantly increased the density of cortical and hippocampal thin spines in WT (CC: 1.2-fold, *p* = 0.0006 and Hipp: 1.3-fold, *p* = 0.0013) and AD^+^ (CC: 1.6-fold, p < 0.0001 and Hipp: 1.5-fold, *p* = 0.0003) mice (Fig. 4I). We also observed a 2-fold increase in the density of stubby spines in the hippocampi of ETP69-treated versus DMSO-injected AD^+^ mice (*p* = 0.0461; Fig. 4J), with no significant change in the cortices. Conversely, there was a 54% decrease in the density of mushroom spines in the hippocampi, but not the cortices, of ETP69-treated AD^+^ mice (*p* = 0.0085; Supplementary Fig. S5D). Importantly, the overall number of dendritic spines was significantly increased by ETP69 treatment in the hippocampi and cortices of both old WT and AD^+^ mice (Fig. 4K). As expected, lower cortical and hippocampal thin spine density correlated with poorer performance in the Barnes maze and contextual fear conditioning tests (*r* = −0.54, *p* = 0.0301 and *r* = 0.67, *p* = 0.0063, respectively; Fig. 4, L and M).

**Fig. 4.**
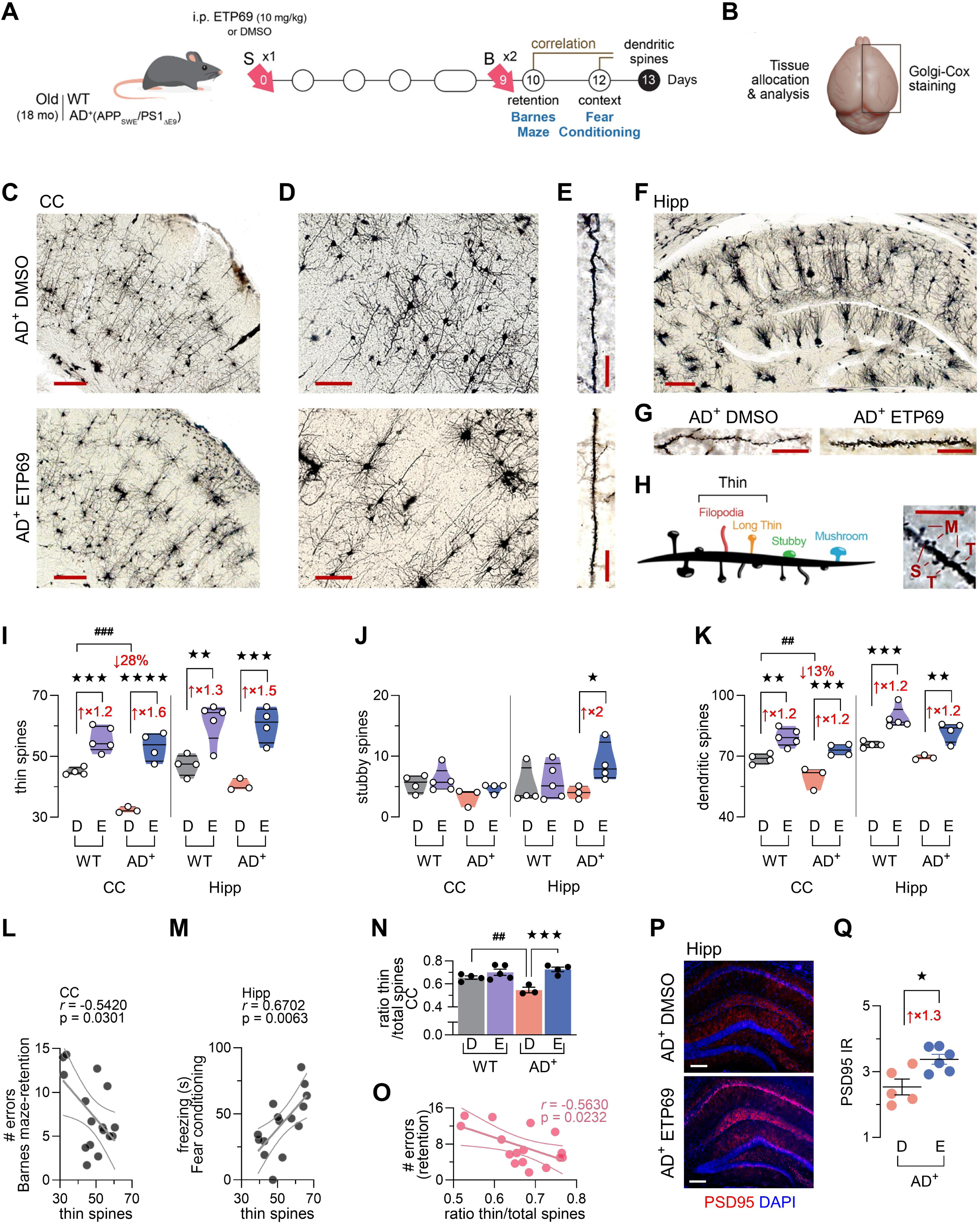
Dendritic spine formation in the cortex and hippocampus of old WT and AD^+^ mice in response to ETP69. (**A**) Simplified experimental timeline and correlations. (**B**) Brain tissue allocation for molecular analyses from 18-month-old APP_SWE_/PS1_ΔE9_ (AD^+^) and WT mice. (**C**) Low-(scale bar: 200 μm), (**D**) medium- (scale bar: 100 μm), and (**E**) high-magnification (scale bar: 20 μm) images of Golgi-Cox staining of neurons in the cerebral cortex (CC). (**F**) Low- (scale bar: 200 μm) and (**G**) high-magnification (scale bar: 20 μm) images of Golgi-Cox staining in the hippocampus (Hipp). (**H**) Illustration and high-magnification (scale bar: 5 μm) photograph of dendritic spines classified as thin (filopodia-like and long-thin; T), stubby (S), and mushroom (M) spines according to size and shape. (**I**) Density of thin, and (**J**) stubby spines as counts per 100 μm of dendrite in the cerebral cortex and hippocampus. (**K**) Density of dendritic spines as counts per 100 μm of dendrite. (**L**) Pearson’s correlation coefficients between cortical thin spine density and behavioural performance in the Barnes maze test, and (**M**) between hippocampal thin spine density and behavioural performance in the contextual fear conditioning test. (**N**) Ratio of thin spines to total spines. (**O**) Pearson’s correlations between the cortical thin spine ratio and animal’s performance in the retention phase of the Barnes maze test. (**P**) Representative images (scale bar: 200 μm) of fluorescent immunostaining for PSD95 (red, postsynaptic marker) and DAPI (blue, nuclei) and (**Q**) quantification of PSD95 IR area (μm^2^×10^3^) in the hippocampi of old AD^+^ mice injected with ETP69 or DMSO. Individual data points are presented with group means ± SEMs. Median and lower and upper quartiles are indicated on violin plots. ## *p* < 0.01 and ### *p* < 0.001: DMSO-AD^+^ mice versus DMSO-WT mice; * *p* < 0.05, ** *p* < 0.01, *** *p* < 0.001, and **** *p* < 0.0001: ETP69-mice versus DMSO-mice; by one-way ANOVA followed by Fisher’s LSD *post hoc* test or two-tailed unpaired Student’s t-test.

Thin spines accounted for the majority of all spine subtypes in old WT mice (60–70%) and, although less prevalent, AD^+^ mice (∼50%) (Fig. 4N, and Supplementary Fig. S5E). Analysis of the dendritic spine ratio showed that ETP69 treatment leads to a higher thin-to-total spine ratio and a lower mushroom-to-total spine ratio (Fig. 4N and Supplementary Fig. S5E), counteracting the effects of AD on these dendritic spine subtypes. Moreover, improved cognitive performance in the Barnes maze test was correlated with higher thin-to-total spine ratio (*r* = 0.56, *p* = 0.0232; Fig. 4O). In addition, hippocampal immunofluorescent staining analysis of the postsynaptic density protein 95 (PSD95) demonstrated a significant 1.3-fold increase in ETP69-treated versus control AD^+^ mice (*p* = 0.0137; Fig. 4, P and Q), suggesting that synaptic integrity is improved by ETP69 in old AD-model mice.

### Tau pathology, along with astrogliosis and Aβ plaque load, is mitigated by ETP69 in AD mice

In the next series of experiments, the effects of ETP69 treatment were assessed in two cohorts of 14-month-old transgenic AD^+^ mice, modelling tauopathy and amyloidosis (Fig. 5 and Supplementary Fig. S6). We first asked whether ETP69 modulates tau pathology, another major hallmark of AD. We administered a single i.p. injection of ETP69 (10 mg/kg) to 14-month-old 3xTg AD^+^ mice (Fig. 5, A and B). These mice develop hyperphosphorylated tau and NFTs by the age of 12 months.^38^ Cortical levels of H3K9me3 and PHF-tau (hyperphosphorylated tau filaments) were analysed by IHC 7 days after ETP69 injection (Fig. 5, C to E). We observed a substantial 62% reduction of H3K9me3 IR (*p* = 0.0064; Fig. 5D), as well as a significant 39% reduction of PHF-tau IR (*p* = 0.0003; Fig. 5E) in ETP69-treated compared to control 3xTg AD^+^ mice. Further, cortical H3K9me3 levels strongly correlated with PHF-tau burden (*r* = 0.78, *p* = 0.0050; Fig. 5F). These results show that ETP69 mitigates tau pathology in 3xTg AD^+^ mice.

**Fig. 5.**
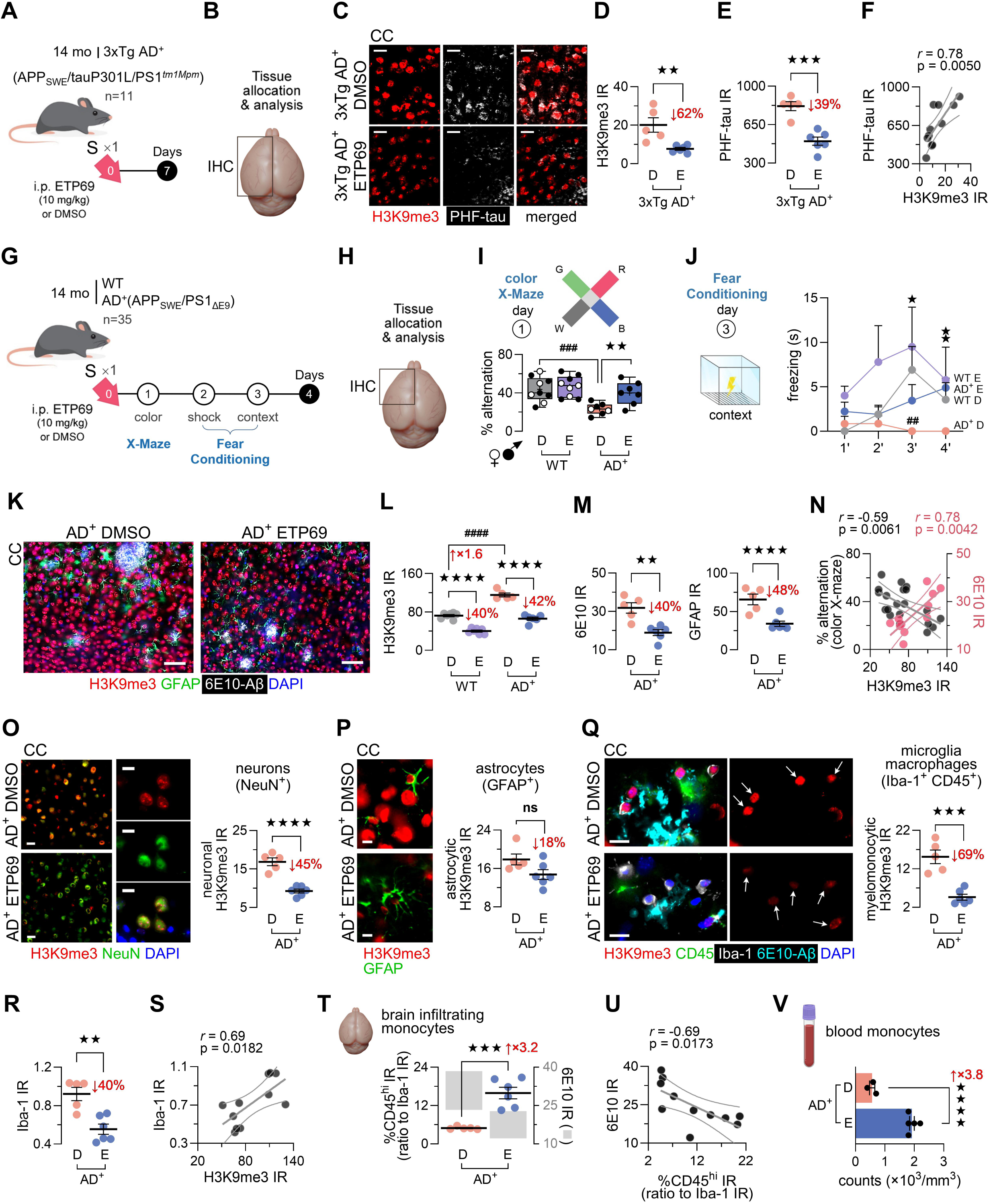
Impact of ETP69 on AD-like pathology and immune responses in WT and triple or double transgenic AD^+^ mice. (**A**) Experimental timeline involving a single (S) i.p. injection of ETP69 or DMSO in 14-month-old females 3xTg AD^+^ mice (ETP69-AD^+^ mice: *n* = 6, DMSO-AD^+^ mice: *n* = 5). (**B**) Brain tissue allocation for molecular analysis. IHC, immunohistochemistry. (**C**) Representative images of immunofluorescent staining (scale bar: 20 μm) for H3K9me3 (red) and PHF-tau (white, NFTs) in the cerebral cortex (CC). (**D**) Quantification of H3K9me3 IR area (μm^2^×100) and (**E**) PHF-tau IR area (μm^2^). (**F**) Pearson’s correlation coefficient between H3K9me3 and PHF-tau cortical levels. (**G**) Timeline of experiment involving a single (S) i.p. injection of ETP69 or DMSO in 14-month-old APP_SWE_/PS1_ΔE9_ (AD^+^) mice and WT littermates, matched for age and sex. ETP69-AD^+^ mice (n = 8), control AD^+^ mice (n = 7), ETP69-WT mice (n = 10), and control WT mice (n = 10). (**H**) Brain tissue allocation for molecular analysis. IHC, immunohistochemistry; MS, mass spectrometry. (**I**) Percentage of alternations (cognition/vision) in the color mode of the X-maze test. (**J**) Freezing time in seconds in the contextual fear conditioning test. (**K**) Representative images of immunofluorescent staining (scale bar: 50 μm) for H3K9me3 (red), 6E10 (white, Aβ plaques), and GFAP (green, reactive astrocytes) IR in the cerebral cortex. (**L**) Quantification of H3K9me3 IR area (μm^2^) and (**M**) GFAP and 6E10 IR area (μm^2^×10^3^) in the cerebral cortex. (**N**) Pearson’s correlation coefficient of cortical H3K9me3 levels with performance in the color mode of the X-maze test and with amyloid plaque burden (6E10). (**O–Q**) Representative images of immunofluorescent staining (scale bar: 20 μm) and quantification of H3K9me3 IR (red; % area) in different cell types: (**O**) NeuN^+^ (green) neurons; (**P**) GFAP^+^ (green) reactive astrocytes; (**Q**) Iba-1^+^ (white) and CD45^+^ (green) myelomonocytic cells. (**R**) Quantification of activated microglia-associated Iba-1 IR area (μm^2^) in the cerebral cortex. (**S**) Pearson’s correlation coefficients of H3K9me3 levels with Iba-1 IR. (**T**) Ratio of CD45^hi^ IR (infiltrating monocyte-derived macrophages) to Iba-1 IR (microglial population) in the vicinity of amyloid plaques. (**U**) Pearson’s correlation coefficient between the amount of infiltrating monocyte-derived macrophages (CD45^hi^ to Iba-1 ratio) and amyloid plaque burden (6E10). (**V**) Monocyte counts in the blood of 18-month-old AD^+^ mice administered a single injection of ETP69 or DMSO. D = DMSO-injected (control) and E = ETP69-treated animals. Individual data points are presented with group means ± SEMs. Median and lower and upper quartiles are indicated on box-and-whisker plots, together with filled and empty circles representing male and female mice, respectively. ## *p* < 0.01, ### *p* < 0.001, and #### *p* < 0.0001: DMSO-AD^+^ mice versus DMSO- WT mice; * *p* < 0.05, ** *p* < 0.01, *** *p* < 0.001, and **** *p* < 0.0001: ETP69-mice versus DMSO-mice; by one-way or two-way ANOVA followed by Fisher’s LSD *post hoc* test, or two- tailed unpaired Student’s t-test.

We next tested the effects of ETP69 single injection on cognition, neuropathology, and proteome profiles in 14-month-old double-transgenic AD^+^ (APP_SWE_/PS1_ΔE9_) mice and their WT littermates (Fig. 5, G and H). We found that ETP69 improves the cognitive functions of AD^+^ and WT mice in the color X-maze and fear conditioning tests (Fig. 5, I and J, and Supplementary Fig. S6, A to D). While locomotor activity, as assessed by arm entry number, was unaffected by ETP69 (Supplementary Fig. S6A), the cognitive deficits of AD^+^ versus WT mice (X-maze: *p* = 0.0009 and Fear conditioning: *p* = 0.0021) was reversed by ETP69, as shown by increased percentage of X-maze alternations (*p* = 0.0089) and fear conditioning freezing times (*p* = 0.0203) (Fig. 5, I and J, and Supplementary Fig. S6, B and D). ETP69 also improved cognition in WT mice (Supplementary Fig. S6, B and D). In AD^+^ versus WT mice, visual impairment, manifested by a decrease in B↔W bidirectional transitions (*p* = 0.0278), was unchanged by the single ETP69 dose (Supplementary Fig. S6C).

Histopathological analysis of cortical sections revealed 40–42% reductions of H3K9me3 levels in ETP69-treated WT and AD^+^ mice (p < 0.0001); levels were substantially increased in the control AD^+^ versus WT mice (1.6-fold, p < 0.0001) (Fig. 5, K and L, and Supplementary Fig. S6E). Consistent with our observations in 18-month-old mice, ETP69 mitigated AD-like pathology, as seen by reduced GFAP^+^ astrogliosis (48%, p < 0.0001) and 6E10^+^ Aβ plaque burden (40%, *p* = 0.0019) (Fig. 5M and Supplementary Fig. S6G). Pearson’s *r* correlation analyses showed that reduced cortical H3K9me3 levels were associated with improved performances in the color mode of the X-maze test (*r* = −0.59, *p* = 0.0061; Fig. 5N) and fear conditioning test (*r* = −0.59, *p* = 0.0037; Supplementary Fig. S6F), and with reduced Aβ plaque load (*r* = 0.78, *p* = 0.0042; Fig. 5N).

### ETP69 induces circulating and brain-infiltrating monocytes, while reducing microgliosis

We next assessed whether the effect of ETP69 on H3K9me3 levels was cell type specific in the brain of 14-month-old AD^+^ mice. Co-immunostaining of H3K9me3 with markers of neurons (NeuN; Fig. 5O), astrocytes (GFAP; Fig. 5P), and myelomonocytes (Iba-1^+^CD45^+^ microglia/macrophages; Fig. 5Q), showed a preferential reduction in H3K9me3 in microglia and macrophages (69%, *p* = 0.0002) as well as neurons (45%, p < 0.0001), but not in astrocytes.

These findings prompted us to determine whether ETP69 administration affected microglial activation and the inflammatory cell milieu surrounding cerebral Aβ plaques in 14-month-old AD^+^ mice. In the cerebral cortex of ETP69-treated AD^+^ mice, we found a reduction of ionized calcium-binding adaptor molecule 1 (Iba-1) IR (40%, *p* = 0.0021; Fig. 5R and Supplementary Fig. S6H), indicating the amelioration of microgliosis and neuroinflammation by ETP69. Cortical H3K9me3 was significantly associated with microgliosis (*r* = 0.69, *p* = 0.0182; Fig. 5S). In the vicinity of 6E10^+^ Aβ plaques, the percentage of Iba-1^+^CD45^high^ IR, representing infiltrating monocyte-derived macrophages, displayed a 1.8-fold increase following ETP69 treatment (*p* = 0.0006; Supplementary Fig. S6, I and J). Along with reduced Aβ plaque burden and microgliosis, a 3.2- fold increase in peripherally derived Iba-1^+^CD45^high^ macrophages relative to the total Iba-1^+^ brain-resident microglial population was observed in ETP69-treated versus control AD^+^ mice (*p* = 0.0004; Fig. 5T). These results suggest that ETP69 may promote cerebral recruitment of monocyte-derived macrophages, which are professional phagocytic cells beneficial for Aβ clearance.^27,39–45^ Indeed, we found that the enhanced area covered by brain-infiltrating CD45^high^ macrophages was significantly associated with a reduced 6E10^+^ Aβ plaque load (*r* = −0.69, *p* = 0.0173; Fig. 5U). Furthermore, evaluation of circulating blood cell counts in a subset of 18-month-old AD^+^ mice indicated that ETP69 treatment substantially induced peripheral innate immune responses, with a 3.8-fold increase in monocytes (*p* = 0.0039; Fig. 5V) and a 2.9-fold increase in granulocytes (*p* = 0.0375; Supplementary Fig. S6K), and no significant effects on lymphocytes and red blood cell numbers (Supplementary Fig. S6, L and M). Overall, these data suggest that ETP69 induction of blood monocytes leads to increased cerebral infiltration of these cells and enhanced Aβ clearance and reduced gliosis in AD^+^ mice.

### Proteome dysregulation in AD mouse brains is reversed by ETP69

We next sought to gain insights into the neuroprotective mechanisms of ETP69 by global proteomics (Figs. 6 and 7, and Supplementary Fig. S7 to S11). To this end, we applied MS analysis on the soluble fraction of brain homogenates extracted from the 14-month-old WT and AD^+^ (APP_SWE_/PS1_ΔE9_) mice cohort. We identified 5097 proteins and quantified 4358 proteins for which expression was detected in at least three animals per group (Fig. 6A and Supplementary Fig. S7, A to C, for MS data validation). A Venn diagram displays distribution and overlap of 1118 differentially expressed proteins (DEPs; p < 0.05) identified in at least one of the following three comparisons between experimental groups: DMSO-injected AD^+^ versus WT mice (henceforth referred to as AD^+^/WT; 747 DEPs), ETP69-treated versus DMSO-injected WT mice (WT ETP69/DMSO; 318 DEPs), and ETP69-treated versus DMSO-injected AD^+^ mice (AD^+^ ETP69/DMSO; 370 DEPs). Interestingly, there were relatively few overlapping DEPs between the AD^+^ ETP69/DMSO and WT ETP69/DMSO comparisons (39 DEPs), potentially reflecting different responses to ETP69 based on the H3K9 trimethylation status in healthy WT versus AD^+^ mouse brains (Fig. 6A). While the proportions of DEPs that were up- or downregulated by ETP69 were similar, when the fold change (FC) threshold was set above 1.2, there were more than twice as many up- than down-regulated DEPs due to ETP69 treatment in both the WT and AD^+^ mice (Fig. 6B).

**Fig. 6.**
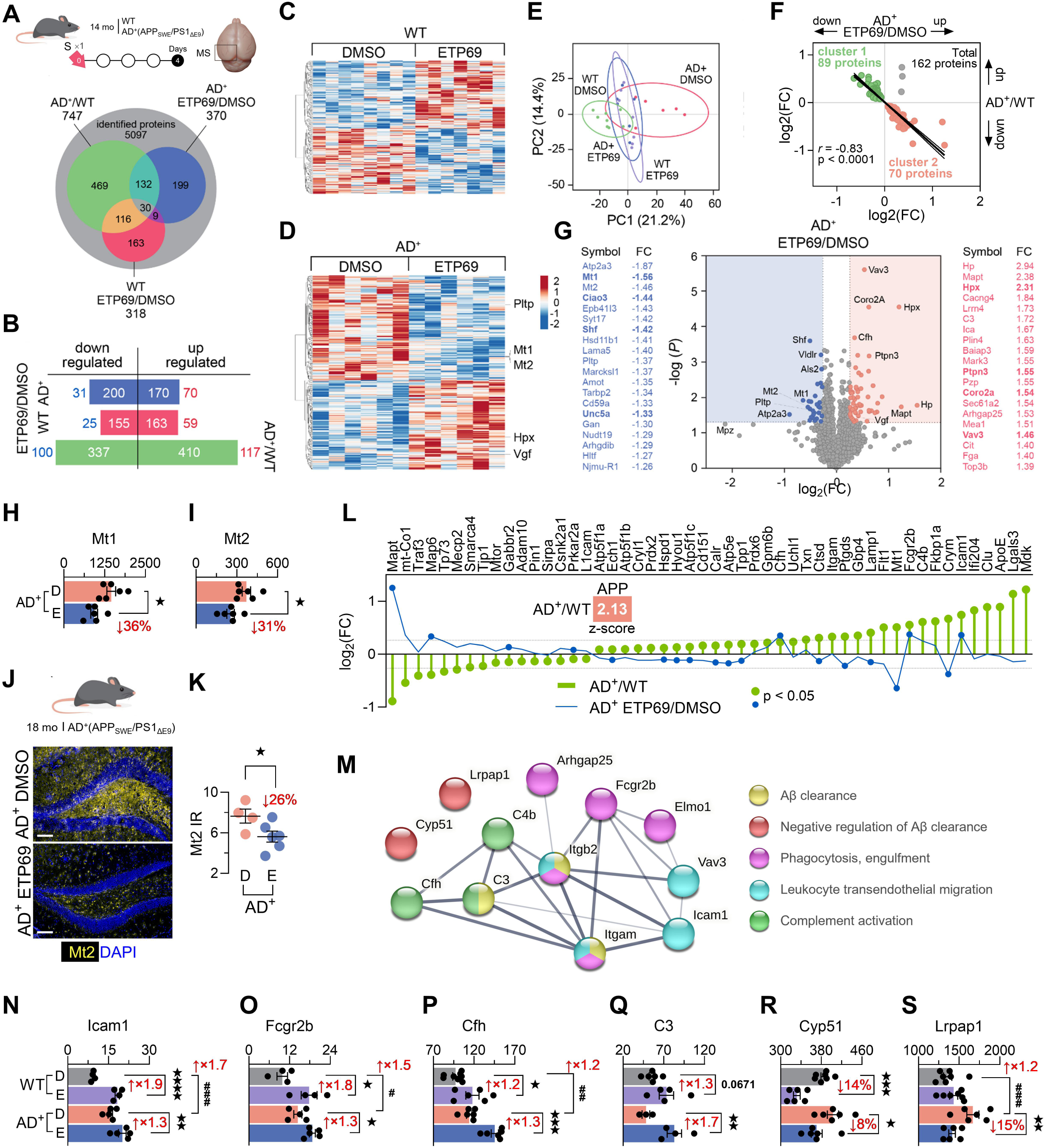
Proteome signatures of ETP69 treatment in WT and AD^+^ mice. (**A**) Venn diagram showing the overlap of all significant differentially expressed proteins (DEPs) in 14-month-old APP_SWE_/PS1_ΔE9_ (AD^+^) mice versus WT mice (*n*=6 ETP69-AD^+^ mice versus *n*=6 DMSO-AD^+^ mice, and *n*=7 ETP69-WT mice versus *n*=8 DMSO-WT mice). All mice were treated with a single (S) i.p. injection of ETP69 or DMSO. (**B**) All significantly upregulated and downregulated DEPs in each comparison. Blue and red numbers represent DEPs with |FC| above 1.2. (**C–D**) Heatmaps displaying the hierarchical clustering of all significant DEPs between the ETP69- and DMSO- groups in (**C**) WT mice and in (**D**) AD^+^ mice. Blue: downregulated proteins. Red: upregulated proteins. (**E**) Principal component analysis of protein expression profiles in AD^+^ and WT mice after ETP69 or DMSO injection. DEPs significant in at least one of the three comparisons (AD^+^ ETP69/DMSO, WT ETP69/DMSO, or AD^+^/WT) are included in the analysis (total: 1118 proteins). (**F**) Inverse relationship of the DEP fold-change (FC) for ETP69- versus DMSO- AD^+^ mice and for AD^+^ versus WT mice. The 162 significant overlapping DEPs in AD^+^ ETP69/DMSO and AD^+^/WT mice are included in this analysis. Cluster 1: 89 proteins upregulated in AD^+^ mice and downregulated after ETP69 injection; cluster 2: 70 proteins downregulated in AD^+^ mice and upregulated after ETP69 injection. (**G**) Volcano plot of all DEPs in ETP69-AD^+^ mice (versus control AD^+^ mice). (**H–I**) Quantification of (**H**) Mt1 and (**I**) Mt2 protein expression in 14-month-old mice by mass spectrometry (MS). (**J**) Representative images (scale bar: 100 μm) of fluorescent immunostaining for Mt2 (yellow) and DAPI (blue, nuclei), and (**K**) quantification of Mt2 IR area (μm^2^×10^3^) in the hippocampi of 18-month-old AD^+^ mice administered ETP69 or DMSO. (**L**) APP activation (z score = 2.13) in AD^+^ (versus WT, green bars) mice using Ingenuity Pathway Analysis (IPA). All changes in protein expression were significant (green dots). DEPs in ETP69-AD^+^ mice (versus control AD^+^ mice) are shown (blue line) with significant DEPs indicated by blue dots. (**M**) Protein association network showing select protein roles in Aβ clearance, phagocytosis, and chemotaxis using the STRING database (v11.5). Nodes (proteins) are color-coded according to function. The thickness of the edges indicates the confidence of the interaction from medium (0.4) to highest confidence (0.9). (**N–S**) Quantification of (**N**) Icam1, (**O**) Fcgr2b, complement components (**P**) Cfh, (**Q**) C3, (**R**) Cyp51, and (**S**) Lrpap1 protein expression by MS. Individual data points are presented with group means ± SEMs. # *p* < 0.05, ## *p* < 0.01, and ### *p* < 0.001: DMSO-AD^+^ mice versus DMSO-WT mice; * *p* < 0.05, ** *p* < 0.01, *** *p* < 0.001, and **** *p* < 0.0001: ETP69-mice versus DMSO-mice; by one-way ANOVA followed by Fisher’s LSD *post hoc* test or two-tailed unpaired Student’s t-test.

**Fig. 7.**
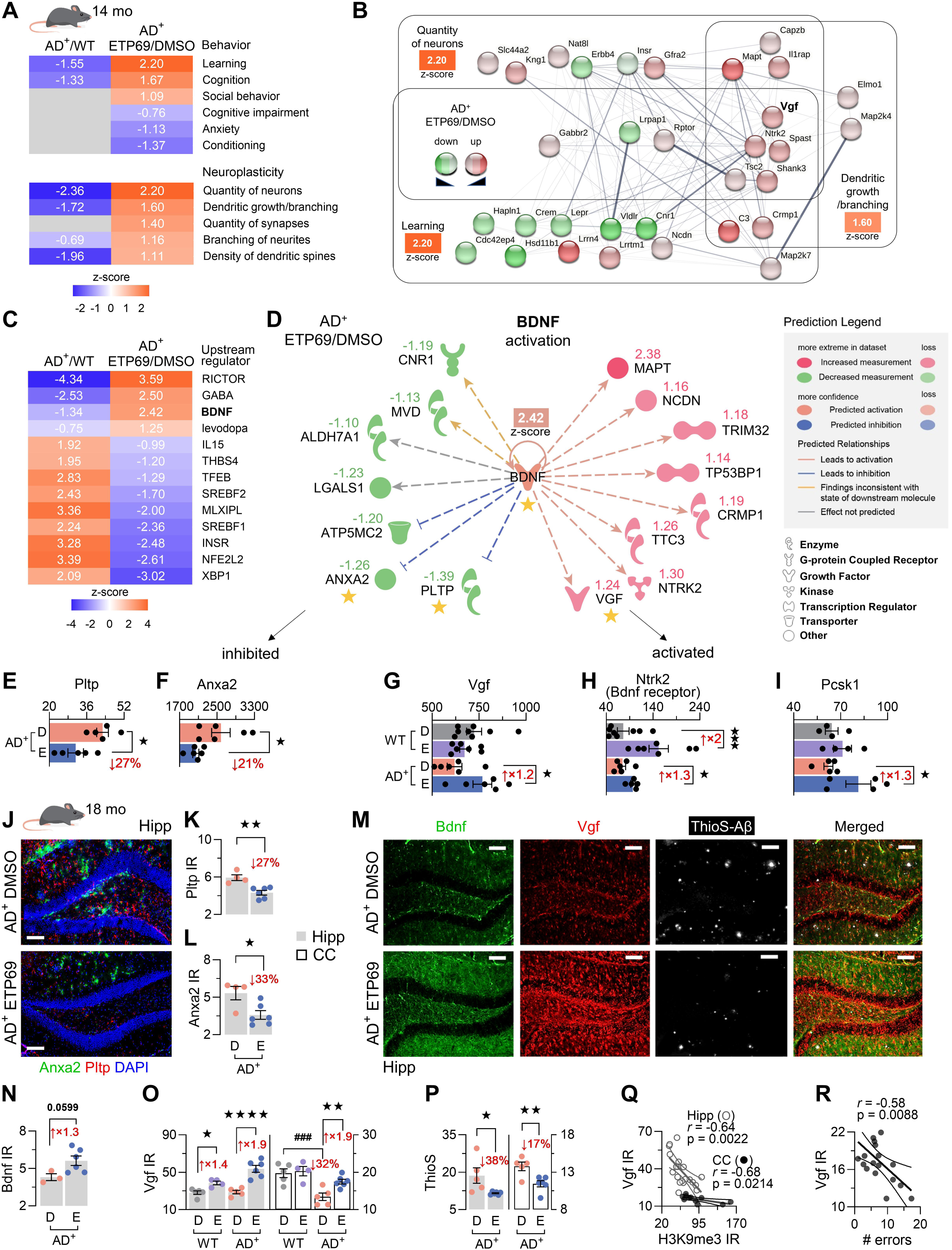
Enriched molecular pathways related to neuronal survival, synaptic plasticity, and cognition due to ETP69. (**A**) Heatmaps comparing activation z-scores for biological processes associated with behaviour and neuroplasticity determined by IPA in brain homogenates of 14- month-old APP_SWE_/PS1_ΔE9_ (AD^+^) versus WT mice and in ETP69-AD^+^ versus DMSO-AD^+^ mice, four days after a single injection of ETP69 or DMSO. (**B**) STRING v11.5 protein association network showing overlapping proteins involved in learning, quantity of neurons, and dendritic growth/branching pathways according to IPA. Red nodes: upregulated proteins; green nodes: downregulated proteins. Edge thickness ranges from a medium (0.4) to high-confidence (0.9) protein association. (**C**) Heatmaps comparing IPA activation z scores for upstream regulators in AD^+^ (versus WT) mice and in ETP69- (versus DMSO) AD^+^ mice. (**D**) IPA diagram of proteins related to the upstream regulator BDNF. The activation z score for BDNF is indicated as well as the observed FC of each target in ETP69- (versus DMSO) AD^+^ mice. (**E–L**) Quantification of (**E**) Pltp, (**F**) Anxa2, (**G**) Vgf, (**H**) Ntrk2 (TrkB), and (**I**) Pcsk1 protein expression by MS. (**J**) Representative images (scale bar: 100 μm) of fluorescent immunostaining for Pltp (red), Anxa2 (green), and DAPI (blue, nuclei), and (**K)** quantification of Pltp (μm^2^×10^3^) and (**L**) Anxa2 IR area (μm^2^×10^3^) in the hippocampi (Hipp) of 18-month-old APP_SWE_/PS1_ΔE9_ (AD^+^) mice administered ETP69 or DMSO. (**M**) Representative fluorescent images (scale bar: 100 μm) and (**N**) quantification of Bdnf IR, (**O**) Vgf IR, and (**P**) ThioS staining (μm^2^×10^3^ for each) in the hippocampi and cerebral cortices (CC) of 18-month-old mice after ETP69 or DMSO administration. (**Q–R**) Pearson’s correlation coefficients between (**Q**) hippocampal and cortical H3K9me3 and Vgf IR, and between (**R**) cortical Vgf IR and performance in the Barnes maze test (reversal phase). Individual data points are presented with group means ± SEMs. ### *p* < 0.001: DMSO-AD^+^ mice versus DMSO-WT mice; * *p* < 0.05, ** *p* < 0.01, *** *p* < 0.001, and **** *p* < 0.0001: ETP69-mice versus DMSO-mice; by one-way ANOVA followed by Fisher’s LSD *post hoc* test or two-tailed unpaired Student’s t-test.

Heatmaps and a principal component analysis (PCA) illustrate the DEP profiles that clustered distinctly for each genotype and treatment group (Fig. 6, C to E, and Supplementary Fig. S7, D to F). Strikingly, of the 162 overlapping DEPs between the AD^+^/WT and AD^+^ ETP69/DMSO comparisons, 159 DEPs that were either up- or down-regulated due to disease state (AD^+^ vs. WT) were reversed by ETP69 treatment (*r* = 0.83, *p* < 0.0001; Fig. 6F). Similarly, analysis of all DEPs with a more lenient threshold (absolute value |FC| ≥ 1.04; 2005 proteins), showed that the expression of 1683 proteins (84%) was reversed between the AD^+^/WT and AD^+^ ETP69/DMSO comparisons. Gene Ontology (GO) enrichment analysis of the 1683 DEPs whose expression was reversed, revealed protein clusters related to cytoskeleton organization and synaptic and dendritic spine function and structure that were upregulated by ETP69 as well as clusters related to metabolism, lipid oxidation, and oxidoreductase activity that were downregulated by ETP69 (Supplementary Fig. S8). Pie charts of PANTHER categories show that the most abundant biological function classes, regardless of group comparisons, were related to metabolite conversion and protein-modifying enzymes, nucleic acid metabolism, transport/trafficking, and translation (Supplementary Fig. S9A).

Volcano plots display the top 20 up- and top 20 down-regulated DEPs for ETP69-treated versus DMSO-injected AD^+^ brains (Fig. 6G) as well as for AD^+^/WT and WT ETP69/DMSO comparisons (Supplementary Fig. S9, B and C; the lists of all significant up- and down-regulated DEPs with |FC| > 1.20 for the three comparisons are in Supplementary Tables S3 to S8). Two of the top upregulated cerebral DEPs in both ETP69-treated WT and AD^+^ mice were the haemoglobin- scavenging neuroprotective proteins, haptoglobin (Hp) and hemopexin (Hpx),^46,47^ whereas Hpx was downregulated in AD^+^ compared to WT mice (Fig. 6G and Supplementary Fig. S10, A and B). In the same mouse cohort, blood Hpx and Hp levels were also substantially increased after ETP69 treatment and were strongly correlated with their respective brain levels (Supplementary Fig. S10, C to H). These results further indicate that ETP69 activated an early innate immune response. Furthermore, two of the top DEPs downregulated by ETP69 in AD^+^ mice were members of the metallothionein (Mt) family, Mt1 and Mt2, which are involved in neuronal stress and inflammation (Fig. 6, G to I).^48,49^ We also confirmed by IHC that cerebral Mt2 expression was significantly decreased in 18-month-old AD^+^ mice following ETP69 treatment (Fig. 6, J and K).

### ETP69 induces innate immune system–mediated Aβ clearance and phagocytosis

To better understand the molecular implications of AD-like pathology and ETP69 treatment, we next performed Ingenuity Pathway Analysis (IPA). As expected, IPA predicted that the amyloid precursor protein (APP) processing pathway was significantly activated in AD^+^ compared to WT mice (z-score = 2.13, *p* < 0.0001; Fig. 6L). This pathway included known markers of AD, such as molecular chaperone clusterin (Clu or apolipoprotein J), cholesterol carrier apolipoprotein E (ApoE), neurodegeneration-related cytokine midkine (Mdk), and neurodegeneration-associated microglial marker galectin-3 (Lgals3). Although expression of these proteins was unaffected by ETP69, 19 other APP-related proteins, including Mt1, disintegrin and metalloproteinase domain– containing protein 10 (Adam10 or CD156c), ketimine reductase mu-crystallin (Crym), lysosome-associated membrane glycoprotein 1 (Lamp1), microtubule-associated protein 6 (Map6), cathepsin D (Ctsd), prostaglandin-H2 D-isomerase (Ptgds), and gamma–aminobutyric acid type B receptor subunit 2 (Gabbr2), were reversed towards WT levels in the brains of AD^+^ mice after ETP69 treatment (Fig. 6L).

Our proteomics data showed that ETP69 promotes Aβ removal and related innate immune cell migration and phagocytosis markers (Fig. 6, M to S, and Supplementary Fig. S10, I to K). Specifically, ETP69 induced the expression of intercellular adhesion molecule 1 (Icam1) and guanine nucleotide exchange factor 3 (Vav3), which engage in cerebral monocyte recruitment and transendothelial migration. In addition, ETP69 upregulated low affinity immunoglobulin gamma Fc region receptor II (Fcgr2b), Rho GTPase-activating protein 25 (Ahrgap25), and engulfment and cell motility protein 1 (Elmo1), which participate in the phagocytic process. Further, complement components such as Cfh and C3 were substantially induced by ETP69, and C3 induction in the brain correlated with the elevation of C3 levels in the blood (Supplementary Fig. S10, L and M). Notably, Icam1, Fcgr2b, and Cfh, which were upregulated in AD^+^ mice, were further increased by ETP69, suggesting that ETP69 leads to a more robust neuro-immune response. Concomitant with the increase in C3, involved in Aβ clearance by microglia,^50^ there was a decrease in cytochrome P450 family 51 (Cyp51) and in low-density lipoprotein receptor-related protein-associated protein 1 (Lrpap1), two negative regulators of microglia-dependent Aβ clearance, in response to ETP69 (Fig. 6, R and S). Overall, these data support the effect of ETP69 on microglia- and monocyte-mediated phagocytosis and chemotaxis, ultimately leading to efficient cerebral Aβ removal.

### Neuronal integrity and cognition-related pathways are activated by ETP69-mediated H3K9me3 inhibition

We next assessed the effects of ETP69 on molecular pathways related to neuroprotection and cognitive function (Fig. 7). IPA revealed that ETP69 administration significantly activated pathways related to learning, cognition, quantity of neurons, as well as dendritic growth and branching, fully reversing the AD-related inhibitory effects of these pathways (Fig. 7A; Pathway diagrams in figure S11, A to D). A STRING network shows overlapping DEPs between the three functions of learning, neuronal survival, and dendritic growth/branching (Fig. 7B). Our data indicated that leucine-rich repeat neuronal protein 4 (Lrrn4), which plays an important role in hippocampus-dependent learning and long-lasting memory,^51^ was a top (1.73-fold) upregulated protein in ETP69-treated versus control AD^+^ mice (*p* = 0.015). Also, glial cell line–derived neurotrophic factor (GDNF) receptor 2 (Gfra2), which promotes neuronal survival,^52^ was a top downregulated protein in AD^+^ mice and was significantly increased in ETP69-treated mice (*p* = 0.0056). Interestingly, microtubule-associated protein tau (Mapt), a major constituent of the neuronal cytoskeleton, was also a top downregulated protein in AD^+^ versus WT mice and a top (2.4-fold) upregulated protein in response to ETP69 in AD^+^ mice (*p* = 0.0188).

The five proteins that overlapped between the three pathways included Bdnf/Nt-3 growth factor receptor (neurotrophic receptor tyrosine kinase 2; Ntrk2 gene encoding TrkB), involved in neuron survival and neuroplasticity,^53,54^ SH3 and multiple ankyrin repeat domains protein 3 (Shank3), a major scaffold postsynaptic density protein,^55^ spastin (Spast) involved in axon growth,^56^ tuberous sclerosis complex 2 (Tsc2, also known as tuberin), and the neurosecretory protein nerve growth factor-inducible Vgf, which is involved in neurogenesis and neuroplasticity (Fig. 7B).^57,58^

### Bdnf/Vgf network is activated by ETP69 and linked to improved cognitive function

To complement our analysis, we investigated the top IPA upstream regulators that were induced or inhibited by ETP69 treatment (Fig. 7C and Supplementary Fig. S11). We found that Rictor, GABA, and Bdnf were top activated networks in response to ETP69 treatment (z scores: 3.59, 2.50, 2.42, respectively), whereas these pathways were among the top inhibited pathways in AD^+^ mice. Given the key role of Bdnf in neuronal survival, synaptic plasticity, learning, and memory, we further studied the involvement of the Bdnf network after ETP69 treatment. IPA predicted activation of the Bdnf pathway in response to ETP69 in the brains of AD^+^ mice (Fig. 7, D to H), with phospholipid transfer protein (Pltp) and annexin A2 (Anxa2) downregulated, and Mapt, Vgf (activated by Bdnf), and Ntrk2 (a receptor for Bdnf) upregulated. Of note, brain Ntrk2 (TrkB) was markedly upregulated after ETP69 treatment in both WT and AD^+^ mice (Fig. 7H), suggesting a role for Bdnf in the effect of ETP69.

Interestingly, in AD^+^ mice, ETP69 administration led to increased expression of proprotein convertase subtilisin/kexin type 1 (Pcsk1; Fig. 7I), which is responsible for the cleavage of Vgf into its active peptides TLQP-62 and TLQP-21.^59^ Vgf belongs to the extended granin family, whose members regulate neuropeptide and growth factor secretion-mediating neuronal communication.^60^ Therefore, we assessed other members of the granin family in our MS data (Supplementary Fig. S11E). Chromogranin (Chg)A, ChgB (or secretogranin (Scg)1), Scg2, neuroendocrine protein 7B2 (or Scg5) were downregulated in AD^+^ mice compared to WT mice, and this was reversed by ETP69 treatment. Scg3 displayed the opposite expression pattern, while ETP69 reversed the disease effect.

We validated the MS data by performing quantitative histology of select Bdnf targets, using brain tissues from the 18-month-old mouse cohort collected 15 days after a single ETP69 or DMSO injection. Pltp and Anxa2 expression were decreased to a similar extent detected by MS, as confirmed in the hippocampi of ETP69-treated AD^+^ mice (Fig. 7, J to L). Hippocampal Bdnf expression displayed a trend of 1.3-fold increase in ETP69-treated mice (*p* = 0.0599; Fig. 7, M and N). Notably, Vgf expression was particularly strong in the subgranular zone of the dentate gyrus after ETP69 treatment (Fig. 7M). We revealed that hippocampal Vgf was significantly (1.4- to 1.9- fold) increased by ETP69 in both WT (*p* = 0.0215) and AD^+^ mice (p < 0.0001). Cortical Vgf, which was significantly decreased in AD^+^ mice (*p* = 0.0003), was increased by ETP69 treatment in the old AD^+^ mice (*p* = 0.0075; Fig. 7O). Along with these changes, ETP69 led to a decrease in ThioS-labelled Aβ plaques in the hippocampi and cortices of AD^+^ mice (Fig. 7P). Importantly, increased brain Vgf levels were significantly correlated with reduced H3K9me3 (Hipp: *r* = −0.64, *p* = 0.0022; CC: *r* = −0.68, *p* = 0.0214; Fig. 7Q) and improved cognition, as measured by a lower number of errors in the Barnes maze test (*r* = −0.58, *p* = 0.0088; Fig. 7R).

## DISCUSSION

This study reveals that targeting H3K9me3 epigenetic repression by ETP69 leads to reduced cerebral amyloidosis, tauopathy, microgliosis, and astrogliosis, alongside increased dendritic spine density and cognitive function in the double-transgenic APP_SWE_/PS1_ΔE9_ and 3xTg AD^+^ murine models. The levels of H3K9me3 were elevated in the brains of AD patients and aged AD-model mice and closely correlated with the magnitude of the cognitive deficit. A single injection of ETP69 was sufficient to ameliorate cognitive impairment as early as 1 day after administration and to last for at least 14 days. The substantial reduction in H3K9me3 in neuronal populations, and moreover, the greater reduction in myelomonocytic populations paralleled the beneficial effects of ETP69 on synaptic plasticity and innate immune responses. Proteomics analysis revealed that a significant proportion of dysregulated proteins in AD^+^ mouse brains was reversed by ETP69. Notably, ETP69 acted through multiple mechanisms to preserve cognition, including immunomodulation by increasing circulating and brain-infiltrating monocytes, mitigating Aβ- plaque burden, tau pathology, and neuroinflammation, and activating Bdnf/Vgf network, restoring neuronal and synaptic integrity. Overall, our data showed that ETP69-induced inhibition of SUV39H1 provides neuroprotection and improves cognition through immunomodulation and Vgf activation.

In recent years, evidence has emerged showing that H3K9me3 is elevated in the brain of AD patients, leading to the repression of synaptic genes.^23,61,62^ Yet, the relationship between H3K9me3 levels and cognitive decline has not been explored. In this study, H3K9me3 levels were 2-fold and 3-fold elevated in the dorsolateral prefrontal cortex of MCI (due to AD) and AD patients, respectively, and strongly correlated with CDR and MMSE scores. The connection of H3K9me3 with clinical dementia ratings and cognitive performance assessments underscores the importance of this epigenetic repression mark in the progression of AD from its earliest functional impairment stage. Further supporting epigenetic modifications and a role for H3K9me3 repression in AD, our proteomics analysis of temporal cortex samples from AD patients revealed alterations of nucleosome and chromatin organisation processes, as well as markers of epigenetic negative regulation of gene expression. Especially, Ingenuity Pathway Analysis predicted the activation of SUV39H1, the histone methyltransferase responsible for the trimethylation of H3K9.

Can targeting H3K9 epigenetic repression preserve cognition and mitigate brain pathology in the context of AD? A preliminary study in old WT animals suggested that treatment with ETP69 protects synapses and memory.^26^ Here, we reveal that ETP69 administration substantially reversed synaptic and cognitive loss as well as curbed amyloidosis, tau pathology and detrimental neuroinflammation in the double-transgenic APP_SWE_/PS1_ΔE9_ and 3xTg AD^+^ murine models. While locomotion was essentially unaffected by ETP69, restoration of color vision was indicated and there were substantial improvements in a wide range of hippocampus-based learning and memory domains. Our IPA and GO analyses revealed that ETP69 treatment, and resulting H3K9me3 reduction, led to the enrichment of proteins involved in learning, synaptic transmission, and signalling in the brains of AD^+^ mice. This agrees with a recent finding of the association between increased cerebral H3K9me3 in AD patients and downregulation of synaptic function-related genes.^23^ Dendritic spine density is closely associated with learning and memory processes, which are highly dynamic in the hippocampus and cerebral cortex.^63,64^ Our proteomics data indeed predicted positive effects of ETP69 on dendritic spine density and branching. Golgi-Cox analysis demonstrated formation of new filopodia and long-thin dendritic spines following ETP69 treatment in both WT and AD-model mice. It should be noted that in our study, mice received a 2 week-long series of behavioural tests that ended with contextual fear conditioning a day prior to dendritic spine counting. Associative learning and in particular contextual fear conditioning have been found to increase dendritic spine density and the prevalence of thin spines in the hippocampus.^65,66^ Thin spines are essential for new memory creation.^67^ Altogether, evidence of enhanced synaptic plasticity and transmission following ETP69 treatment explains, at least in part, the robust cognitive recovery observed in the AD-model mice.

Our proteomics analysis predicted Bdnf network activation in response to ETP69, which was validated by brain histology, including upregulation of Vgf, a central component of the Bdnf pathway.^58^ Previous studies have indicated that *Bdnf* transcription is influenced by epigenetic processes,^68^ including H3K9 di- and tri-methylation,^26,69,70^ and that cAMP-response element binding protein (Creb)/Bdnf/TrkB(Ntrk2) signalling is reduced in the presence of AD.^71–74^ Bdnf promotes neuronal survival, synaptic plasticity, and learning and memory functions,^75,76^ which can explain the cognitive benefits of ETP69 treatment. Notably, we identified increases in Vgf in response to ETP69 in normal ageing and old AD^+^ mouse brains that were significantly correlated with reduced cerebral H3K9me3. Indeed, according to Harmonizome analysis of the mouse brain, *Vgf* is one of 401 genes with high H3K9me3 abundance at its promoter,^77^ suggesting that Vgf is a direct target of ETP69. Beyond Vgf, our data indicate that other members of the extended granin family,^60^ including Chga, Chgb, Scg2, 7B2, and Pcsk1n, which are downregulated in AD^+^ mice, are upregulated by ETP69. Multiple omics studies have also shown downregulation of Vgf along with, albeit to a lesser extent, all other granin family members (Chga, Chgb, Scg2, Scg3, 7B2 and Pcsk1n) in AD patients.^78–92^ as well as in a murine AD model.^79,88^ ETP69 induction of the granin family, which regulates the biogenesis of vesicles and release of neuropeptides critical for neuronal activity and cognitive function,^93–96^ further supports ETP69’s neuroprotective effects on neurotransmission and synaptic function.

Of note, ETP69 upregulated Pcsk1, responsible for the proteolytic cleavage of Vgf,^97^ suggesting the presence of active Vgf-derived TLQP-62 and TLQP-21 peptides. Vgf-derived TLQP-62 promotes dendritic branching, synaptic plasticity, and long-term hippocampal memory formation.^57,58,98,99^ This is consistent with our IPA identification of ETP69-induced molecular pathways related to neuronal quantity, learning, and dendritic growth and branching. Since TLQP- 62 was shown to trigger Vgf translation via mTor signalling,^99^ requiring Raptor and Rictor,^100^ the inhibition of the Rictor pathway in AD^+^ mice and its strong activation by ETP69 suggest induced Vgf expression in an mTor-dependent manner. TLQP-62 involvement in synaptogenesis and neurogenesis ^101,102^ agrees with our finding of a marked Vgf increase in the subgranular zone of the dentate gyrus in response to ETP69. TLQP-62 is required for hippocampal memory formation and consolidation via the Bdnf receptor Ntrk2 (TrkB),^58^ which was substantially increased by ETP69 in WT and AD^+^ mice, implying that the Vgf/Pcsk1/TrkB axis is a critical component of ETP69’s effects on cognition.

This study revealed that ETP69 induces immunomodulation and mitigates AD-like neuropathology. Vgf-derived TLQP-21 peptide activates C3a receptor 1 (C3ar1) on microglia and induces microglial phagocytosis and chemotaxis, which in turn decreases Aβ plaque load and dystrophic neurites.^103–105^ Indeed, we found that markers of dystrophic neurites (e.g., Lamp1) and cerebral Aβ-plaque burden were reduced in response to ETP69, which could be attributed, at least in part, to TLQP-21. In addition, H3K9me3 was markedly decreased in the population of Iba- 1^+^/CD45^+^ microglia and macrophages surrounding Aβ plaques in ETP69-treated AD^+^ brains, suggesting that these myelomonocytic cells are particularly targeted by ETP69. Further, ETP69 substantially increased the supply of blood monocytes and recruited Iba-1^+^/CD45^hi^ monocyte– derived macrophages around cerebral Aβ lesion sites. Moreover, plasma and brain levels of haptoglobin and other mediators of early innate immune response (e.g., hemopexin, C3) were substantially increased by ETP69 in both WT and AD^+^ mice. Growing evidence shows that enhancing chemotaxis of monocytes into the brain hinders AD progression and preserves cognition by a mechanism involving macrophage-mediated phagocytosis and enzymatic degradation of Aβ plaque.^27,39–45,106–113^ Here, AD^+^ mouse brains exhibited elevated expression of key mediators of monocyte recruitment to the brain and phagocytosis such as Icam1 and its receptor Itgam/Itgb2 complex^114^ as well as Cfh and Fcgr2b. Notably, ETP69 treatment further increased levels of these immune mediators and specifically triggered C3, which binds Itgam/Itgb2,^50^ implying enhanced Aβ clearance. We postulate that these immune responses are the physiological mechanisms needed to cope with Aβ deposition and other neuropathology; in the presence of AD they are insufficient but are enhanced by ETP69. Future investigation of the effects of ETP69 on microglia and peripheral monocyte phenotypes in the context of AD and ageing is warranted.

In conclusion, we have provided evidence that H3K9me3 epigenome derepression restores cognitive function through multiple mechanisms and is a promising target for AD treatment. Indeed, ETP69 inhibition of SUV39H1, which trimethylates H3K9, successfully triggered regenerative synaptic formation and innate immunomodulation, leading to reduced AD neuropathology, including amyloidosis, tauopathy and gliosis, and improved memory and learning. Global proteome analysis revealed that ETP69 reversed a vast majority of dysregulated proteins in AD, re-balancing their expression levels to those in healthy WT mice. Overall, since a cure for AD needs to address multiple dysregulated pathways, inhibitors of epigenomic alterations appear to be an appealing new class of compounds for AD therapy.

## Data availability

The data supporting the findings of this study are available from the corresponding author upon reasonable request. The mass spectrometry proteomics data have been deposited to the ProteomeXchange Consortium via the PRIDE [1] partner repository with the dataset identifiers PXD040225 (human data) and PXD041527 (mouse data).

## Acknowledgements

The authors thank Jo Ann Eliason and Elijiah Maxfield for help in editing this manuscript. We thank the Cedars-Sinai Biobehavioral Research Core for assistance with and access to equipment for testing. We are also indebted to Drs. Carol Ann Miller and Debra Hawes (ADRC Neuropathology Core; University of Southern California) and Dr Rodrigo Medeiros (Institute of Memory Impairments; University of California, Irvine) for providing human brain tissues and neuropathological diagnoses. We thank late Prof. Peter Davies for providing the PHF-1 antibody. Finally, we thank Samuel Fuchs, Ella Maru Studio, and Biorender.com for illustrations and figure artwork.

## Author contributions

Study conception and design: MKH, DTF and KLB

Animal experiments: DTF, JPV, JS and YK

Data acquisition: DTF, AR, JPV, JS, HS, MD and JW

Mass spectrometry and data analysis: MM, VG, SG, JS, JPV and MKH

Statistical analysis: JPV, DTF, AR and MKH

Interpretation of the data: JPV, DTF, AR, JS and MKH

Drafting of the manuscript: JPV, DTF and MKH

Manuscript editing: JPV, DTF, AR and MKH

Study supervision: MKH

All authors read and approved the final manuscript.

## Supplementary Materials

Supplementary material (Supplementary Fig. S1 to S11 and Supplementary Tables S1 to S8).

## REFERENCES

1. 2023 Alzheimer’s disease facts and figures. Alzheimers Dement. Apr 2023;19(4):1598–1695. doi:10.1002/alz.13016

2. Jack CR, Jr., Bennett DA, Blennow K, et al. NIA-AA Research Framework: Toward a biological definition of Alzheimer’s disease. Alzheimers Dement. Apr 2018;14(4):535–562. doi:10.1016/j.jalz.2018.02.018

3. Heneka MT, Carson MJ, El Khoury J, et al. Neuroinflammation in Alzheimer’s disease. Lancet Neurol. Apr 2015;14(4):388–405. doi:10.1016/S1474-4422(15)70016-5

4. Palop JJ, Mucke L. Amyloid-beta-induced neuronal dysfunction in Alzheimer’s disease: from synapses toward neural networks. Nat Neurosci. Jul 2010;13(7):812–8. doi:10.1038/nn.2583

5. Carr DB, Goate A, Phil D, Morris JC. Current concepts in the pathogenesis of Alzheimer’s disease. Am J Med. Sep 22 1997;103(3A):3S–10S. doi:10.1016/s0002-9343(97)00262-3

6. Blennow K, de Leon MJ, Zetterberg H. Alzheimer’s disease. Lancet. Jul 29 2006;368(9533):387-403. doi:10.1016/S0140-6736(06)69113-7

7. Bowman GD, Poirier MG. Post-translational modifications of histones that influence nucleosome dynamics. Chem Rev. Mar 25 2015;115(6):2274–95. doi:10.1021/cr500350x

8. Kouzarides T. Chromatin modifications and their function. Cell. Feb 23 2007;128(4):693–705. doi:10.1016/j.cell.2007.02.005

9. Sultan FA, Day JJ. Epigenetic mechanisms in memory and synaptic function. Epigenomics. Apr 2011;3(2):157–81. doi:10.2217/epi.11.6

10. Herre M, Korb E. The chromatin landscape of neuronal plasticity. Curr Opin Neurobiol. Dec 2019;59:79–86. doi:10.1016/j.conb.2019.04.006

11. Singh P, Srivas S, Thakur MK. Epigenetic Regulation of Memory-Therapeutic Potential for Disorders. Curr Neuropharmacol. Nov 14 2017;15(8):1208–1221. doi:10.2174/1570159X15666170404144522

12. Campbell RR, Wood MA. How the epigenome integrates information and reshapes the synapse. Nat Rev Neurosci. Mar 2019;20(3):133–147. doi:10.1038/s41583-019-0121-9

13. Fraga MF, Esteller M. Epigenetics and aging: the targets and the marks. Trends Genet. Aug 2007;23(8):413–8. doi:10.1016/j.tig.2007.05.008

14. Berson A, Nativio R, Berger SL, Bonini NM. Epigenetic Regulation in Neurodegenerative Diseases. Trends Neurosci. Sep 2018;41(9):587–598. doi:10.1016/j.tins.2018.05.005

15. Delgado-Morales R, Agis-Balboa RC, Esteller M, Berdasco M. Epigenetic mechanisms during ageing and neurogenesis as novel therapeutic avenues in human brain disorders. Clin Epigenetics. 2017;9:67. doi:10.1186/s13148-017-0365-z

16. Kandlur A, Satyamoorthy K, Gangadharan G. Oxidative Stress in Cognitive and Epigenetic Aging: A Retrospective Glance. Front Mol Neurosci. 2020;13:41. doi:10.3389/fnmol.2020.00041

17. Nikolac Perkovic M, Videtic Paska A, Konjevod M, et al. Epigenetics of Alzheimer’s Disease. Biomolecules. Jan 30 2021;11(2) doi:10.3390/biom11020195

18. Coppede F. The potential of epigenetic therapies in neurodegenerative diseases. Front Genet. 2014;5:220. doi:10.3389/fgene.2014.00220

19. Adwan L, Zawia NH. Epigenetics: a novel therapeutic approach for the treatment of Alzheimer’s disease. Pharmacol Ther. Jul 2013;139(1):41–50. doi:10.1016/j.pharmthera.2013.03.010

20. Kouzarides T. Histone methylation in transcriptional control. Curr Opin Genet Dev. Apr 2002;12(2):198–209. doi:10.1016/s0959-437x(02)00287-3

21. Basavarajappa BS, Subbanna S. Histone Methylation Regulation in Neurodegenerative Disorders. Int J Mol Sci. Apr 28 2021;22(9)doi:10.3390/ijms22094654

22. Walker MP, LaFerla FM, Oddo SS, Brewer GJ. Reversible epigenetic histone modifications and Bdnf expression in neurons with aging and from a mouse model of Alzheimer’s disease. Age (Dordr). Jun 2013;35(3):519–31. doi:10.1007/s11357-011-9375-5

23. Lee MY, Lee J, Hyeon SJ, et al. Epigenome signatures landscaped by histone H3K9me3 are associated with the synaptic dysfunction in Alzheimer’s disease. Aging Cell. Jun 2020;19(6):e13153. doi:10.1111/acel.13153

24. Zheng Y, Liu A, Wang ZJ, et al. Inhibition of EHMT1/2 rescues synaptic and cognitive functions for Alzheimer’s disease. Brain. Mar 1 2019;142(3):787–807. doi:10.1093/brain/awy354

25. Baumann M, Dieskau AP, Loertscher BM, et al. Tricyclic Analogues of Epidithiodioxopiperazine Alkaloids with Promising In Vitro and In Vivo Antitumor Activity. Chem Sci. 2015;6:4451–4457. doi:10.1039/C5SC01536G

26. Snigdha S, Prieto GA, Petrosyan A, et al. H3K9me3 Inhibition Improves Memory, Promotes Spine Formation, and Increases BDNF Levels in the Aged Hippocampus. J Neurosci. Mar 23 2016;36(12):3611–22. doi:10.1523/JNEUROSCI.2693-15.2016

27. Koronyo Y, Salumbides BC, Sheyn J, et al. Therapeutic effects of glatiramer acetate and grafted CD115(+) monocytes in a mouse model of Alzheimer’s disease. Brain. Aug 2015;138(Pt 8):2399–422. doi:10.1093/brain/awv150

28. Vit JP, Fuchs DT, Angel A, et al. Color and contrast vision in mouse models of aging and Alzheimer’s disease using a novel visual-stimuli four-arm maze. Sci Rep. Jan 13 2021;11(1):1255. doi:10.1038/s41598-021-80988-0

29. Vit JP, Fuchs DT, Angel A, et al. Visual-stimuli Four-arm Maze test to Assess Cognition and Vision in Mice. Bio Protoc. Nov 20 2021;11(22):e4234. doi:10.21769/BioProtoc.4234

30. Risher WC, Ustunkaya T, Singh Alvarado J, Eroglu C. Rapid Golgi analysis method for efficient and unbiased classification of dendritic spines. PLoS One. 2014;9(9):e107591. doi:10.1371/journal.pone.0107591

31. Metsalu T, Vilo J. ClustVis: a web tool for visualizing clustering of multivariate data using Principal Component Analysis and heatmap. Nucleic Acids Res. Jul 1 2015;43(W1):W566–70. doi:10.1093/nar/gkv468

32. Koronyo Y, Rentsendorj A, Mirzaei N, et al. Retinal pathological features and proteome signatures of Alzheimer’s disease. Acta Neuropathol. Apr 2023;145(4):409–438. doi:10.1007/s00401-023-02548-2

33. Fyodorov DV, Zhou BR, Skoultchi AI, Bai Y. Emerging roles of linker histones in regulating chromatin structure and function. Nat Rev Mol Cell Biol. Mar 2018;19(3):192–206. doi:10.1038/nrm.2017.94

34. Tvardovskiy A, Schwammle V, Kempf SJ, Rogowska-Wrzesinska A, Jensen ON. Accumulation of histone variant H3.3 with age is associated with profound changes in the histone methylation landscape. Nucleic Acids Res. Sep 19 2017;45(16):9272–9289. doi:10.1093/nar/gkx696

35. Chapman PF, White GL, Jones MW, et al. Impaired synaptic plasticity and learning in aged amyloid precursor protein transgenic mice. Nat Neurosci. Mar 1999;2(3):271–6. doi:10.1038/6374

36. Butterfield DA. Phosphoproteomics of Alzheimer disease brain: Insights into altered brain protein regulation of critical neuronal functions and their contributions to subsequent cognitive loss. Biochim Biophys Acta Mol Basis Dis. Aug 1 2019;1865(8):2031–2039. doi:10.1016/j.bbadis.2018.08.035

37. Cardozo PL, de Lima IBQ, Maciel EMA, Silva NC, Dobransky T, Ribeiro FM. Synaptic Elimination in Neurological Disorders. Curr Neuropharmacol. 2019;17(11):1071–1095. doi:10.2174/1570159X17666190603170511

38. Oddo S, Caccamo A, Shepherd JD, et al. Triple-transgenic model of Alzheimer’s disease with plaques and tangles: intracellular Abeta and synaptic dysfunction. Neuron. Jul 31 2003;39(3):409–21. doi:10.1016/s0896-6273(03)00434-3

39. Butovsky O, Koronyo-Hamaoui M, Kunis G, et al. Glatiramer acetate fights against Alzheimer’s disease by inducing dendritic-like microglia expressing insulin-like growth factor 1. Proc Natl Acad Sci U S A. Aug 1 2006;103(31):11784–9. doi:10.1073/pnas.0604681103

40. Doustar J, Rentsendorj A, Torbati T, et al. Parallels between retinal and brain pathology and response to immunotherapy in old, late-stage Alzheimer’s disease mouse models. Aging Cell. Oct 14 2020;19(11):e13246. doi:10.1111/acel.13246

41. Kasindi A, Fuchs DT, Koronyo Y, Rentsendorj A, Black KL, Koronyo-Hamaoui M. Glatiramer Acetate Immunomodulation: Evidence of Neuroprotection and Cognitive Preservation. Cells. May 7 2022;11(9)doi:10.3390/cells11091578

42. Koronyo-Hamaoui M, Ko MK, Koronyo Y, et al. Attenuation of AD-like neuropathology by harnessing peripheral immune cells: local elevation of IL-10 and MMP-9. J Neurochem. Dec 2009;111(6):1409–24. doi:10.1111/j.1471-4159.2009.06402.x

43. Bernstein KE, Koronyo Y, Salumbides BC, et al. Angiotensin-converting enzyme overexpression in myelomonocytes prevents Alzheimer’s-like cognitive decline. J Clin Invest. Mar 2014;124(3):1000–12. doi:10.1172/JCI66541

44. Koronyo-Hamaoui M, Sheyn J, Hayden EY, et al. Peripherally derived angiotensin converting enzyme-enhanced macrophages alleviate Alzheimer-related disease. Brain. Jan 1 2020;143(1):336–358. doi:10.1093/brain/awz364

45. Li S, Hayden EY, Garcia VJ, et al. Activated Bone Marrow-Derived Macrophages Eradicate Alzheimer’s-Related Abeta(42) Oligomers and Protect Synapses. Front Immunol. 2020;11:49. doi:10.3389/fimmu.2020.00049

46. Zhao X, Song S, Sun G, et al. Neuroprotective role of haptoglobin after intracerebral hemorrhage. J Neurosci. Dec 16 2009;29(50):15819–27. doi:10.1523/JNEUROSCI.3776-09.2009

47. Hahl P, Davis T, Washburn C, Rogers JT, Smith A. Mechanisms of neuroprotection by hemopexin: modeling the control of heme and iron homeostasis in brain neurons in inflammatory states. J Neurochem. Apr 2013;125(1):89–101. doi:10.1111/jnc.12165

48. Dostie KE, Thees AV, Lynes MA. Metallothionein: A Novel Therapeutic Target for Treatment of Inflammatory Bowel Disease. Curr Pharm Des. 2018;24(27):3155–3161. doi:10.2174/1381612824666180717110236

49. West AK, Hidalgo J, Eddins D, Levin ED, Aschner M. Metallothionein in the central nervous system: Roles in protection, regeneration and cognition. Neurotoxicology. May 2008;29(3):489–503. doi:10.1016/j.neuro.2007.12.006

50. Maier M, Peng Y, Jiang L, Seabrook TJ, Carroll MC, Lemere CA. Complement C3 deficiency leads to accelerated amyloid beta plaque deposition and neurodegeneration and modulation of the microglia/macrophage phenotype in amyloid precursor protein transgenic mice. J Neurosci. Jun 18 2008;28(25):6333–41. doi:10.1523/JNEUROSCI.0829-08.2008

51. Bando T, Sekine K, Kobayashi S, et al. Neuronal leucine-rich repeat protein 4 functions in hippocampus-dependent long-lasting memory. Mol Cell Biol. May 2005;25(10):4166–75. doi:10.1128/MCB.25.10.4166-4175.2005

52. Soler RM, Dolcet X, Encinas M, Egea J, Bayascas JR, Comella JX. Receptors of the glial cell line-derived neurotrophic factor family of neurotrophic factors signal cell survival through the phosphatidylinositol 3-kinase pathway in spinal cord motoneurons. J Neurosci. Nov 1 1999;19(21):9160–9. doi:10.1523/JNEUROSCI.19-21-09160.1999

53. Chen AI, Nguyen CN, Copenhagen DR, et al. TrkB (tropomyosin-related kinase B) controls the assembly and maintenance of GABAergic synapses in the cerebellar cortex. J Neurosci. Feb 23 2011;31(8):2769–80. doi:10.1523/JNEUROSCI.4991-10.2011

54. Glerup S, Bolcho U, Molgaard S, et al. SorCS2 is required for BDNF-dependent plasticity in the hippocampus. Mol Psychiatry. Dec 2016;21(12):1740–1751. doi:10.1038/mp.2016.108

55. Wang X, McCoy PA, Rodriguiz RM, et al. Synaptic dysfunction and abnormal behaviors in mice lacking major isoforms of Shank3. Hum Mol Genet. Aug 1 2011;20(15):3093–108. doi:10.1093/hmg/ddr212

56. Riano E, Martignoni M, Mancuso G, et al. Pleiotropic effects of spastin on neurite growth depending on expression levels. J Neurochem. Mar 2009;108(5):1277–88. doi:10.1111/j.1471-4159.2009.05875.x

57. Alder J, Thakker-Varia S, Bangasser DA, et al. Brain-derived neurotrophic factor-induced gene expression reveals novel actions of VGF in hippocampal synaptic plasticity. J Neurosci. Nov 26 2003;23(34):10800–8.

58. Lin WJ, Jiang C, Sadahiro M, et al. VGF and Its C-Terminal Peptide TLQP-62 Regulate Memory Formation in Hippocampus via a BDNF-TrkB-Dependent Mechanism. J Neurosci. Jul 15 2015;35(28):10343–56. doi:10.1523/JNEUROSCI.0584-15.2015

59. Quinn JP, Kandigian SE, Trombetta BA, Arnold SE, Carlyle BC. VGF as a biomarker and therapeutic target in neurodegenerative and psychiatric diseases. Brain Commun. 2021;3(4):fcab261. doi:10.1093/braincomms/fcab261

60. Bartolomucci A, Possenti R, Mahata SK, Fischer-Colbrie R, Loh YP, Salton SR. The extended granin family: structure, function, and biomedical implications. Endocr Rev. Dec 2011;32(6):755–97. doi:10.1210/er.2010-0027

61. Alves VC, Figueiro-Silva J, Ferrer I, Carro E. Epigenetic silencing of OR and TAS2R genes expression in human orbitofrontal cortex at early stages of sporadic Alzheimer’s disease. Cell Mol Life Sci. Jul 5 2023;80(8):196. doi:10.1007/s00018-023-04845-1

62. Gil L, Chi-Ahumada E, Nino SA, et al. Pathological Nuclear Hallmarks in Dentate Granule Cells of Alzheimer’s Patients: A Biphasic Regulation of Neurogenesis. Int J Mol Sci. Oct 25 2022;23(21)doi:10.3390/ijms232112873

63. Bloss EB, Janssen WG, Ohm DT, et al. Evidence for reduced experience-dependent dendritic spine plasticity in the aging prefrontal cortex. J Neurosci. May 25 2011;31(21):7831–9. doi:10.1523/JNEUROSCI.0839-11.2011

64. Mahmmoud RR, Sase S, Aher YD, et al. Spatial and Working Memory Is Linked to Spine Density and Mushroom Spines. PLoS One. 2015;10(10):e0139739. doi:10.1371/journal.pone.0139739

65. Sase S, Sase A, Sialana FJ, Jr., et al. Individual phases of contextual fear conditioning differentially modulate dorsal and ventral hippocampal GluA1-3, GluN1-containing receptor complexes and subunits. Hippocampus. Dec 2015;25(12):1501–16. doi:10.1002/hipo.22470

66. Petsophonsakul P, Richetin K, Andraini T, Roybon L, Rampon C. Memory formation orchestrates the wiring of adult-born hippocampal neurons into brain circuits. Brain Struct Funct. Aug 2017;222(6):2585–2601. doi:10.1007/s00429-016-1359-x

67. Xu B, Sun A, He Y, et al. Loss of thin spines and small synapses contributes to defective hippocampal function in aged mice. Neurobiol Aging. Nov 2018;71:91–104. doi:10.1016/j.neurobiolaging.2018.07.010

68. Karpova NN. Role of BDNF epigenetics in activity-dependent neuronal plasticity. Neuropharmacology. Jan 2014;76 Pt C:709–18. doi:10.1016/j.neuropharm.2013.04.002

69. Gupta-Agarwal S, Franklin AV, Deramus T, et al. G9a/GLP histone lysine dimethyltransferase complex activity in the hippocampus and the entorhinal cortex is required for gene activation and silencing during memory consolidation. J Neurosci. Apr 18 2012;32(16):5440–53. doi:10.1523/JNEUROSCI.0147-12.2012

70. Ionescu-Tucker A, Butler CW, Berchtold NC, Matheos DP, Wood MA, Cotman CW. Exercise Reduces H3K9me3 and Regulates Brain Derived Neurotrophic Factor and GABRA2 in an Age Dependent Manner. Front Aging Neurosci. 2021;13:798297. doi:10.3389/fnagi.2021.798297

71. Amidfar M, de Oliveira J, Kucharska E, Budni J, Kim YK. The role of CREB and BDNF in neurobiology and treatment of Alzheimer’s disease. Life Sci. Sep 15 2020;257:118020. doi:10.1016/j.lfs.2020.118020

72. Jiao SS, Shen LL, Zhu C, et al. Brain-derived neurotrophic factor protects against tau-related neurodegeneration of Alzheimer’s disease. Transl Psychiatry. Oct 4 2016;6(10):e907. doi:10.1038/tp.2016.186

73. O’Bryant SE, Hobson V, Hall JR, et al. Brain-derived neurotrophic factor levels in Alzheimer’s disease. J Alzheimers Dis. 2009;17(2):337–41. doi:10.3233/JAD-2009-1051

74. Song JH, Yu JT, Tan L. Brain-Derived Neurotrophic Factor in Alzheimer’s Disease: Risk, Mechanisms, and Therapy. Mol Neurobiol. Dec 2015;52(3):1477–1493. doi:10.1007/s12035-014-8958-4

75. Wang CS, Kavalali ET, Monteggia LM. BDNF signaling in context: From synaptic regulation to psychiatric disorders. Cell. Jan 6 2022;185(1):62–76. doi:10.1016/j.cell.2021.12.003

76. Kowianski P, Lietzau G, Czuba E, Waskow M, Steliga A, Morys J. BDNF: A Key Factor with Multipotent Impact on Brain Signaling and Synaptic Plasticity. Cell Mol Neurobiol. Apr 2018;38(3):579–593. doi:10.1007/s10571-017-0510-4

77. Rouillard AD, Gundersen GW, Fernandez NF, et al. The harmonizome: a collection of processed datasets gathered to serve and mine knowledge about genes and proteins. Database (Oxford). 2016;2016 doi:10.1093/database/baw100

78. Tasaki S, Gaiteri C, Mostafavi S, De Jager PL, Bennett DA. The Molecular and Neuropathological Consequences of Genetic Risk for Alzheimer’s Dementia. Front Neurosci. 2018;12:699. doi:10.3389/fnins.2018.00699

79. Bai B, Wang X, Li Y, et al. Deep Multilayer Brain Proteomics Identifies Molecular Networks in Alzheimer’s Disease Progression. Neuron. Mar 18 2020;105(6):975–991 e7. doi:10.1016/j.neuron.2019.12.015

80. Carrette O, Demalte I, Scherl A, et al. A panel of cerebrospinal fluid potential biomarkers for the diagnosis of Alzheimer’s disease. Proteomics. Aug 2003;3(8):1486–94. doi:10.1002/pmic.200300470

81. Khoonsari PE, Shevchenko G, Herman S, et al. Improved Differential Diagnosis of Alzheimer’s Disease by Integrating ELISA and Mass Spectrometry-Based Cerebrospinal Fluid Biomarkers. J Alzheimers Dis. 2019;67(2):639–651. doi:10.3233/JAD-180855

82. Pedrero-Prieto CM, Garcia-Carpintero S, Frontinan-Rubio J, et al. A comprehensive systematic review of CSF proteins and peptides that define Alzheimer’s disease. Clin Proteomics. 2020;17:21. doi:10.1186/s12014-020-09276-9

83. Holtta M, Minthon L, Hansson O, et al. An integrated workflow for multiplex CSF proteomics and peptidomics-identification of candidate cerebrospinal fluid biomarkers of Alzheimer’s disease. J Proteome Res. Feb 6 2015;14(2):654–63. doi:10.1021/pr501076j

84. Hendrickson RC, Lee AY, Song Q, et al. High Resolution Discovery Proteomics Reveals Candidate Disease Progression Markers of Alzheimer’s Disease in Human Cerebrospinal Fluid. PLoS One. 2015;10(8):e0135365. doi:10.1371/journal.pone.0135365

85. Llano DA, Bundela S, Mudar RA, Devanarayan V, Alzheimer’s Disease Neuroimaging I. A multivariate predictive modeling approach reveals a novel CSF peptide signature for both Alzheimer’s Disease state classification and for predicting future disease progression. PLoS One. 2017;12(8):e0182098. doi:10.1371/journal.pone.0182098

86. Duits FH, Brinkmalm G, Teunissen CE, et al. Synaptic proteins in CSF as potential novel biomarkers for prognosis in prodromal Alzheimer’s disease. Alzheimers Res Ther. Jan 15 2018;10(1):5. doi:10.1186/s13195-017-0335-x

87. Sathe G, Na CH, Renuse S, et al. Quantitative Proteomic Profiling of Cerebrospinal Fluid to Identify Candidate Biomarkers for Alzheimer’s Disease. Proteomics Clin Appl. Jul 2019;13(4):e1800105. doi:10.1002/prca.201800105

88. Beckmann ND, Lin WJ, Wang M, et al. Multiscale causal networks identify VGF as a key regulator of Alzheimer’s disease. Nat Commun. Aug 7 2020;11(1):3942. doi:10.1038/s41467-020-17405-z

89. Spellman DS, Wildsmith KR, Honigberg LA, et al. Development and evaluation of a multiplexed mass spectrometry based assay for measuring candidate peptide biomarkers in Alzheimer’s Disease Neuroimaging Initiative (ADNI) CSF. Proteomics Clin Appl. Aug 2015;9(7- 8):715–31. doi:10.1002/prca.201400178

90. Jahn H, Wittke S, Zurbig P, et al. Peptide fingerprinting of Alzheimer’s disease in cerebrospinal fluid: identification and prospective evaluation of new synaptic biomarkers. PLoS One. 2011;6(10):e26540. doi:10.1371/journal.pone.0026540

91. Wang X, Allen M, Li S, et al. Deciphering cellular transcriptional alterations in Alzheimer’s disease brains. Mol Neurodegener. Jul 13 2020;15(1):38. doi:10.1186/s13024-020-00392-6

92. Park SA, Jung JM, Park JS, et al. SWATH-MS analysis of cerebrospinal fluid to generate a robust battery of biomarkers for Alzheimer’s disease. Sci Rep. May 4 2020;10(1):7423. doi:10.1038/s41598-020-64461-y

93. Fargali S, Garcia AL, Sadahiro M, et al. The granin VGF promotes genesis of secretory vesicles, and regulates circulating catecholamine levels and blood pressure. FASEB J. May 2014;28(5):2120–33. doi:10.1096/fj.13-239509

94. Gondre-Lewis MC, Park JJ, Loh YP. Cellular mechanisms for the biogenesis and transport of synaptic and dense-core vesicles. Int Rev Cell Mol Biol. 2012;299:27–115. doi:10.1016/B978-0-12-394310-1.00002-3

95. Hariri AR, Goldberg TE, Mattay VS, et al. Brain-derived neurotrophic factor val66met polymorphism affects human memory-related hippocampal activity and predicts memory performance. J Neurosci. Jul 30 2003;23(17):6690–4. doi:10.1523/JNEUROSCI.23-17-06690.2003

96. Egan MF, Kojima M, Callicott JH, et al. The BDNF val66met polymorphism affects activity-dependent secretion of BDNF and human memory and hippocampal function. Cell. Jan 24 2003;112(2):257–69. doi:10.1016/s0092-8674(03)00035-7

97. Trani E, Giorgi A, Canu N, et al. Isolation and characterization of VGF peptides in rat brain. Role of PC1/3 and PC2 in the maturation of VGF precursor. J Neurochem. May 2002;81(3):565–74. doi:10.1046/j.1471-4159.2002.00842.x

98. Li C, Li M, Yu H, et al. Neuropeptide VGF C-Terminal Peptide TLQP-62 Alleviates Lipopolysaccharide-Induced Memory Deficits and Anxiety-like and Depression-like Behaviors in Mice: The Role of BDNF/TrkB Signaling. ACS Chem Neurosci. Sep 20 2017;8(9):2005–2018. doi:10.1021/acschemneuro.7b00154

99. Lin WJ, Zhao Y, Li Z, et al. An increase in VGF expression through a rapid, transcription-independent, autofeedback mechanism improves cognitive function. Transl Psychiatry. Jul 8 2021;11(1):383. doi:10.1038/s41398-021-01489-2

100. Takei N, Nawa H. mTOR signaling and its roles in normal and abnormal brain development. Front Mol Neurosci. 2014;7:28. doi:10.3389/fnmol.2014.00028

101. Behnke J, Cheedalla A, Bhatt V, et al. Neuropeptide VGF Promotes Maturation of Hippocampal Dendrites That Is Reduced by Single Nucleotide Polymorphisms. Int J Mol Sci. Mar 11 2017;18(3)doi:10.3390/ijms18030612

102. Thakker-Varia S, Behnke J, Doobin D, et al. VGF (TLQP-62)-induced neurogenesis targets early phase neural progenitor cells in the adult hippocampus and requires glutamate and BDNF signaling. Stem Cell Res. May 2014;12(3):762–77. doi:10.1016/j.scr.2014.03.005

103. El Gaamouch F, Audrain M, Lin WJ, et al. VGF-derived peptide TLQP-21 modulates microglial function through C3aR1 signaling pathways and reduces neuropathology in 5xFAD mice. Mol Neurodegener. Jan 10 2020;15(1):4. doi:10.1186/s13024-020-0357-x

104. Elmadany N, de Almeida Sassi F, Wendt S, et al. The VGF-derived Peptide TLQP21 Impairs Purinergic Control of Chemotaxis and Phagocytosis in Mouse Microglia. J Neurosci. Apr 22 2020;40(17):3320–3331. doi:10.1523/JNEUROSCI.1458-19.2020

105. Cho K, Jang YJ, Lee SJ, et al. TLQP-21 mediated activation of microglial BV2 cells promotes clearance of extracellular fibril amyloid-beta. Biochem Biophys Res Commun. Apr 9 2020;524(3):764–771. doi:10.1016/j.bbrc.2020.01.111

106. Frenkel D, Maron R, Burt DS, Weiner HL. Nasal vaccination with a proteosome-based adjuvant and glatiramer acetate clears beta-amyloid in a mouse model of Alzheimer disease. J Clin Invest. Sep 2005;115(9):2423–33. doi:10.1172/JCI23241

107. Bakalash S, Pham M, Koronyo Y, et al. Egr1 expression is induced following glatiramer acetate immunotherapy in rodent models of glaucoma and Alzheimer’s disease. Invest Ophthalmol Vis Sci. Nov 21 2011;52(12):9033–46. doi:10.1167/iovs.11-7498

108. Zuroff L, Daley D, Black KL, Koronyo-Hamaoui M. Clearance of cerebral Abeta in Alzheimer’s disease: reassessing the role of microglia and monocytes. Cell Mol Life Sci. Jun 2017;74(12):2167–2201. doi:10.1007/s00018-017-2463-7

109. Lebson L, Nash K, Kamath S, et al. Trafficking CD11b-positive blood cells deliver therapeutic genes to the brain of amyloid-depositing transgenic mice. J Neurosci. Jul 21 2010;30(29):9651–8. doi:10.1523/JNEUROSCI.0329-10.2010

110. Theriault P, ElAli A, Rivest S. The dynamics of monocytes and microglia in Alzheimer’s disease. Alzheimers Res Ther. 2015;7(1):41. doi:10.1186/s13195-015-0125-2

111. Rosenzweig N, Dvir-Szternfeld R, Tsitsou-Kampeli A, et al. PD-1/PD-L1 checkpoint blockade harnesses monocyte-derived macrophages to combat cognitive impairment in a tauopathy mouse model. Nat Commun. Jan 28 2019;10(1):465. doi:10.1038/s41467-019-08352-5

112. Munoz-Castro C, Mejias-Ortega M, Sanchez-Mejias E, et al. Monocyte-derived cells invade brain parenchyma and amyloid plaques in human Alzheimer’s disease hippocampus. Acta Neuropathol Commun. Feb 28 2023;11(1):31. doi:10.1186/s40478-023-01530-z

113. Deczkowska A, Amit I, Schwartz M. Microglial immune checkpoint mechanisms. Nat Neurosci. Jun 2018;21(6):779–786. doi:10.1038/s41593-018-0145-x

114. Bui TM, Wiesolek HL, Sumagin R. ICAM-1: A master regulator of cellular responses in inflammation, injury resolution, and tumorigenesis. J Leukoc Biol. Sep 2020;108(3):787–799. doi:10.1002/JLB.2MR0220-549R

