## Supplementary materials for "Epigenetic derepression of H3K9me3 mitigates Alzheimer-related pathology and improves cognition via immunomodulation and Vgf induction"

**List of Supplementary Materials**

Supplementary Figs. S1 to S11

Supplementary Tables S1 to S8

#### Supplementary Table S1. Human donor information.

| Donor | Diagnosis | Age | Sex | Race | PMI | MMSE | CDR | Braak stage | ABC score | Aβ score (A9) | NFT score (A9) | Atrophy score (A9) |
|---|---|---|---|---|---|---|---|---|---|---|---|---|
| #1 | CN | 81 | M | H | 7.25 | 23 | 0 | 1.5 | 2.3 | 3 | 0 | 1 |
| #2 | CN | 95 | F | W | 7.25 | 30 | 0 | 5.5 | 2.7 | 2.3 | 0 | 1 |
| #3 | CN | 93 | F | W | 12 | 27 | 1 | 3.5 | 2.7 | 2.7 | 0 | 3 |
| #4 | CN | 85 | F | H | 4.5 | 30 | 0 | 1.5 | 1.7 | 2.3 | 0.5 | 0 |
| #5 | CN | 99 | F | W | 21 | 29 | 0 | 3 | 1.3 | 0.7 | 0 | 1 |
| #6 | CN | 95 | M | W | 6.5 | 30 | 0 | 1 | 1 | 0.7 | 0 | 3 |
| #7 | MCI | 94 | F | B | 10.5 | 29 | 0.5 | 1.5 | 1.7 | 2.3 | 0 | 3 |
| #8 | MCI | 87 | F | W | 4 | 13 | 3 | 5.5 | 3 | 3.7 | 3 | 3 |
| #9 | MCI | 88 | M | W | 1.5 | 28 | 3 | 3 | 1.7 | 1 | 1 | 1 |
| #10 | MCI | 85 | M | W | 16.5 | N/A | 0.5 | 1.5 | 1.3 | 2 | 0 | 0 |
| #11 | MCI | 89 | F | W | 1.5 | 24 | 0.5 | 3.5 | 1.7 | 0.5 | 0.5 | 1 |
| #12 | AD | 88 | M | W | 7.5 | 16 | 1 | 5.5 | 3 | 5 | 1 | 1 |
| #13 | AD | 93 | F | A | 9 | 20 | 3 | 3.5 | 2.3 | 1 | 0 | 3 |
| #14 | AD | 86 | F | W | 6 | 18 | 3 | 5.5 | 2.7 | 5 | 1 | 5 |
| #15 | AD | 93 | F | B | 12 | 17 | 3 | 3.5 | 2 | 3 | 0 | 1 |
| #16 | AD | 90 | F | W | 19 | N/A | 3 | 5.5 | 3 | 4.3 | 1 | 3 |
| #17 | AD | 81 | M | A | 6.5 | 12 | 3 | 5 | 3 | 5 | 5 | 5 |
| #18 | AD | 90 | M | W | 19 | N/A | 3 | 6 | 3 | 5 | 3 | 1 |
| #19 | AD | 77 | M | W | 8.25 | 18 | 2 | 6 | 3 | 5 | 3 | 1 |
| #20 | AD | 66 | F | W | 17 | 2 | 3 | 6 | 3 | 5 | 5 | 5 |
| #21 | AD | 88 | M | W | 7.5 | 18 | 1 | 5.5 | 2.3 | N/A | N/A | 1 |
| #22 | AD | 100 | F | W | 7 | 16 | 2 | 5.5 | 2.7 | 1.7 | 1 | 3 |

CN—cognitively normal, MCI—mild cognitive impairment, AD—Alzheimer's disease. Age at death in years. M—male, F—female. H—Hispanic, W—White, B—Black, A—Asian. PMI—post-mortem interval in hours. MMSE—mini mental state examination. CDR—clinical dementia rating: 0—normal cognition, 1—mild dementia, 2—moderate dementia, 3—severe dementia. ABC score: A—Aβ plaque score modified from Thal, B—NFT stage modified from Braak, C—Neuritic plaque score modified from CERAD. Aβ, NFT and atrophy severity scores: 0—none, 1—sparse, 3—moderate, 5—frequent. A9—Brodmann area A9, dorsolateral prefrontal cortex. N/A—not available.

**Supplementary Table S2. List of antibodies.**

| Primary antibody | Host | IHC dilution | WB dilution | Source | Catalog# |
|---|---|---|---|---|---|
| Annexin A2 | Rat | 1:200 | | Biolegend | 671001 |
| β-actin | Mouse | | 1:1000 | Santa Cruz | sc-47778 |
| β-amyloid 1-16 (6E10) | Mouse | 1:200-1:500 | | Biolegend | 803001 |
| β-amyloid 17-24 (4G8) | Mouse | 1:200 | | Biolegend | 800701 |
| BDNF | Mouse | 1:200 | | Abcam | ab205067 |
| CD45 | Rat | 1:25 | | BD Pharmagen | 550539 |
| GFAP | Goat | 1:500 | | Novus | NB100-53809 |
| GFAP | Rat | | 1:1000 | Invitrogen | 13-0300 |
| H3K9me3 | Rabbit | 1:500 | 1:1000 | Abcam | ab8898 |
| Iba1 | Rabbit | 1:250 | | Wako | 019-19741 |
| Iba1 | Goat | 1:250 | | Novus | NB-100-1028 |
| MT2A | Rabbit | 1:100 | | Invitrogen | PA5-76652 |
| NeuN | Mouse | 1:1000 | | Abcam | ab177487 |
| PLTP | Sheep | 1:40 | | R&D system | AF4918 |
| PSD95 | Rabbit | 1:600 | | Abcam | ab76115 |
| Tau phospho-Ser396/Ser404 (PHF-tau) | Mouse | 1:200 | | Provided by Prof. Peter Davies lab | |
| VGF | Rabbit | 1:200 | | Novus | NBP1-76906 |
| **Secondary antibody** | | | | | |
| Cy2 | Goat, Mouse, Rabbit, Rat, Sheep | 1:200 | | Jackson ImmunoResearch Laboratories | |
| Cy3 | | 1:200 | | | |
| Cy5 | | 1:200 | | | |
| Cy7 | | 1:200 | | | |
| HRP | Mouse, Rabbit, Rat | | 1:10000 | | |

IHC, Immunohistochemistry; WB, Western blotting.

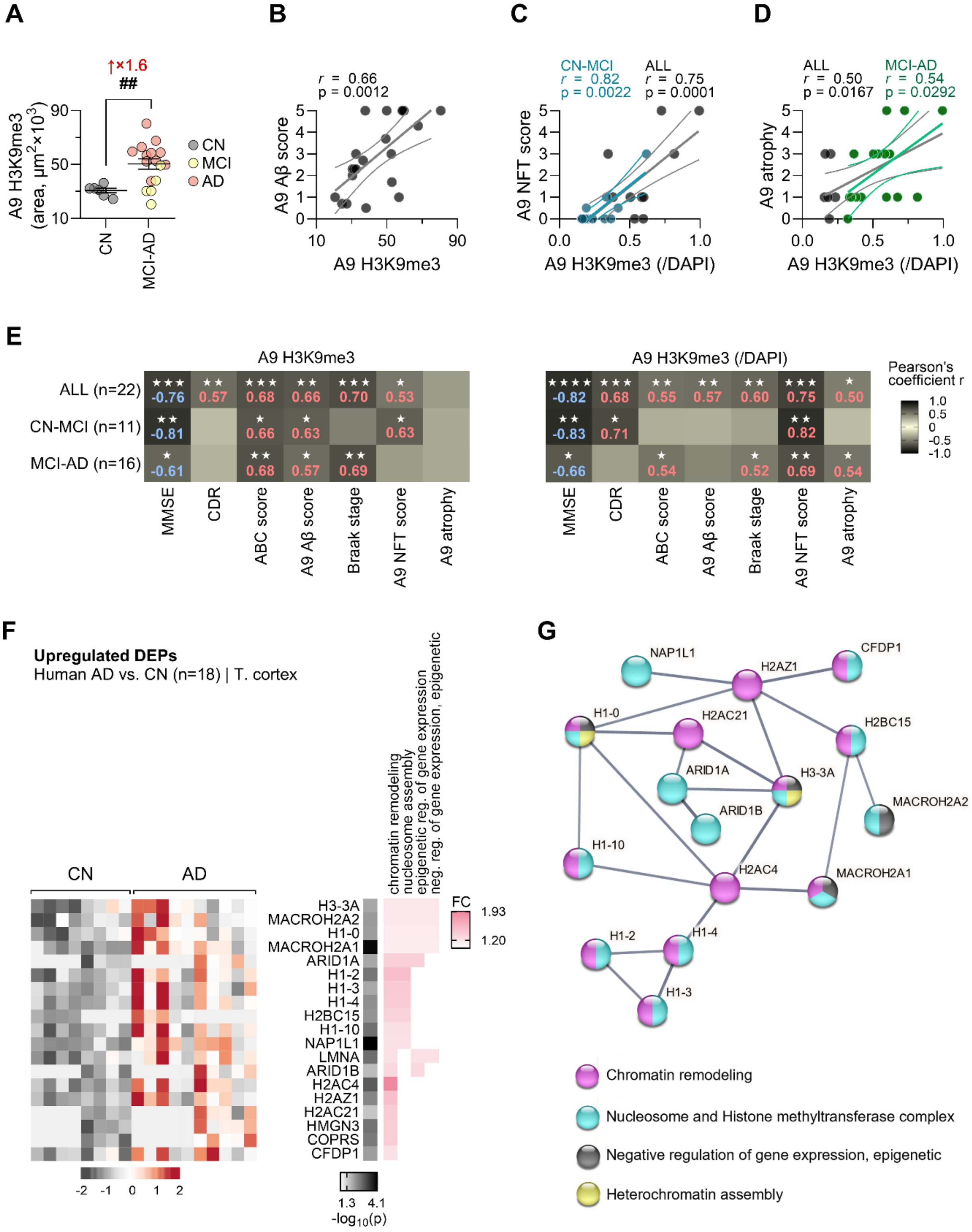
A
↑×1.6
##
A9 H3K9me3 (area, μm²×10³)
90
50
10
CN
MCI-AD
CN
MCI
AD
B
r = 0.66
p = 0.0012
A9 Aβ score
A9 H3K9me3
C
CN-MCI
r = 0.82
p = 0.0022
ALL
r = 0.75
p = 0.0001
A9 NFT score
A9 H3K9me3 (/DAPI)
D
ALL
r = 0.50
p = 0.0167
MCI-AD
r = 0.54
p = 0.0292
A9 atrophy
A9 H3K9me3 (/DAPI)
E
A9 H3K9me3
ALL (n=22)
CN-MCI (n=11)
MCI-AD (n=16)
-0.76 0.57 0.68 0.66 0.70 0.53
-0.81 0.66 0.63 0.63
-0.61 0.68 0.57 0.69
MMSE
CDR
ABC score
A9 Aβ score
Braak stage
A9 NFT score
A9 atrophy
A9 H3K9me3 (/DAPI)
-0.82 0.68 0.55 0.57 0.60 0.75 0.50
-0.83 0.71 0.82
-0.66 0.54 0.52 0.69 0.54
Pearson's coefficient r
1.0
0.5
0
-0.5
-1.0
F
Upregulated DEPs
Human AD vs. CN (n=18) | T. cortex
CN
AD
chromatin remodeling
nucleosome assembly
epigenetic reg. of gene expression
neg. reg. of gene expression, epigenetic
H3-3A
MACROH2A2
H1-0
MACROH2A1
ARID1A
H1-2
H1-3
H1-4
H2BC15
H1-10
NAP1L1
LMNA
ARID1B
H2AC4
H2AZ1
H2AC21
HMGN3
COPRS
CFDP1
FC
1.93
1.20
-2 -1 0 1 2
1.3 4.1
-log10(p)
G
NAP1L1
H2AZ1
CFDP1
H1-0
H2AC21
H2BC15
ARID1A
H3-3A
ARID1B
MACROH2A2
H1-10
H2AC4
MACROH2A1
H1-2
H1-4
H1-3
Chromatin remodeling
Nucleosome and Histone methyltransferase complex
Negative regulation of gene expression, epigenetic
Heterochromatin assembly

**Supplementary Fig. S1. Increased cerebral H3K9me3 correlates with cognitive deficits and AD-related neuropathology**. (**A**) Quantification of H3K9me3 immunoreactivity (IR area, $\mu m^2 \times 10^3$). ## $p < 0.01$ by two-tailed unpaired Student's t-test between AD patients, which include MCI patients (early stage of AD), and CN subjects. (**B**) Pearson's correlation coefficient between A9 H3K9me3 IR and A9 Aβ scores, (**C–D**) Pearson's correlation coefficients of A9 H3K9me3 IR (corrected for DAPI nuclei count) with (C) A9 NFT scores and (D) A9 atrophy scores. (**E**) Summary heatmaps of Pearson's correlation coefficients of A9 H3K9me3 IR (uncorrected and corrected for DAPI nuclei count) with MMSE, CDR, ABC scores, A9 Aβ scores, Braak stages, A9 NFT scores, and A9 atrophy scores, for the entire cohort of subjects, or limited to CN-MCI or MCI-AD individuals. (**F**) Expression profiles of upregulated DEPs in AD patients, involved in chromatin organization and epigenetic regulation of gene expression. (**C**) Protein association network showing select protein roles in chromatin remodeling and modification using the STRING database (v11.5). Nodes (proteins) are color-coded according to function.

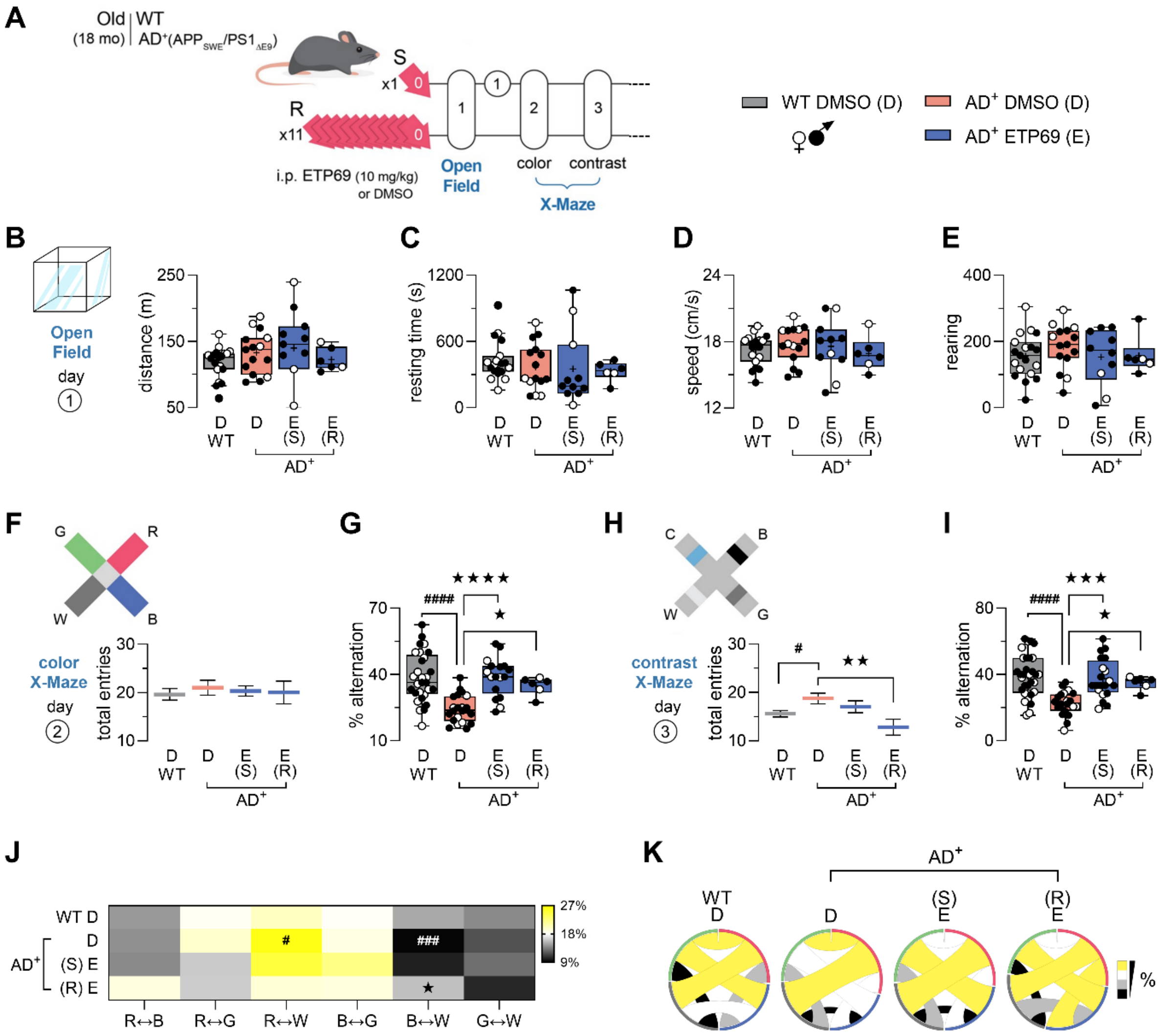

**Supplementary Fig. S2. Effects of single or repeated i.p. injections of ETP69 on locomotor activity, cognition, and vision in 18-month-old WT and AD model mice.** (**A**) Experimental timeline and schedule of i.p. injection of ETP69 (10 mg/kg) or DMSO in 18-month-old AD$^{+}$ mice and age/sex-matched WT littermates. Regimens: S, single; R, repeated. (**B**) Distance travelled, (**C**) resting time, (**D**) average speed, and (**E**) rearing number during the 30 min-long open field test. (**F**) Total number of entries (locomotor activity) and (**G**) percentage of alternations (cognition/vision) in the colour mode of the visual-stimuli X-maze test. (**H**) Number of entries and (**I**) percent alternations in the contrast mode of the X-maze test. (**J**) Percentage and (**K**) chord diagrams of bidirectional transitions between arms in the colour mode of the X-maze test. Circled numbers represent the days of behavioural testing according to the experimental timeline. Group means ± SEMs are presented. Individual data points and median as well as lower and upper quartiles are shown on each box-and-whisker plot. Filled and empty circles represent male and female mice, respectively. # $p < 0.05$, ### $p < 0.001$, and #### $p < 0.0001$: DMSO-injected $AD^{+}$ mice versus DMSO-injected WT mice; ★ $p < 0.05$, ★★★ $p < 0.001$, and ★★★★ $p < 0.0001$: ETP69-treated mice versus DMSO-injected mice; by one-way ANOVA followed by Fisher's least significant difference (LSD) *post hoc* test.

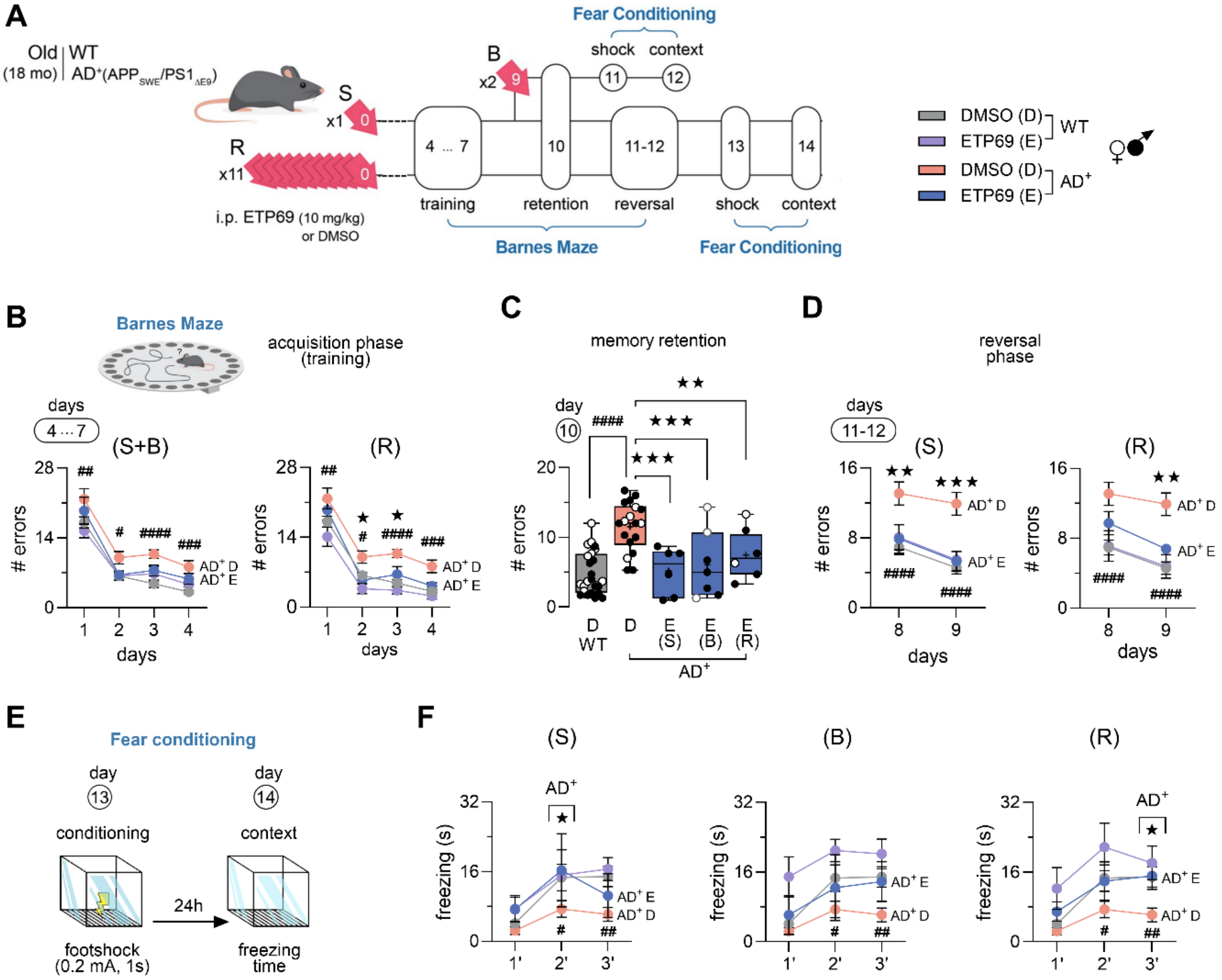


**Supplementary Fig. S3. Effects of different administration regimens of i.p. ETP69 on cognitive function in 18-month-old WT and AD model mice.** (**A**) Experimental timeline and schedule of the i.p. injection of ETP69 (10 mg/kg) or DMSO in 18-month-old AD$^+$ mice and age- and sex-matched WT littermates. Regimens: S, single; R, repeated; and B, booster. (**B**–**D**) Number of errors before finding the escape box are shown for S, R, and B regimens (B) across the 4 training days, (C) on the 7th day (memory retention) of the Barnes maze test, and (D) during the reversal phase of the Barnes maze. (**E**) Conditions of the contextual fear conditioning test. (**F**) Freezing time over a 3-min period in the context-specific fear conditioning test. Circled numbers represent the day(s) of behavioural testing according to the experimental timeline. Group means ± SEMs are presented. Individual data points and median as well as lower and upper quartiles are indicated on each box-and-whisker plot. Filled and empty circles represent male and female mice, respectively. # $p < 0.05$, ## $p < 0.01$, ### $p < 0.001$, and #### $p < 0.0001$: DMSO-injected $AD^+$ mice versus DMSO-injected WT mice; ★ $p < 0.05$, ★★ $p < 0.01$, and ★★★ $p < 0.001$: ETP69-treated mice versus DMSO-injected mice; by one-way or two-way ANOVA followed by Fisher's LSD *post hoc* test.

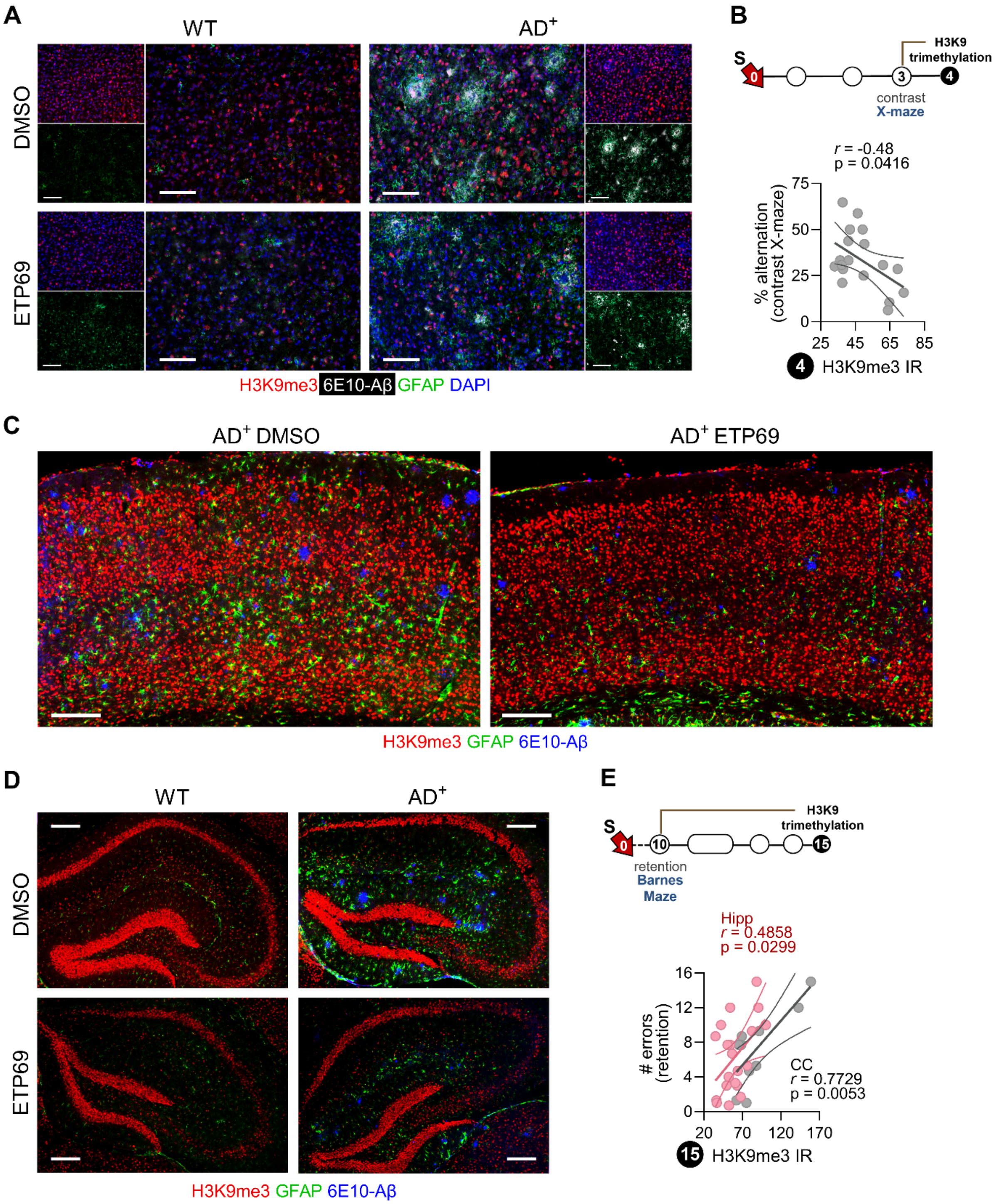


**Supplementary Fig. S4. Effects of a single i.p. injection of ETP69 on H3K9me3 and AD pathological markers.** (**A**) Representative images of immunofluorescent staining for H3K9me3 (red), 6E10 (white, Aβ plaques), and GFAP (green, reactive astrocytes) in the cerebral cortices of

18-month-old $AD^+$ and WT mice 4 days after i.p. injection of ETP69 (10 mg/kg) or DMSO. Scale bar: 100 μm. (**B**) Timeline showing elapsed time and Pearson's correlation coefficient between cortical H3K9me3 immunoreactivity (IR) and performance in the contrast mode of the X-maze test. (**C–D**) Representative images of immunofluorescent staining for H3K9me3 (red), 6E10 (blue), and GFAP (green) in the (C) cerebral cortices ($AD^+$) and (D) hippocampi (WT and $AD^+$) of 18-month-old mice 15 days after i.p. injection of ETP69 (10 mg/kg) or DMSO. Scale bars: 200 μm. (**E**) Timeline showing elapsed time and Pearson's correlations between performance in the memory retention phase of the Barnes maze test and H3K9me3 IR in both the cerebral cortex (CC) and hippocampus (Hipp). Black circled numbers represent the end-point day according to the experimental timeline.

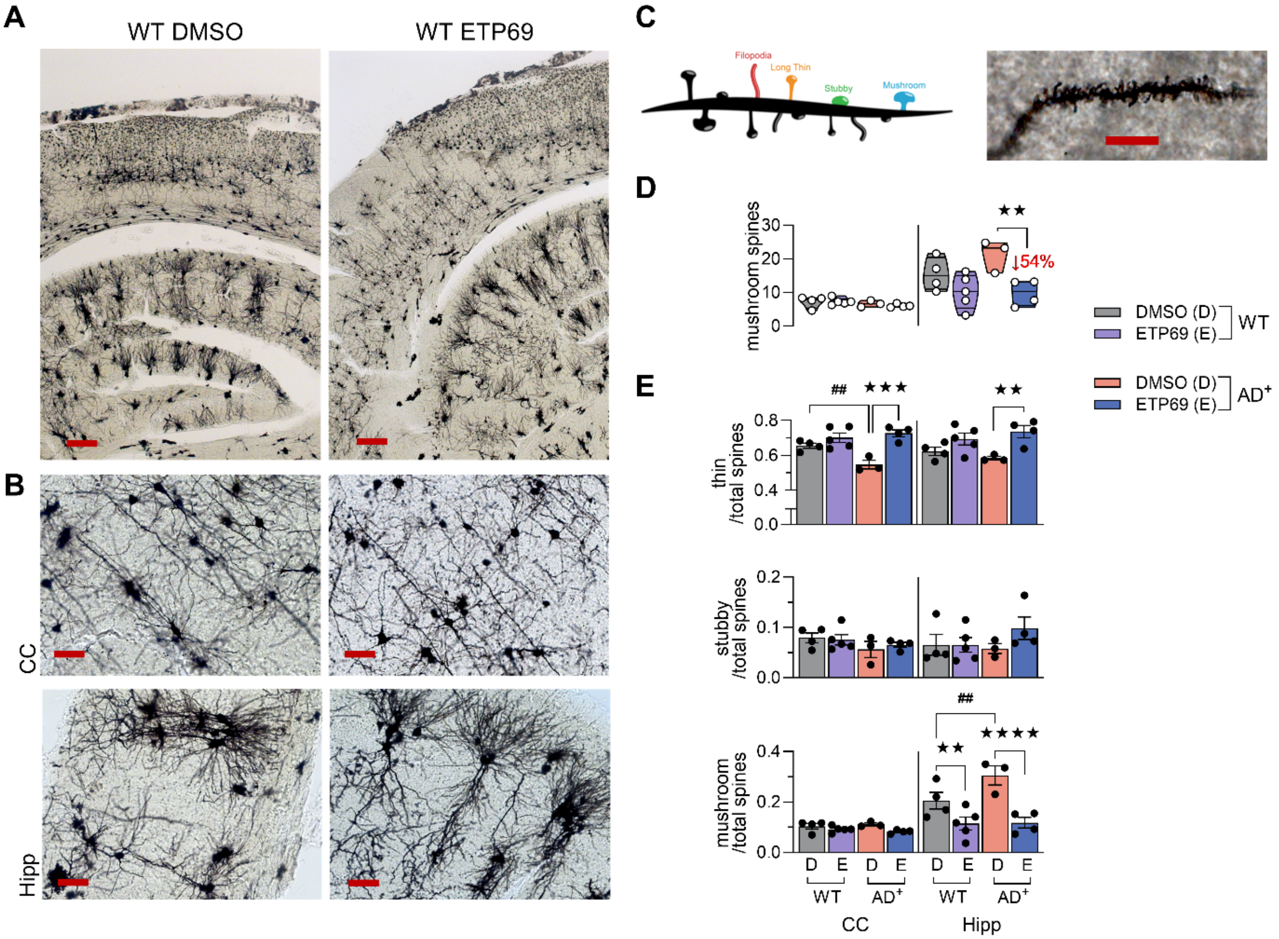


**Supplementary Fig. S5. Effects of ETP69 administration on dendritic spine subtypes.** (**A**) Low- (scale bars: 200 µm) and (**B**) high-magnification (scale bars: 50 µm) representative images of Golgi-Cox staining of neurons in the cerebral cortex (CC) and hippocampus (Hipp) of 18-month-old WT mice following i.p. injection of ETP69 (10 mg/kg) or DMSO. (**C**) Illustration and high-magnification (scale bar: 10 µm) photograph of dendritic spines classified as filopodia-like, long-thin, stubby, or mushroom spines according to size and shape. (**D**) Density of mushroom spines as counts per 100 µm of dendrite in the cerebral cortex (CC) and hippocampus (Hipp). (**E**) Ratios of thin, mushroom, and stubby spines relative to total dendritic spines. Individual data points are presented with group means ± SEMs. Median and lower and upper quartiles are indicated on each violin plot. ## $p < 0.01$: DMSO-injected $AD^+$ mice versus DMSO-injected WT mice; ★★ $p < 0.01$, ★★★ $p < 0.001$, and ★★★★ $p < 0.0001$: ETP69-treated mice versus DMSO-injected mice; by one-way ANOVA followed by Fisher's LSD *post hoc* test.

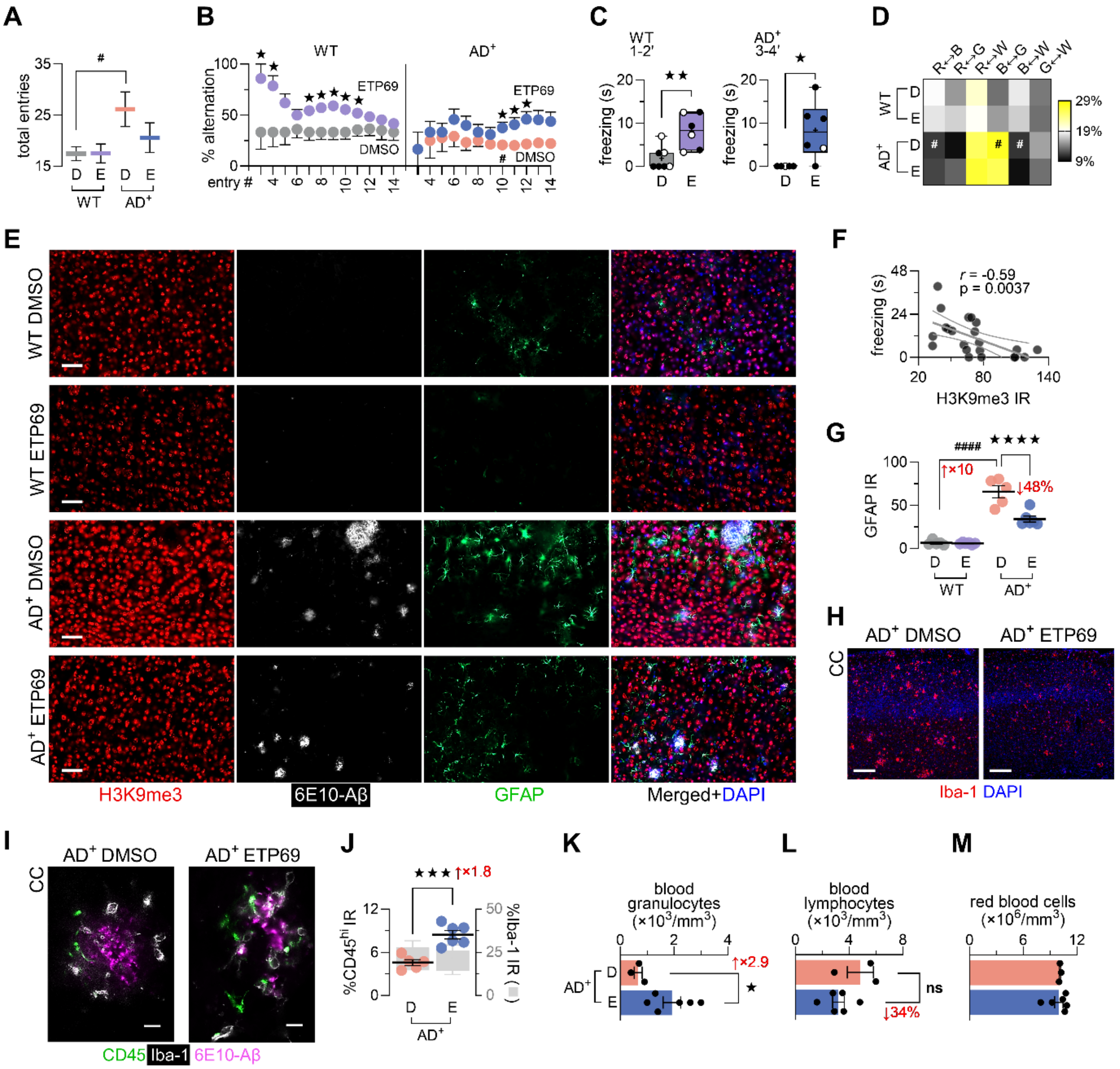

**Supplementary Fig. S6. Therapeutic effects of a single injection of ETP69 in 14-month-old WT and AD model mice.** (**A–D**) Behavioral testing of 14-month-old AD⁺ mice and age- and sex-matched WT littermates following a single i.p. injection of ETP69 (10 mg/kg) or DMSO (control). (A) Total number of entries (locomotor activity) in the colour mode of the X-maze test. (B) Progression in percentage of alternations in the X-maze test with each additional entry from the start of the test until the mouse reached 14 entries. (C) Cumulative freezing times of WT and AD⁺

mice in the contextual fear conditioning test. (D) Percentage of bidirectional transitions between maze arms in the colour mode of the X-maze test. (**E**) Representative images (scale bars: 50 μm) of immunofluorescent staining for H3K9me3 (red), 6E10 (white), and GFAP (green) in the cerebral cortex of 14-month-old $AD^+$ and WT mice, 4 days after a single i.p. injection of ETP69 or DMSO. (**F**) Pearson's correlation coefficient of cortical H3K9me3 levels with performance in the context-specific fear conditioning test. (**G**) Quantitative analysis of GFAP IR ($\mu m^2 \times 10^3$) in the cerebral cortex. (**H**) Representative images of immunofluorescent staining (scale bar: 200 μm) for Iba-1 (red, activated microglia) and DAPI (blue, nuclei). (**I**) Representative images (scale bars: 10 μm) of immunofluorescent staining for Iba-1 (white), CD45 (green) and 6E10 (purple) in the cerebral cortex. (**J**) Quantitative analysis of $CD45^{hi}$ and Iba-1 IR (% area) near amyloid plaques in the cerebral cortex. (**K–M**) Cell counts, including (K) granulocytes, (L) lymphocytes, and (M) red blood cells, in the blood of 18-month-old $AD^+$ mice administered ETP69 or DMSO. Individual data points are presented with group means ± SEMs. Lower and upper quartiles are indicated in the box-and-whisker plot. # $p < 0.05$ and #### $p < 0.0001$: DMSO-injected $AD^+$ mice versus DMSO-injected WT mice; ★ $p < 0.05$, ★★ $p < 0.01$, ★★★ $p < 0.001$, and ★★★★ $p < 0.0001$: ETP69-treated mice versus DMSO-injected mice; by one-way or two-way ANOVA followed by Fisher's LSD *post hoc* test or two-tailed unpaired Student's t-test.

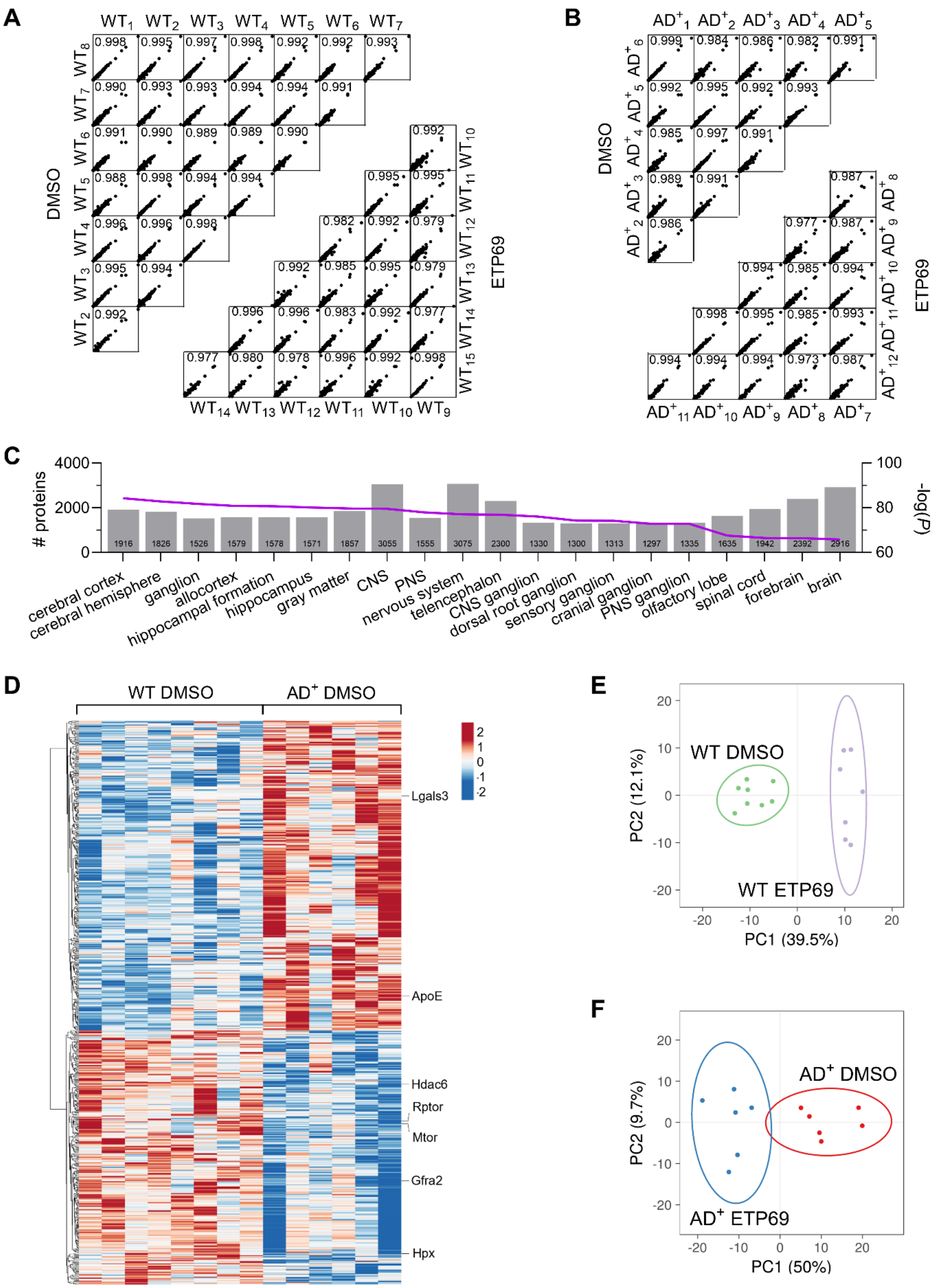
A
DMSO
ETP69
B
DMSO
ETP69
C
### proteins
-log(P)
cerebral cortex
cerebral hemisphere
ganglion
allocortex
hippocampal formation
hippocampus
gray matter
CNS
PNS
nervous system
telencephalon
CNS ganglion
dorsal root ganglion
sensory ganglion
cranial ganglion
PNS ganglion
olfactory lobe
spinal cord
forebrain
brain
D
WT DMSO
AD+ DMSO
Lgals3
ApoE
Hdac6
Rptor
Mtor
Gfra2
Hpx
E
WT DMSO
WT ETP69
PC1 (39.5%)
PC2 (12.1%)
F
AD+ DMSO
AD+ ETP69
PC1 (50%)
PC2 (9.7%)

**Supplementary Fig. S7. Reproducibility and clustering of MS proteomics data from 14-month-old WT and AD model mice.** (**A–B**) Pearson's correlation coefficients for all the quantified proteins between animals within each group for (A) WT groups and (B) $AD^+$ groups. Pearson's coefficients greater than 0.973 between animals from the same group indicate high reproducibility of the MS data. (**C**) Anatomical enrichment analysis of the 4358 proteins quantified by MS. CNS, central nervous system; PNS, peripheral nervous system. (**D**) Heatmap displaying the hierarchical clustering of all significant DEPs between ETP69-treated WT mice and DMSO-injected (control) WT mice. Blue: downregulated proteins. Red: upregulated proteins. (**E–F**) Principal component analysis of protein expression profiles in (E) WT mice and (F) $AD^+$ mice after ETP69 or DMSO treatment. All significant DEPs were included in the analysis (WT mice: 318 proteins, $AD^+$ mice: 370 proteins).

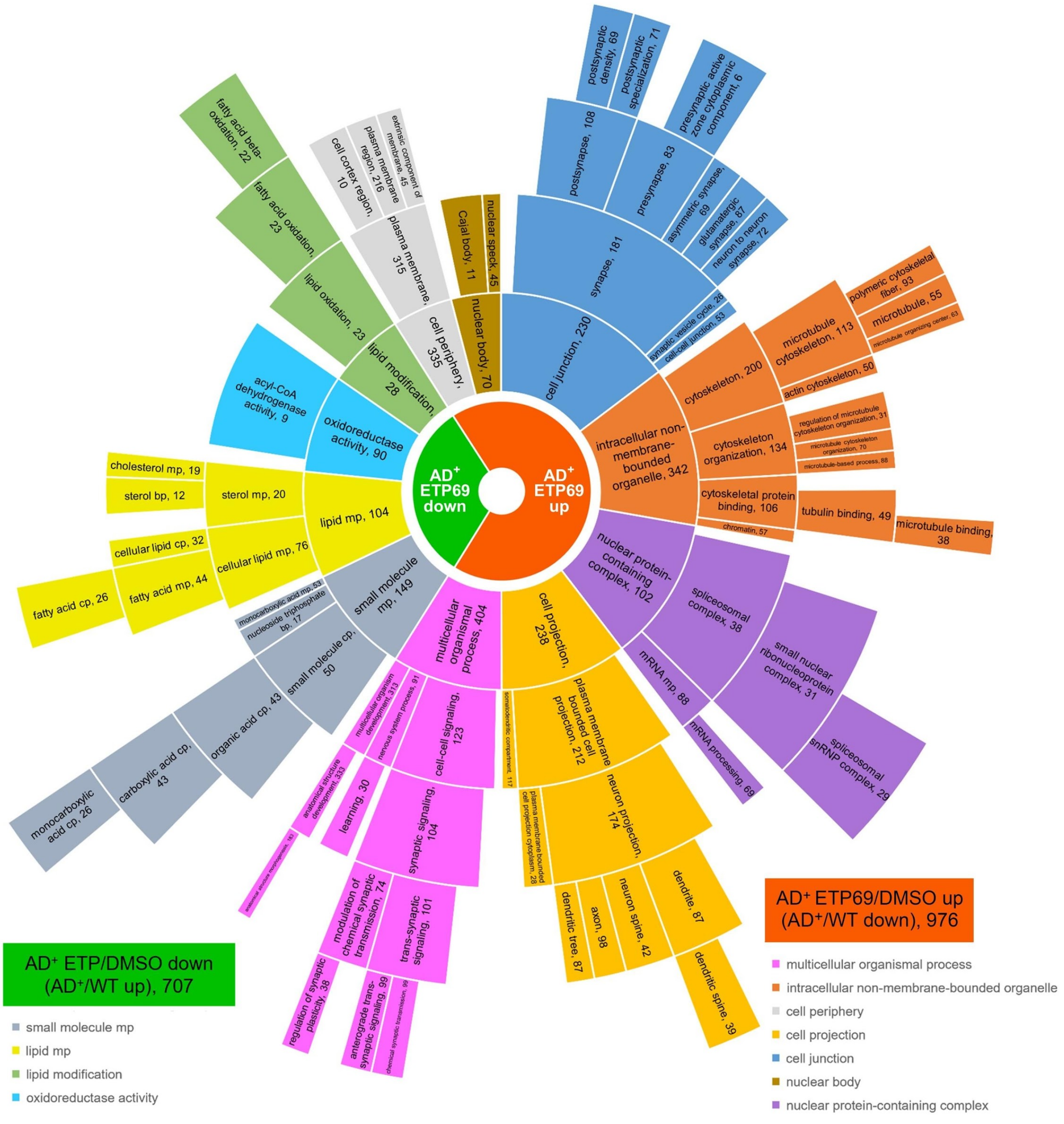


**Supplementary Fig. S8. Select GO terms.** Select GO terms (combination of biological process, cellular compartment, and molecular function terms) displayed as a sunburst chart. GO enrichment analysis of proteins that showed reversal of expression in AD$^{+}$ mice after treatment with ETP69. All proteins with a |FC| > 1.04 were included in this analysis. Cell sizes are relative to the total number of proteins for each GO term.

**A**

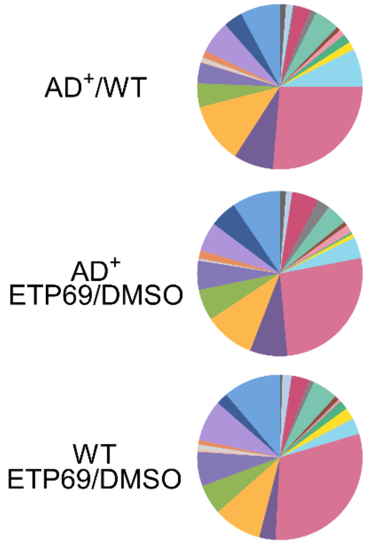


| | AD+/WT | AD+ ETP69/DMSO | WT ETP69/DMSO |
|---|---|---|---|
| metabolite interconversion enzyme | 26.5% | 26.5% | 30.6% |
| protein modifying enzyme | 11.9% | 9.8% | 9.5% |
| nucleic acid metabolism protein | 7.8% | 7.3% | 3.2% |
| transporter | 7.8% | 9.4% | 11.3% |
| membrane traffic protein | 7.4% | 4.1% | 3.2% |
| translational protein | 6.4% | 5.7% | 8.1% |
| cytoskeletal protein | 4.9% | 4.1% | 5.0% |
| protein-binding activity modulator | 4.7% | 6.1% | 5.9% |
| scaffold/adaptor protein | 4.1% | 5.7% | 6.8% |
| transmembrane signal receptor | 3.7% | 5.3% | 2.3% |
| chaperone | 3.3% | 5.3% | 3.6% |
| gene-specific transcriptional regulator | 1.8% | 0.4% | 1.8% |
| intercellular signal molecule | 1.6% | 0.8% | 2.3% |
| cell adhesion molecule | 1.4% | 1.2% | 1.8% |
| transfer/carrier protein | 1.4% | 1.6% | 0.9% |
| calcium-binding protein | 1.2% | 1.2% | 0.5% |
| chromatin/chromatin-binding | 1.2% | 2.4% | 0.9% |
| extracellular matrix protein | 1.2% | 1.6% | 0.5% |
| defense/immunity protein | 0.8% | 0.8% | 0.9% |
| structural protein | 0.6% | 0.4% | 0.9% |
| storage protein | 0.4% | - | 0.5% |
| cell junction protein | 0.2% | - | - |

**B**

| Symbol | FC |
|---|---|
| Slc4a1 | -2.67 |
| Mapt | -1.85 |
| Mea1 | -1.81 |
| Slc39a7 | -1.55 |
| **Gfra2** | **-1.46** |
| Mtco1 | -1.45 |
| Prpf40a | -1.45 |
| Bcas2 | -1.45 |
| Nop2 | -1.44 |
| Prkcq | -1.43 |
| Plcb4 | -1.43 |
| Pwwp3a | -1.42 |
| Pik3r3 | -1.42 |
| Pde4a | -1.41 |
| **Cit** | **-1.39** |
| **Efr3a** | **-1.38** |
| Zmat4 | -1.38 |
| **Vav3** | **-1.37** |
| Eogt | -1.37 |
| Ccdc136 | -1.37 |

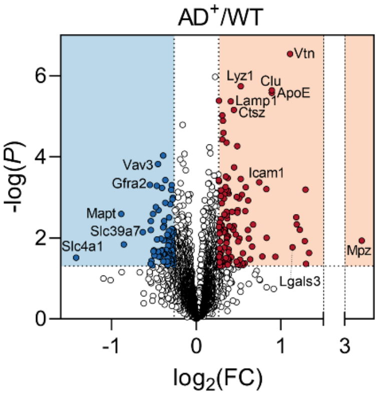


| Symbol | FC |
|---|---|
| Mpz | 9.77 |
| Gh1 | 2.51 |
| Olfr703 | 2.45 |
| Pde10a | 2.44 |
| Ccdc126 | 2.43 |
| Mdk | 2.33 |
| Ahnak2 | 2.27 |
| Clstn3 | 2.27 |
| Lgals3 | 2.20 |
| **Vtn** | **2.15** |
| Slc6a5 | 1.91 |
| **ApoE** | **1.86** |
| **Clu** | **1.85** |
| Borcs5 | 1.78 |
| Ifi204 | 1.78 |
| Vmn2r3 | 1.71 |
| **Icam1** | **1.67** |
| Crym | 1.59 |
| Cnn1 | 1.58 |
| Rpl14 | 1.56 |

**C**

| Symbol | FC |
|---|---|
| **Tnks** | **-1.54** |
| Mtco1 | -1.50 |
| Prkcq | -1.41 |
| Papss2 | -1.36 |
| Pcsk1n | -1.34 |
| Nlgn4l | -1.34 |
| Kcnt1 | -1.32 |
| **Zmat4** | **-1.32** |
| Slc17a6 | -1.32 |
| **Guf1** | **-1.31** |
| Rbm8a | -1.28 |
| **Kcnab3** | **-1.27** |
| Ctbs | -1.27 |
| Nup214 | -1.26 |
| Ppp1r3g | -1.24 |
| Ccdc136 | -1.24 |
| Cask | -1.23 |
| Bod1 | -1.23 |
| Slc38a3 | -1.23 |
| Fam169a | -1.22 |

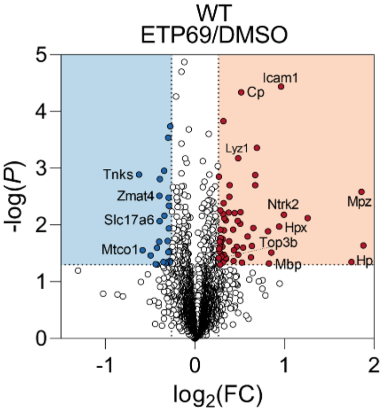


| Symbol | FC |
|---|---|
| Ints3 | 3.69 |
| Mpz | 3.63 |
| Hp | 3.36 |
| Ahnak2 | 2.39 |
| Ntrk2 | 1.99 |
| **Icam1** | **1.95** |
| Hpx | 1.92 |
| Fcgr2 | 1.81 |
| Mbp | 1.77 |
| Pde10a | 1.76 |
| **H4c1** | **1.61** |
| Ahsg | 1.60 |
| **Ppp1r1b** | **1.60** |
| Borcs5 | 1.57 |
| Chd7 | 1.55 |
| Smc | 1.54 |
| Cnn1 | 1.50 |
| Abcb8 | 1.45 |
| Cavin3 | 1.44 |
| **Cp** | **1.43** |

**Supplementary Fig. S9. MS proteomics analysis of 14-month-old WT and AD-model mice.** (**A**) PANTHER categories for all significant DEPs in $AD^+$ mice, in ETP69-treated (versus DMSO-injected) $AD^+$ mice, and in ETP69-treated (versus DMSO-injected) WT mice. (**B–C**) Volcano plots of all DEPs in (B) $AD^+$ (versus WT) mice and (C) ETP69-treated (versus DMSO-injected) WT mice. Red: upregulated proteins ($FC > 1.2$, $p < 0.05$). Blue: downregulated proteins ($FC < -1.2$, $p < 0.05$). The top 20 downregulated and upregulated proteins are listed.

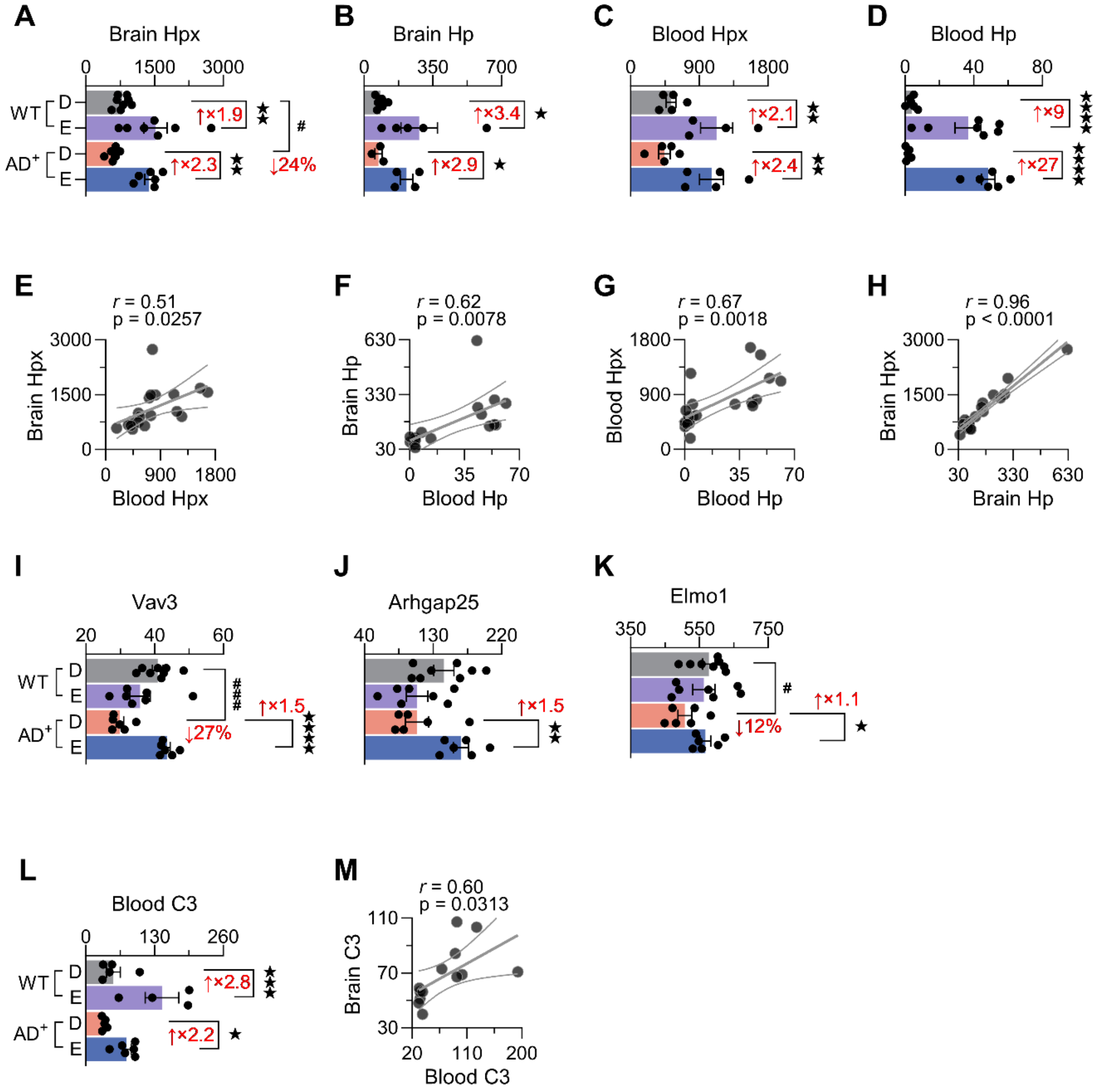


**Supplementary Fig. S10. Innate immune response and changes associated with Aβ clearance, microglial phagocytosis, and leukocyte recruitment in WT and AD-model mice.** (**A–B**) Quantification of (A) Hpx and (B) Hp protein expression by MS in 14-month-old AD$^{+}$ and WT mice in response to ETP69 or DMSO (control) administration. (**C–D**) Quantification of blood (C) Hpx and (D) Hp levels. (**E–H**) Pearson's correlation coefficient between brain and blood (E) Hpx and (F) Hp levels, and between Hpx and Hp levels in (G) the blood and (H) brain. (**I–K**) Quantification of (I) Vav3, (J) Arhgap25, and (K) Elmo1 protein expression by MS in 14-month-old WT and AD$^+$ mice in response to ETP69 or DMSO administration. (**L**) Quantification of blood C3 levels. (**M**) Pearson's correlation coefficient between brain and blood C3 levels. Individual data points are presented with group means ± SEMs. # $p < 0.05$, and ### $p < 0.001$: DMSO-injected AD$^+$ mice versus DMSO-injected WT mice; ★ $p < 0.05$, ★★ $p < 0.01$, and ★★★★ $p < 0.0001$: ETP69-treated mice versus DMSO-injected mice; by one-way ANOVA followed by Fisher's LSD *post hoc* test.

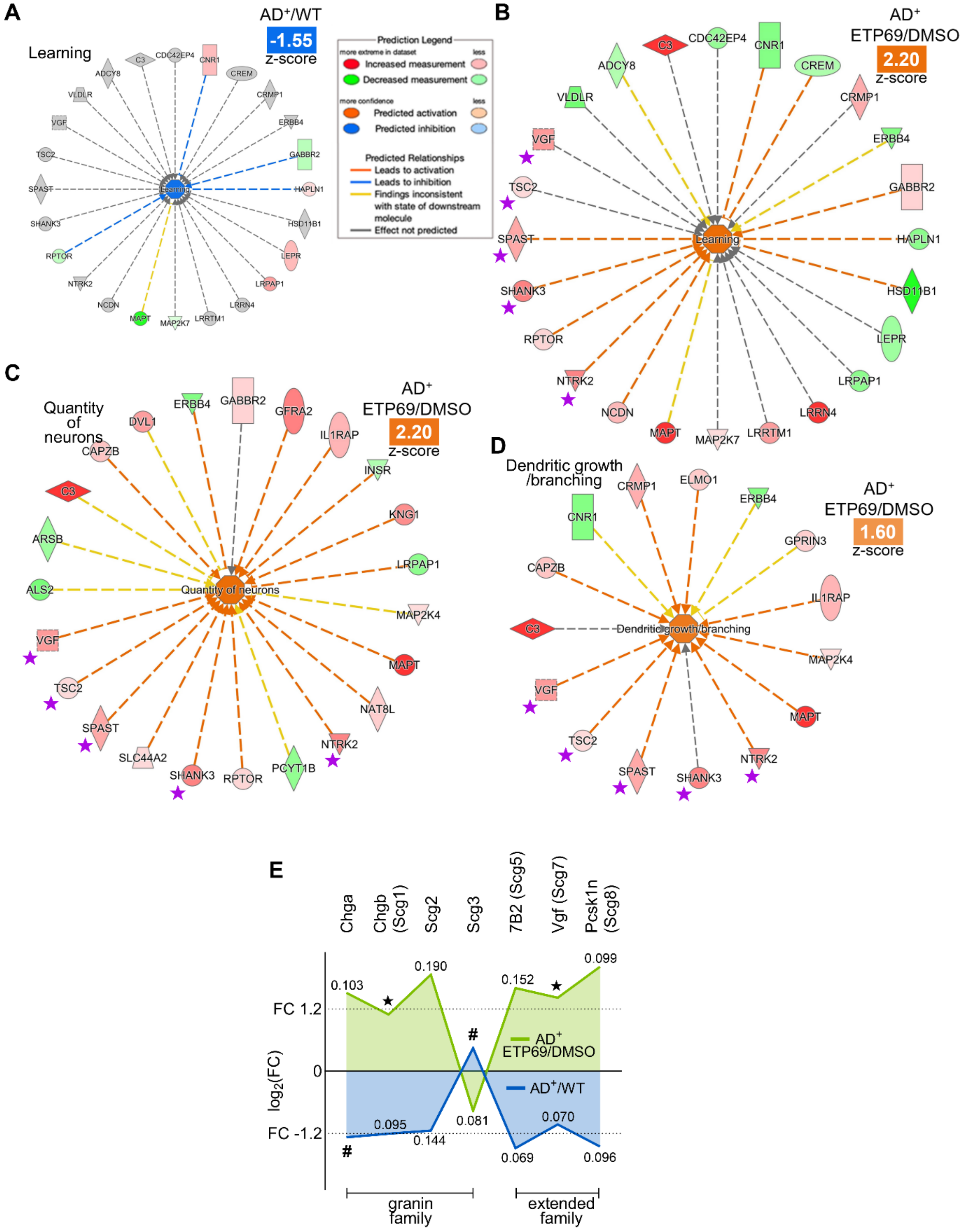
A
Learning
AD+/WT
-1.55
z-score
Prediction Legend
more extreme in dataset
less
Increased measurement
Decreased measurement
more confidence
less
Predicted activation
Predicted inhibition
Predicted Relationships
Leads to activation
Leads to inhibition
Findings inconsistent with state of downstream molecule
Effect not predicted
B
AD+ ETP69/DMSO
2.20
z-score
Learning
C3
CDC42EP4
CNR1
ADCY8
CREM
VLDLR
CRMP1
VGF
ERBB4
TSC2
GABBR2
SPAST
HAPLN1
SHANK3
HSD11B1
RPTOR
LEPR
NTRK2
LRPAP1
NCDN
LRRN4
MAPT
MAP2K7
LRRTM1
C
Quantity of neurons
AD+ ETP69/DMSO
2.20
z-score
DVL1
ERBB4
GABBR2
GFRA2
IL1RAP
CAPZB
INSR
C3
KNG1
ARSB
LRPAP1
ALS2
MAP2K4
VGF
MAPT
TSC2
NAT8L
SPAST
NTRK2
SLC44A2
SHANK3
RPTOR
PCYT1B
D
Dendritic growth /branching
AD+ ETP69/DMSO
1.60
z-score
CRMP1
ELMO1
ERBB4
CNR1
GPRIN3
CAPZB
IL1RAP
C3
MAP2K4
VGF
MAPT
TSC2
SPAST
SHANK3
NTRK2
E
Chga
Chgb (Scg1)
Scg2
Scg3
7B2 (Scg5)
Vgf (Scg7)
Pcsk1n (Scg8)
FC 1.2
0
FC -1.2
log2(FC)
0.103
0.190
0.152
0.099
0.095
0.144
0.081
0.070
0.069
0.096
AD+ ETP69/DMSO
AD+/WT
granin family
extended family

**Supplementary Fig. S11. MS proteomics analysis of 14-month-old WT and AD model mice.** (**A–B**) Ingenuity Pathway Analysis (IPA) diagrams of DEPs associated with learning in (A) $AD^+$ (versus WT) mice and (B) ETP69-treated (versus DMSO-injected) $AD^+$ mice. (**C–D**) IPA diagrams of DEPs in ETP69-treated (versus DMSO-injected) $AD^+$ mice associated with (C) quantity of neurons and (D) dendritic growth/branching. Purple stars indicate the five overlapping proteins in the three pathways: Ntrk2, Shank3, Spast, Tsc2, and Vgf. (**E**) Members of the extended granin family differentially expressed in ETP69-treated versus DMSO-injected $AD^+$ mice (green line) and in $AD^+$ versus WT mice (blue line). Group means ± SEMs are presented. # $p < 0.05$: DMSO-injected $AD^+$ mice versus DMSO-injected WT mice; ★ $p < 0.05$: ETP69-treated $AD^+$ mice versus DMSO-injected $AD^+$ mice; by two-tailed unpaired Student's t test.

**Supplementary Table S3. Significant brain DEPs upregulated in ETP69-treated AD$^+$ mice versus DMSO-injected (control) AD$^+$ mice.**

| Accession | Symbol | Description | AD$^+$ ETP69 (Mean) | AD$^+$ DMSO (Mean) | FC | p-value |
|---|---|---|---|---|---|---|
| Q61646 | HP | Haptoglobin | 216.0 | 73.4 | 2.94 | 0.0170 |
| P10637-3 | MAPT | Isoform Tau-B of Microtubule-associated protein tau | 56.0 | 23.5 | 2.38 | 0.0188 |
| Q91X72 | HPX | Hemopexin | 1384.5 | 599.6 | 2.31 | 0.00003 |
| Q9JJV4 | CACNG4 | Voltage-dependent calcium channel gamma-4 subunit | 18.5 | 10.1 | 1.84 | 0.0097 |
| P59383 | LRRN4 | Leucine-rich repeat neuronal protein 4 | 247.9 | 142.9 | 1.73 | 0.0149 |
| P01027 | C3 | Complement C3 | 82.9 | 48.1 | 1.72 | 0.0265 |
| Q9DBD0 | ICA | Inhibitor of carbonic anhydrase | 253.8 | 152.1 | 1.67 | 0.0279 |
| O88492 | PLIN4 | Perilipin-4 | 613.6 | 375.4 | 1.63 | 0.0047 |
| Q80TT2 | BAIAP3 | BAI1-associated protein 3 | 315.7 | 198.0 | 1.59 | 0.0243 |
| A0A1Y7VNZ6 | MARK3 | Non-specific serine/threonine protein kinase | 103.0 | 66.4 | 1.55 | 0.0252 |
| A2ALK8 | PTPN3 | Tyrosine-protein phosphatase non-receptor type 3 | 68.7 | 44.3 | 1.55 | 0.0007 |
| Q61838 | PZP | Pregnancy zone protein | 209.9 | 135.7 | 1.55 | 0.0240 |
| Q8C0P5 | CORO2A | Coronin-2A | 790.0 | 512.2 | 1.54 | 0.00003 |
| Q9JLR1 | SEC61A2 | Protein transport protein Sec61 subunit alpha isoform 2 | 30.2 | 19.6 | 1.54 | 0.0018 |
| Q8BYW1 | ARHGAP25 | Rho GTPase-activating protein 25 | 166.5 | 109.0 | 1.53 | 0.0098 |
| Q64327 | MEA1 | Male-enhanced antigen 1 | 62.0 | 41.1 | 1.51 | 0.0474 |
| Q9R0C8 | VAV3 | Guanine nucleotide exchange factor VAV3 | 43.6 | 29.9 | 1.46 | 0.000002 |
| P49025 | CIT | Citron Rho-interacting kinase | 2577.5 | 1845.0 | 1.40 | 0.0023 |
| E9PV24 | FGA | Fibrinogen alpha chain | 79.0 | 56.6 | 1.40 | 0.0172 |
| Q9Z321 | TOP3B | DNA topoisomerase 3-beta-1 | 33.5 | 24.1 | 1.39 | 0.0057 |
| Q9D2G5 | SYNJ2 | Synaptojanin-2 | 272.1 | 196.9 | 1.38 | 0.0260 |
| P19258 | MPV17 | Protein Mpv17 | 25.2 | 18.4 | 1.37 | 0.0173 |

| | | | | | | |
|---|---|---|---|---|---|---|
| Q8CEI3 | CCDC28A | Coiled-coil domain-containing 28A | 32.7 | 23.9 | 1.36 | 0.0159 |
| Q3TVA9 | CCDC136 | Coiled-coil domain-containing protein 136 | 368.5 | 272.4 | 1.35 | 0.0021 |
| Q9R1C7 | PRPF40A | Pre-mRNA-processing factor 40 homolog A | 120.2 | 89.6 | 1.34 | 0.0338 |
| O70496 | CLCN7 | H(+)/Cl(-) exchange transporter 7 | 30.9 | 23.2 | 1.33 | 0.0100 |
| Q9R0N8 | SYT6 | Synaptotagmin-6 | 36.8 | 27.6 | 1.33 | 0.0070 |
| Q8BZR9 | NCBP3 | Nuclear cap-binding protein subunit 3 | 21.3 | 16.0 | 1.33 | 0.0006 |
| P70170-2 | ABCC9 | Isoform SUR2B of ATP-binding cassette sub-family C member 9 | 113.3 | 86.2 | 1.31 | 0.0174 |
| O08842 | GFRA2 | GDNF family receptor alpha-2 | 351.1 | 268.3 | 1.31 | 0.0056 |
| Q4ACU6 | SHANK3 | SH3 and multiple ankyrin repeat domains protein 3 | 1989.3 | 1525.3 | 1.30 | 0.0459 |
| Q64343 | ABCG1 | ATP-binding cassette sub-family G member 1 | 32.0 | 24.5 | 1.30 | 0.0052 |
| Q9DB72 | BTBD17 | BTB/POZ domain-containing protein 17 | 669.3 | 515.4 | 1.30 | 0.0014 |
| P15209 | NTRK2 | BDNF/NT-3 growth factors receptor | 97.0 | 74.8 | 1.30 | 0.0287 |
| P08101 | FCGR2 | Low affinity immunoglobulin gamma Fc region receptor II | 18.7 | 14.5 | 1.29 | 0.0216 |
| P62311 | LSM3 | U6 snRNA-associated Sm-like protein LSm3 | 70.0 | 54.1 | 1.29 | 0.0202 |
| F7AA26 | PAKAP | Paralemmin A kinase anchor protein (Fragment) | 709.8 | 549.1 | 1.29 | 0.0277 |
| P13597 | ICAM1 | Intercellular adhesion molecule 1 | 20.1 | 15.6 | 1.28 | 0.0269 |
| P58058 | NADK | NAD kinase | 32.7 | 25.5 | 1.28 | 0.0204 |
| P06909 | CFH | Complement factor H | 145.5 | 114.0 | 1.28 | 0.0002 |
| O08677 | KNG1 | Kininogen-1 | 55.5 | 43.6 | 1.27 | 0.0476 |
| P47743 | GRM8 | Metabotropic glutamate receptor 8 | 46.9 | 36.9 | 1.27 | 0.0116 |
| Q9JLQ0 | CD2AP | CD2-associated protein | 87.9 | 69.2 | 1.27 | 0.0476 |
| Q6PEE2 | CTIF | Isoform 2 of CBP80/20-dependent translation initiation factor | 66.0 | 52.1 | 1.27 | 0.0494 |
| Q8BIE6-2 | FRMD4A | Isoform 2 of FERM domain-containing protein 4A | 64.3 | 51.0 | 1.26 | 0.0426 |
| A2AAE1-2 | FSA | Isoform 2 of Transmembrane protein KIAA1109 | 141.9 | 113.0 | 1.26 | 0.0150 |
| O88196 | TTC3 | E3 ubiquitin-protein ligase TTC3 | 22.5 | 18.0 | 1.25 | 0.0103 |

| | | | | | | |
|---|---|---|---|---|---|---|
| Q8BWP8 | B4GAT1 | Beta-1,4-glucuronyltransferase 1 | 45.2 | 36.0 | 1.25 | 0.0205 |
| Q9JIK5 | DDX21 | Nucleolar RNA helicase 2 | 48.2 | 38.7 | 1.25 | 0.0204 |
| O89084 | PDE4A | cAMP-specific 3',5'-cyclic phosphodiesterase 4A | 138.3 | 111.1 | 1.24 | 0.0437 |
| P39098 | MAN1A2 | Mannosyl-oligosaccharide 1,2-alpha-mannosidase IB | 112.0 | 90.1 | 1.24 | 0.0073 |
| Q91X58 | ZFAND2B | AN1-type zinc finger protein 2B | 57.1 | 46.0 | 1.24 | 0.0220 |
| Q0VGU4 | VGF | Neurosecretory protein VGF | 767.9 | 619.4 | 1.24 | 0.0417 |
| Q8K377 | LRRTM1 | Leucine-rich repeat transmembrane neuronal protein 1 | 177.1 | 143.9 | 1.23 | 0.0074 |
| Q91UZ1 | PLCB4 | 1-phosphatidylinositol 4,5-bisphosphate phosphodiesterase | 920.5 | 748.9 | 1.23 | 0.0345 |
| Q922J6 | TSN2 | Tetraspanin-2 | 528.5 | 430.2 | 1.23 | 0.0077 |
| P51141 | DVL1 | Segment polarity protein dishevelled homolog DVL-1 | 33.9 | 27.7 | 1.23 | 0.0364 |
| Q6P5D3 | DHX57 | Putative ATP-dependent RNA helicase DHX57 | 76.2 | 62.3 | 1.22 | 0.0251 |
| Q9ER47 | KCNH7 | Potassium voltage-gated channel subfamily H member 7 | 59.4 | 48.8 | 1.22 | 0.0236 |
| Q9Z2B2 | SLC25A14 | Brain mitochondrial carrier protein 1 | 122.6 | 101.0 | 1.21 | 0.0196 |
| Q7TPV4 | MYBBP1A | Myb-binding protein 1A | 39.5 | 32.6 | 1.21 | 0.0363 |
| P63056 | OLFM3 | Noelin-3 | 159.4 | 131.4 | 1.21 | 0.0206 |
| Q9JI39 | ABCB10 | ATP-binding cassette sub-family B member 10, mitochondrial | 217.2 | 179.1 | 1.21 | 0.0309 |
| Q8VBY2 | CAMKK1 | Calcium/calmodulin-dependent protein kinase kinase 1 | 1249.3 | 1031.2 | 1.21 | 0.0064 |
| Q921T2 | TOR1AIP1 | Torsin-1A-interacting protein 1 | 51.5 | 42.6 | 1.21 | 0.0305 |
| Q8VHE0 | SEC63 | Translocation protein SEC63 homolog | 113.1 | 93.6 | 1.21 | 0.0105 |
| Q9Z2C5 | MTM1 | Myotubularin | 37.9 | 31.4 | 1.21 | 0.0073 |
| Q9R049-2 | AMFR | Isoform 2 of E3 ubiquitin-protein ligase AMFR | 39.2 | 32.5 | 1.21 | 0.0320 |
| Q6P542 | ABCF1 | ATP-binding cassette sub-family F member 1 | 766.1 | 636.2 | 1.20 | 0.0051 |
| Q8BJL0 | SMARCAL1 | SWI/SNF-related matrix-associated actin-dependent regulator of chromatin subfamily A-like protein 1 | 60.6 | 50.3 | 1.20 | 0.0191 |
| Q9QYY8 | SPAST | Spastin | 137.7 | 114.4 | 1.20 | 0.0298 |

FC, fold change. FC > 1.20, $p < 0.05$.

**Supplementary Table S4. Significant brain DEPs downregulated in ETP69-treated AD+ mice versus DMSO-injected AD+ mice.**

| Accession | Symbol | Description | AD+ ETP69 (Mean) | AD+ DMSO (Mean) | FC | p-value |
|---|---|---|---|---|---|---|
| Q64518 | ATP2A3 | Sarcoplasmic/endoplasmic reticulum calcium ATPase 3 | 128.0 | 239.2 | -1.87 | 0.0308 |
| P02802 | MT1 | Metallothionein-1 | 932.2 | 1455.5 | -1.56 | 0.0123 |
| P02798 | MT2 | Metallothionein-2 | 254.1 | 370.3 | -1.46 | 0.0189 |
| Q7TMW6-2 | CIAO3 | Isoform 2 of Cytosolic iron-sulfur assembly component 3 | 15.9 | 22.9 | -1.44 | 0.0130 |
| A0A5F8MPE1 | EPB41L3 | Band 4.1-like protein 3 | 350.4 | 500.0 | -1.43 | 0.0449 |
| Q920M7 | SYT17 | Synaptotagmin-17 | 57.3 | 81.6 | -1.42 | 0.0212 |
| A2AQ89 | SHF | SH2 domain-containing adapter protein F (Fragment) | 48.9 | 69.6 | -1.42 | 0.0003 |
| P50172 | HSD11B1 | Corticosteroid 11-beta-dehydrogenase isozyme 1 | 111.1 | 156.1 | -1.41 | 0.0136 |
| Q61001 | LAMA5 | Laminin subunit alpha-5 | 38.2 | 53.5 | -1.40 | 0.0370 |
| P55065 | PLTP | Phospholipid transfer protein | 31.3 | 42.8 | -1.37 | 0.0242 |
| P28667 | MARCKSL1 | MARCKS-related protein | 1633.4 | 2233.7 | -1.37 | 0.0381 |
| Q8VHG2 | AMOT | Angiomotin | 40.3 | 54.6 | -1.35 | 0.0304 |
| D3YZS5 | TARBP2 | RISC-loading complex subunit TARBP2 (Fragment) | 103.7 | 139.4 | -1.34 | 0.0281 |
| O55186 | CD59A | CD59A glycoprotein | 19.8 | 26.4 | -1.33 | 0.0454 |
| Q8K1S4 | UNC5A | Netrin receptor UNC5A | 107.2 | 142.2 | -1.33 | 0.0093 |
| Q8CA72 | GAN | Gigaxonin | 16.1 | 21.0 | -1.30 | 0.0403 |
| P11930 | NUDT19 | Nucleoside diphosphate-linked moiety X motif 19 | 15.8 | 20.5 | -1.29 | 0.0160 |
| Q61599 | ARHGDIB | Rho GDP-dissociation inhibitor 2 | 174.3 | 224.5 | -1.29 | 0.0042 |
| Q6PCN7 | HLTF | Helicase-like transcription factor | 118.1 | 149.7 | -1.27 | 0.0183 |
| Q9CYI0 | NJMU | Protein Njmu-R1 | 27.1 | 34.3 | -1.26 | 0.0278 |
| P07356 | ANXA2 | Annexin A2 | 2047.1 | 2583.4 | -1.26 | 0.0276 |
| P30681 | HMGB2 | High mobility group protein B2 | 136.0 | 169.3 | -1.24 | 0.0140 |

| | | | | | | |
|---|---|---|---|---|---|---|
| Q64522 | H2A2B | Histone H2A type 2-B | 92.4 | 115.0 | -1.24 | 0.0195 |
| Q3UNH4 | GPRIN1 | G protein-regulated inducer of neurite outgrowth 1 | 11647.8 | 14487.9 | -1.24 | 0.0136 |
| P62911 | RPL32 | 60S ribosomal protein L32 | 905.5 | 1115.1 | -1.23 | 0.0040 |
| P16045 | LGALS1 | Galectin-1 | 760.7 | 935.6 | -1.23 | 0.0474 |
| P98156 | VLDLR | Very low-density lipoprotein receptor | 56.5 | 69.2 | -1.22 | 0.0006 |
| Q920R0 | ALS2 | Alsin | 504.0 | 611.8 | -1.21 | 0.0016 |
| Q8BH27 | MEGF9 | Multiple epidermal growth factor-like domains protein 9 | 140.2 | 169.8 | -1.21 | 0.0335 |
| Q9Z2X1 | HNRNPF | Heterogeneous nuclear ribonucleoprotein F | 158.3 | 191.6 | -1.21 | 0.0348 |
| Q9D0J8 | PTMS | Parathymosin | 886.8 | 1069.3 | -1.21 | 0.0051 |

FC, fold change. $|FC| > 1.20$, $p < 0.05$.

**Supplementary Table S5. Significant brain DEPs upregulated in DMSO-injected AD+ mice versus DMSO-injected WT mice.**

| Accession | Symbol | Description | AD+ DMSO (Mean) | WT DMSO (Mean) | FC | p-value |
|---|---|---|---|---|---|---|
| A0A5F8MPM4 | MPZ | Myelin protein P0 | 152.6 | 15.6 | 9.77 | 0.0118 |
| P06880 | GH1 | Somatotropin | 1111.9 | 442.3 | 2.51 | 0.0239 |
| Q9EPF5 | OLFR703 | Olfactory receptor 703 | 1556.1 | 635.0 | 2.45 | 0.0441 |
| Q8CA95 | PDE10A | Isoform 3 of cAMP and cAMP-inhibited cGMP 3',5'-cyclic phosphodiesterase 10A | 671.5 | 274.7 | 2.44 | 0.0006 |
| Q8BIS8 | CCDC126 | Coiled-coil domain-containing protein 126 | 719.3 | 296.4 | 2.43 | 0.0122 |
| P12025 | MDK | Midkine | 58.3 | 25.0 | 2.33 | 0.0064 |
| F7DBB3 | AHNAK2 | AHNAK nucleoprotein 2 (Fragment) | 1709.4 | 751.9 | 2.27 | 0.0047 |
| Q99JH7 | CLSTN3 | Calsyntenin-3 | 187.7 | 82.8 | 2.27 | 0.0031 |
| P16110 | LGALS3 | Galectin-3 | 69.8 | 31.8 | 2.20 | 0.0173 |
| P29788 | VTN | Vitronectin | 318.7 | 148.0 | 2.15 | 0.0000003 |
| Q761V0 | SLC6A5 | Sodium- and chloride-dependent glycine transporter 2 | 733.4 | 384.5 | 1.91 | 0.0295 |
| P08226 | APOE | Apolipoprotein E | 8623.5 | 4647.7 | 1.86 | 0.000003 |
| Q06890 | CLU | Clusterin | 2737.4 | 1477.2 | 1.85 | 0.000002 |
| Q9D920 | BORCS5 | BLOC-1-related complex subunit 5 | 17.1 | 9.6 | 1.78 | 0.0100 |
| P0DOV2 | IFI204 | Interferon-activable protein 204 | 36.9 | 20.8 | 1.78 | 0.0006 |
| H3BJ88 | VMN2R3 | Vomeronasal 2, receptor 3 | 145.9 | 85.1 | 1.71 | 0.0292 |
| P13597 | ICAM1 | Intercellular adhesion molecule 1 | 15.6 | 9.3 | 1.67 | 0.0004 |
| O54983 | CRYM | Ketimine reductase mu-crystallin | 5886.7 | 3703.6 | 1.59 | 0.0048 |
| Q08091 | CNN1 | Calponin-1 | 60.4 | 38.3 | 1.58 | 0.0166 |
| Q9CR57 | RPL14 | 60S ribosomal protein L14 | 1046.2 | 672.7 | 1.56 | 0.0342 |
| P26883 | FKBP1A | Peptidyl-prolyl cis-trans isomerase FKBP1A | 1357.8 | 884.0 | 1.54 | 0.0105 |

| | | | | | | |
|---|---|---|---|---|---|---|
| Q99LB4 | CAPG | Capping protein (Actin filament), gelsolin-like | 195.0 | 127.6 | 1.53 | 0.0012 |
| P01029 | C4B | Complement C4-B | 87.0 | 57.1 | 1.52 | 0.0065 |
| Q9D3X9 | MFAP3L | Microfibrillar-associated protein 3-like | 48.0 | 32.0 | 1.50 | 0.0006 |
| O09159 | MAN2B1 | Lysosomal alpha-mannosidase | 137.9 | 92.4 | 1.49 | 0.0022 |
| Q61001 | LAMA5 | Laminin subunit alpha-5 | 53.5 | 36.0 | 1.49 | 0.0430 |
| P08101 | FCGR2 | Low affinity immunoglobulin gamma Fc region receptor II | 14.5 | 9.9 | 1.47 | 0.0473 |
| Q62422 | OSTF1 | Osteoclast-stimulating factor 1 | 94.8 | 64.8 | 1.46 | 0.0240 |
| Q8K1X1 | WDR11 | WD repeat-containing protein 11 | 134.7 | 92.4 | 1.46 | 0.0430 |
| Q9CYI0 | NJMU-R1 | Protein Njmu-R1 | 34.3 | 23.6 | 1.45 | 0.0009 |
| P17897 | LYZ1 | Lysozyme C-1 | 128.5 | 89.3 | 1.44 | 0.000002 |
| P12265 | GUSB | Beta-glucuronidase | 53.9 | 37.5 | 1.44 | 0.0004 |
| P62878 | RBX1 | E3 ubiquitin-protein ligase RBX1 | 376.8 | 263.3 | 1.43 | 0.0346 |
| P02802 | MT1 | Metallothionein-1 | 1455.5 | 1022.5 | 1.42 | 0.0105 |
| P35969 | FLT1 | Vascular endothelial growth factor receptor 1 | 1000.1 | 704.4 | 1.42 | 0.0011 |
| Q8BLC3 | LYPD1 | Ly6/PLAUR domain-containing protein 1 | 33.5 | 23.6 | 1.42 | 0.0468 |
| Q9D1J1 | NECAP2 | Adaptin ear-binding coat-associated protein 2 | 47.4 | 33.7 | 1.41 | 0.0124 |
| Q6AXF6 | SIDT1 | SID1 transmembrane family member 1 | 95.8 | 68.3 | 1.40 | 0.0021 |
| P07356 | ANXA2 | Annexin A2 | 2583.4 | 1845.5 | 1.40 | 0.0024 |
| Q9JI11 | STK4 | Serine/threonine-protein kinase 4 | 223.9 | 160.1 | 1.40 | 0.0450 |
| Q8BH64 | EHD2 | EH domain-containing protein 2 | 145.4 | 104.2 | 1.40 | 0.0001 |
| Q922S4 | PDE2A | cGMP-dependent 3',5'-cyclic phosphodiesterase | 6307.5 | 4534.3 | 1.39 | 0.0010 |
| Q8BGZ1 | HPCAL4 | Hippocalcin-like protein 4 | 1623.2 | 1169.8 | 1.39 | 0.0391 |
| P28063 | PSMB8 | Proteasome subunit beta type-8 | 52.9 | 38.3 | 1.38 | 0.0084 |
| P57716 | NCSTN | Nicastrin | 78.2 | 57.4 | 1.36 | 0.0002 |
| Q9WUU7 | CTSZ | Cathepsin Z | 1127.0 | 828.3 | 1.36 | 0.00001 |
| O08992 | SDCBP | Syntenin-1 | 652.2 | 479.6 | 1.36 | 0.0051 |

| | | | | | | |
|---|---|---|---|---|---|---|
| P49710 | HCLS1 | Hematopoietic lineage cell-specific protein | 148.6 | 109.8 | 1.35 | 0.0063 |
| A2TJV2 | PALM3 | Paralemmin-3 | 76.0 | 56.8 | 1.34 | 0.0023 |
| Q3TBL6 | TNFAIP8L3 | Tumor necrosis factor alpha-induced protein 8-like protein 3 | 150.9 | 113.3 | 1.33 | 0.0055 |
| O35417 | PDYN | Proenkephalin-B | 116.7 | 87.7 | 1.33 | 0.0302 |
| P11438 | LAMP1 | Lysosome-associated membrane glycoprotein 1 | 455.8 | 344.2 | 1.32 | 0.000004 |
| Q61599 | ARHGDIB | Rho GDP-dissociation inhibitor 2 | 224.5 | 171.5 | 1.31 | 0.0052 |
| Q8CA71 | SHISA4 | Protein shisa-4 | 88.3 | 67.5 | 1.31 | 0.0390 |
| P59108 | CPNE2 | Copine-2 | 95.6 | 73.4 | 1.30 | 0.0194 |
| Q5U5V2 | HYKK | Hydroxylysine kinase | 83.9 | 64.4 | 1.30 | 0.0008 |
| Q920M7 | SYT17 | Synaptotagmin-17 | 81.6 | 62.7 | 1.30 | 0.0347 |
| Q9R1J0 | NSDHL | Sterol-4-alpha-carboxylate 3-dehydrogenase, decarboxylating | 55.5 | 42.7 | 1.30 | 0.0227 |
| P50285 | FMO1 | Dimethylaniline monooxygenase [N-oxide-forming] 1 | 32.2 | 24.8 | 1.30 | 0.0091 |
| P11835 | ITGB2 | Integrin beta-2 | 321.8 | 248.6 | 1.29 | 0.0007 |
| Q64669 | NQO1 | NAD(P)H dehydrogenase [quinone] 1 | 484.9 | 374.6 | 1.29 | 0.0362 |
| F7BX42 | NPTXR | Neuronal pentraxin receptor | 997.2 | 771.0 | 1.29 | 0.0465 |
| E9Q236 | ABCC4 | ATP-binding cassette, sub-family C (CFTR/MRP), member 4 | 47.7 | 36.9 | 1.29 | 0.0007 |
| Q9CQE1 | NIPSNAP3B | Protein NipSnap homolog 3B | 921.9 | 713.2 | 1.29 | 0.0003 |
| Q8R143 | PTTG1IP | Pituitary tumor-transforming gene 1 protein-interacting protein | 45.0 | 34.8 | 1.29 | 0.0329 |
| P20060 | HEXB | Beta-hexosaminidase subunit beta | 2238.0 | 1739.9 | 1.29 | 0.0033 |
| P61514 | RPL37A | 60S ribosomal protein L37a | 1595.8 | 1241.6 | 1.29 | 0.0095 |
| Q8R4V2 | DUSP15 | Dual specificity protein phosphatase 15 | 239.3 | 186.5 | 1.28 | 0.0106 |
| Q920R0 | ALS2 | Alsin | 611.8 | 477.0 | 1.28 | 0.0028 |
| Q9D379 | EPHX1 | Epoxide hydrolase 1 | 1054.2 | 822.9 | 1.28 | 0.00004 |
| Q9DCJ9 | NPL | N-acetylneuraminate lyase | 418.2 | 326.5 | 1.28 | 0.0023 |
| Q91WG7 | DGKG | Diacylglycerol kinase gamma | 926.6 | 724.0 | 1.28 | 0.0126 |
| P14069 | S100A6 | Protein S100-A6 | 238.7 | 186.6 | 1.28 | 0.0134 |

| | | | | | | |
|---|---|---|---|---|---|---|
| Q8VHC3 | SELENOM | Selenoprotein M | 115.1 | 90.4 | 1.27 | 0.0320 |
| P97372 | PSME2 | Proteasome activator complex subunit 2 | 394.8 | 310.2 | 1.27 | 0.0006 |
| Q9WV32 | ARPC1B | Actin-related protein 2/3 complex subunit 1B | 198.3 | 155.9 | 1.27 | 0.0003 |
| P29416 | HEXA | Beta-hexosaminidase subunit alpha | 298.4 | 234.9 | 1.27 | 0.0006 |
| P28798 | GRN | Progranulin | 437.0 | 345.0 | 1.27 | 0.0072 |
| Q8K215 | LYRM4 | LYR motif-containing protein 4 | 119.1 | 94.2 | 1.27 | 0.0064 |
| O09114 | PTGDS | Prostaglandin-H2 D-isomerase | 3154.4 | 2507.3 | 1.26 | 0.0027 |
| P70202 | LXN | Latexin | 1483.0 | 1179.2 | 1.26 | 0.0063 |
| P05555 | ITGAM | Integrin alpha-M | 112.8 | 89.8 | 1.26 | 0.0009 |
| Q61809 | LRRN1 | Leucine-rich repeat neuronal protein 1 | 41.2 | 32.9 | 1.26 | 0.0176 |
| P01887 | B2M | Beta-2-microglobulin | 79.3 | 63.3 | 1.25 | 0.0011 |
| E9PWM3 | ARMCX4 | Armadillo repeat-containing, X-linked 4 | 37.8 | 30.3 | 1.25 | 0.0073 |
| P34914 | EPHX2 | Bifunctional epoxide hydrolase 2 | 931.7 | 746.6 | 1.25 | 0.00003 |
| Q920E5 | FDPS | Farnesyl pyrophosphate synthase | 1422.9 | 1145.6 | 1.24 | 0.00001 |
| Q9WTR5 | CDH13 | Cadherin-13 | 1857.1 | 1500.7 | 1.24 | 0.0488 |
| P51880 | FABP7 | Fatty acid-binding protein, brain | 699.5 | 565.3 | 1.24 | 0.0398 |
| O70456 | SFN | 14-3-3 protein sigma | 5980.2 | 4834.2 | 1.24 | 0.00001 |
| P18242 | CTSD | Cathepsin D | 2314.3 | 1872.5 | 1.24 | 0.00004 |
| Q9ERE7 | MESD | LRP chaperone MESD | 780.6 | 633.4 | 1.23 | 0.0014 |
| Q8VHL0 | SLC14A1 | Isoform 2 of Urea transporter 1 | 187.1 | 152.0 | 1.23 | 0.0049 |
| Q9DBG5 | PLIN3 | Perilipin-3 | 765.2 | 624.0 | 1.23 | 0.0401 |
| P55302 | LRPAP1 | Alpha-2-macroglobulin receptor-associated protein | 1667.4 | 1360.1 | 1.23 | 0.0051 |
| Q91XE4 | ACY3 | N-acyl-aromatic-L-amino acid amidohydrolase (carboxylate-forming) | 102.2 | 83.4 | 1.23 | 0.0218 |
| Q3UNZ8 | CRYZL2 | Quinone oxidoreductase-like protein 2 | 642.0 | 524.5 | 1.22 | 0.0383 |
| Q8C7K6 | PCYOX1L | Prenylcysteine oxidase-like | 446.3 | 364.9 | 1.22 | 0.0007 |

| | | | | | | |
|---|---|---|---|---|---|---|
| A3KFX0 | NT5C1A | Cytosolic 5'-nucleotidase 1A | 142.0 | 116.2 | 1.22 | 0.0061 |
| Q9QZ08 | NAGK | N-acetyl-D-glucosamine kinase | 189.1 | 154.9 | 1.22 | 0.0009 |
| O70370 | CTSS | Cathepsin S | 33.7 | 27.6 | 1.22 | 0.0035 |
| Q91W43 | GLDC | Glycine dehydrogenase (decarboxylating), mitochondrial | 301.2 | 247.7 | 1.22 | 0.0258 |
| Q3SXD3 | HDDC2 | 5'-deoxynucleotidase HDDC2 | 38.0 | 31.3 | 1.21 | 0.0338 |
| Q9DBL2 | GDAP2 | Isoform 2 of Ganglioside-induced differentiation-associated protein 2 | 84.7 | 69.8 | 1.21 | 0.0125 |
| Q76LS9 | MINDY1 | Ubiquitin carboxyl-terminal hydrolase MINDY-1 | 566.5 | 466.9 | 1.21 | 0.0260 |
| Q920P5 | AK5 | Adenylate kinase isoenzyme 5 | 1707.4 | 1408.5 | 1.21 | 0.0492 |
| Q60854 | SERPINB6 | Serpin B6 | 3546.6 | 2929.7 | 1.21 | 0.0043 |
| P59672 | ANKS1A | Isoform 2 of Ankyrin repeat and SAM domain-containing protein 1A | 45.0 | 37.2 | 1.21 | 0.0184 |
| Q61112 | SDF4 | 45 kDa calcium-binding protein | 194.4 | 160.8 | 1.21 | 0.0057 |
| Q6NS52 | DGKB | Diacylglycerol kinase beta | 1997.7 | 1652.2 | 1.21 | 0.0074 |
| P10639 | TXN | Thioredoxin | 3409.9 | 2821.1 | 1.21 | 0.0403 |
| Q8R1B5 | CPLX3 | Complexin-3 | 383.8 | 317.9 | 1.21 | 0.0079 |
| Q8BR86 | KIRREL3 | Kin of IRRE-like protein 3 | 77.8 | 64.5 | 1.21 | 0.0296 |
| Q00915 | CRBPI | Retinol-binding protein 1 | 945.8 | 785.8 | 1.20 | 0.0004 |
| O88456 | CAPNS1 | Calpain small subunit 1 | 726.6 | 603.6 | 1.20 | 0.0054 |
| P51910 | APOD | Apolipoprotein D | 824.5 | 686.0 | 1.20 | 0.000004 |
| Q920A5 | SCPEP1 | Retinoid-inducible serine carboxypeptidase | 227.1 | 189.1 | 1.20 | 0.0004 |

FC, fold change. FC > 1.20, $p < 0.05$.

**Supplementary Table S6. Significant brain DEPs downregulated in DMSO-injected AD$^+$ mice versus DMSO-injected WT mice**.

| Accession | Symbol | Description | AD$^+$ DMSO (Mean) | WT DMSO (Mean) | FC | p-value |
|---|---|---|---|---|---|---|
| P04919 | SLC4A1 | Band 3 anion transport protein | 18.5 | 49.5 | -2.67 | 0.0311 |
| P10637-3 | MAPT | Isoform Tau-B of Microtubule-associated protein tau | 23.5 | 43.4 | -1.85 | 0.0026 |
| Q64327 | MEA1 | Male-enhanced antigen 1 | 41.1 | 74.5 | -1.81 | 0.0145 |
| Q31125 | SLC39A7 | Zinc transporter SLC39A7 | 142.2 | 220.2 | -1.55 | 0.0073 |
| O08842 | GFRA2 | GDNF family receptor alpha-2 | 268.3 | 392.4 | -1.46 | 0.0005 |
| P00397 | MTCO1 | Cytochrome c oxidase subunit 1 | 171.4 | 249.3 | -1.45 | 0.0441 |
| Q9R1C7 | PRPF40A | Pre-mRNA-processing factor 40 homolog A | 89.6 | 130.2 | -1.45 | 0.0064 |
| Q9D287 | BCAS2 | Pre-mRNA-splicing factor SPF27 | 38.4 | 55.8 | -1.45 | 0.0038 |
| Q922K7 | NOP2 | Probable 28S rRNA (cytosine-C(5))-methyltransferase | 28.9 | 41.7 | -1.44 | 0.0372 |
| Q02111 | PRKCQ | Protein kinase C theta type | 72.7 | 104.0 | -1.43 | 0.0482 |
| Q91UZ1 | PLCB4 | 1-phosphatidylinositol 4,5-bisphosphate phosphodiesterase | 748.9 | 1067.3 | -1.43 | 0.0026 |
| Q6DID5 | PWWP3A | PWWP domain-containing DNA repair factor 3A | 23.0 | 32.7 | -1.42 | 0.0216 |
| Q64143 | PIK3R3 | Phosphatidylinositol 3-kinase regulatory subunit gamma | 201.2 | 285.0 | -1.42 | 0.0206 |
| O89084 | PDE4A | cAMP-specific 3',5'-cyclic phosphodiesterase 4A | 111.1 | 156.8 | -1.41 | 0.0275 |
| P49025 | CIT | Citron Rho-interacting kinase | 1845.0 | 2569.0 | -1.39 | 0.0018 |
| Q8BG67-2 | EFR3A | Isoform 2 of Protein EFR3 homolog A | 81.8 | 113.1 | -1.38 | 0.0005 |
| Q8BZ94 | ZMAT4 | Zinc finger matrin-type protein 4 | 108.6 | 149.9 | -1.38 | 0.0110 |
| Q9R0C8 | VAV3 | Guanine nucleotide exchange factor VAV3 | 29.9 | 40.9 | -1.37 | 0.0002 |
| Q8BYW9 | EOGT | EGF domain-specific O-linked N-acetylglucosamine transferase | 14.5 | 19.8 | -1.37 | 0.0220 |
| Q3TVA9 | CCDC136 | Coiled-coil domain-containing protein 136 | 272.4 | 371.9 | -1.37 | 0.0021 |
| E0CXZ8 | DNM3 | Dynamin-3 | 124.1 | 169.3 | -1.36 | 0.0292 |
| Q9CWZ3 | RBM8A | RNA-binding protein 8A | 90.1 | 121.5 | -1.35 | 0.0297 |

| | | | | | | |
|---|---|---|---|---|---|---|
| H3BJD6 | PPP1R9A | Protein phosphatase 1, regulatory subunit 9A | 948.1 | 1272.3 | -1.34 | 0.0284 |
| F7BNZ5 | BCAS1 | Breast carcinoma-amplified sequence 1 homolog (Fragment) | 935.6 | 1253.4 | -1.34 | 0.0281 |
| Q99LT0 | DPY30 | Protein dpy-30 homolog | 27.4 | 36.5 | -1.33 | 0.0105 |
| Q99KX1 | MLF2 | Myeloid leukemia factor 2 | 663.3 | 883.5 | -1.33 | 0.0209 |
| Q60803 | TRAF3 | TNF receptor-associated factor 3 | 1148.7 | 1522.3 | -1.33 | 0.0006 |
| Q91X72 | HPX | Hemopexin | 599.6 | 793.3 | -1.32 | 0.0203 |
| Q9DB72 | BTBD17 | BTB/POZ domain-containing protein 17 | 515.4 | 676.3 | -1.31 | 0.0001 |
| Q7TSJ2 | MAP6 | Microtubule-associated protein 6 | 12016.1 | 15755.0 | -1.31 | 0.0255 |
| Q8C1M2 | ZNF428 | Zinc finger protein 428 | 26.0 | 34.1 | -1.31 | 0.0055 |
| E9Q8T1 | TACC2 | Transforming acidic coiled-coil-containing protein 2 | 118.3 | 155.1 | -1.31 | 0.0385 |
| Q5BLK4 | TUT7 | Terminal uridylyltransferase 7 | 35.5 | 46.3 | -1.31 | 0.0252 |
| Q9CW07 | PPP1R3G | Protein phosphatase 1 regulatory subunit 3G | 36.1 | 47.2 | -1.31 | 0.0116 |
| Q8C0P5 | CORO2A | Coronin-2A | 512.2 | 667.3 | -1.30 | 0.0051 |
| Q99MR1 | GIGYF1 | GRB10-interacting GYF protein 1 | 30.9 | 40.3 | -1.30 | 0.0039 |
| Q61464 | ZNF638 | Zinc finger protein 638 | 297.4 | 385.6 | -1.30 | 0.0251 |
| Q7TPV4 | MYBBP1A | Myb-binding protein 1A | 32.6 | 42.0 | -1.29 | 0.0004 |
| O70496 | CLCN7 | H(+)/Cl(-) exchange transporter 7 | 23.2 | 29.9 | -1.29 | 0.0014 |
| E9Q5K9 | YTHDC1 | YTH domain-containing protein 1 | 143.8 | 184.3 | -1.28 | 0.0199 |
| B2M1R6 | HNRNPK | Heterogeneous nuclear ribonucleoprotein K | 4963.5 | 6346.6 | -1.28 | 0.0230 |
| Q91VR8 | BRK1 | Protein BRICK1 | 81.7 | 103.8 | -1.27 | 0.0385 |
| Q8BJH1 | ZC2HC1A | Zinc finger C2HC domain-containing protein 1A | 1421.4 | 1805.7 | -1.27 | 0.0308 |
| Q923D5 | WBP11 | WW domain-binding protein 11 | 207.6 | 263.6 | -1.27 | 0.0085 |
| O09044 | SNAP23 | Synaptosomal-associated protein 23 | 496.7 | 628.1 | -1.26 | 0.0462 |
| Q810U5 | CCDC50 | Coiled-coil domain-containing protein 50 | 77.3 | 97.7 | -1.26 | 0.0256 |
| Q9QX47 | SON | Protein SON | 274.4 | 346.3 | -1.26 | 0.0089 |
| Q14B02 | RNF133 | E3 ubiquitin-protein ligase RNF133 | 100.7 | 126.9 | -1.26 | 0.0157 |

| | | | | | | |
|---|---|---|---|---|---|---|
| Q8K2F8 | LSM14A | Protein LSM14 homolog A | 202.5 | 255.0 | -1.26 | 0.0097 |
| Q9QXL2-4 | KIF21A | Isoform 4 of Kinesin-like protein KIF21A | 29.8 | 37.5 | -1.26 | 0.0218 |
| Q8VDM6 | HNRNPUL1 | Heterogeneous nuclear ribonucleoprotein U-like protein 1 | 422.0 | 531.1 | -1.26 | 0.0345 |
| P09405 | NCL | Nucleolin | 13088.4 | 16463.8 | -1.26 | 0.0153 |
| Q9JJP2 | TP73 | Tumor protein p73 | 61.6 | 77.3 | -1.26 | 0.0396 |
| P98191 | CDS1 | Phosphatidate cytidylyltransferase 1 | 107.1 | 134.4 | -1.26 | 0.0412 |
| P13864 | DNMT1 | DNA (cytosine-5)-methyltransferase 1 | 147.1 | 184.3 | -1.25 | 0.0259 |
| Q8VD73 | KCNAB3 | Potassium voltage-gated channel, shaker-related subfamily, beta member 3 | 374.3 | 468.5 | -1.25 | 0.0068 |
| Q8BI72 | CDKN2AIP | CDKN2A-interacting protein | 20.1 | 25.1 | -1.25 | 0.0301 |
| Q8VHM5 | HNRNPR | Heterogeneous nuclear ribonucleoprotein R | 1712.3 | 2141.8 | -1.25 | 0.0232 |
| Q9ET77 | JPH3 | Junctophilin-3 | 246.4 | 308.2 | -1.25 | 0.0128 |
| B2KF50 | UHRF1BP1 | UHRF1 (ICBP90)-binding protein 1 | 99.5 | 124.4 | -1.25 | 0.0032 |
| Q8K327 | CHAMP1 | Chromosome alignment-maintaining phosphoprotein 1 | 113.3 | 141.2 | -1.25 | 0.0101 |
| O88532 | ZFR | Zinc finger RNA-binding protein | 306.1 | 380.6 | -1.24 | 0.0156 |
| Q9DBS9 | OSBPL3 | Oxysterol-binding protein-related protein 3 | 41.5 | 51.4 | -1.24 | 0.0015 |
| Q8R4U7 | LUZP1 | Leucine zipper protein 1 | 646.4 | 798.9 | -1.24 | 0.0433 |
| Q9D824 | FIP1L1 | Pre-mRNA 3'-end-processing factor FIP1 | 132.0 | 162.8 | -1.23 | 0.0359 |
| Q8BG81 | POLDIP3 | Polymerase delta-interacting protein 3 | 551.7 | 680.1 | -1.23 | 0.0329 |
| Q69Z26 | CNTN4 | Contactin-4 | 162.4 | 199.5 | -1.23 | 0.0072 |
| Q64343 | ABCG1 | ATP-binding cassette subfamily G member 1 | 24.5 | 30.1 | -1.23 | 0.0007 |
| E9Q7G0 | NUMA1 | Nuclear mitotic apparatus protein 1 | 480.2 | 589.2 | -1.23 | 0.0148 |
| Q62376 | SNRNP70 | U1 small nuclear ribonucleoprotein 70 kDa | 814.1 | 996.0 | -1.22 | 0.0363 |
| A2AJI0 | MAP7D1 | MAP7 domain-containing protein 1 | 1580.6 | 1933.2 | -1.22 | 0.0437 |
| Q9ESK9 | RB1CC1 | RB1-inducible coiled-coil protein 1 | 202.4 | 247.5 | -1.22 | 0.0061 |
| Q9D2U5 | NAA38 | N-alpha-acetyltransferase 38, NatC auxiliary subunit | 55.1 | 67.3 | -1.22 | 0.0203 |

| | | | | | | |
|---|---|---|---|---|---|---|
| Q922J6 | TSPAN2 | Tetraspanin-2 | 430.2 | 526.0 | -1.22 | 0.0131 |
| Q61730 | IL1RAP | Interleukin-1 receptor accessory protein | 260.4 | 318.4 | -1.22 | 0.0005 |
| Q9Z2D6-2 | MECP2 | Isoform B of Methyl-CpG-binding protein 2 | 1963.4 | 2398.3 | -1.22 | 0.0417 |
| A2AAE1-2 | K1109 | Isoform 2 of Transmembrane protein KIAA1109 | 113.0 | 138.0 | -1.22 | 0.0092 |
| Q80YR5 | SAFB2 | Scaffold attachment factor B2 | 113.8 | 138.8 | -1.22 | 0.0429 |
| Q62172 | RALBP1 | RalA-binding protein 1 | 68.3 | 83.3 | -1.22 | 0.0108 |
| Q9JJF3 | RIOX1 | Ribosomal oxygenase 1 | 39.9 | 48.6 | -1.22 | 0.0116 |
| Q9CY66 | GAR1 | H/ACA ribonucleoprotein complex subunit 1 | 169.1 | 205.6 | -1.22 | 0.0011 |
| Q80TY0 | FNBP1 | Formin-binding protein 1 | 990.4 | 1203.4 | -1.22 | 0.0153 |
| F8VQ70 | SCAPER | S phase cyclin A-associated protein in the ER | 143.0 | 173.7 | -1.21 | 0.0215 |
| P62311 | LSM3 | U6 snRNA-associated Sm-like protein LSm3 | 54.1 | 65.6 | -1.21 | 0.0061 |
| P26339 | CHGA | Chromogranin-A | 238.9 | 289.9 | -1.21 | 0.0257 |
| Q99LE6 | ABCF2 | ATP-binding cassette sub-family F member 2 | 395.7 | 479.5 | -1.21 | 0.0010 |
| Q8K4R4 | PITPNC1 | Cytoplasmic phosphatidylinositol transfer protein 1 | 1453.6 | 1761.1 | -1.21 | 0.0137 |
| Q5XG69 | FAM169A | Soluble lamin-associated protein of 75 kDa | 414.2 | 500.8 | -1.21 | 0.0207 |
| P30999 | CTNND1 | Catenin delta-1 | 870.9 | 1052.9 | -1.21 | 0.0270 |
| Q6NV83 | U2SURP | U2 snRNP-associated SURP motif-containing protein | 299.4 | 361.8 | -1.21 | 0.0185 |
| Q67BT3 | SLC13A5 | Solute carrier family 13 member 5 | 172.1 | 207.9 | -1.21 | 0.0306 |
| Q0P678 | ZC3H18 | Zinc finger CCCH domain-containing protein 18 | 119.7 | 144.5 | -1.21 | 0.0429 |
| O55003 | BNIP3 | BCL2/adenovirus E1B 19 kDa protein-interacting protein 3 | 37.2 | 44.9 | -1.21 | 0.0354 |
| O88196 | TTC3 | E3 ubiquitin-protein ligase TTC3 | 18.0 | 21.6 | -1.20 | 0.0065 |
| Q9WV18 | GABBR1 | Gamma-aminobutyric acid type B receptor subunit 1 | 1516.0 | 1826.1 | -1.20 | 0.0306 |
| Q8K296 | MTMR3 | Myotubularin-related protein 3 | 30.5 | 36.8 | -1.20 | 0.0468 |
| Q5DTX6 | JCAD | Junctional protein associated with coronary artery disease | 170.4 | 205.2 | -1.20 | 0.0376 |
| Q66L44 | CBARP | Voltage-dependent calcium channel beta subunit-associated regulatory protein | 548.2 | 658.7 | -1.20 | 0.0073 |

| | | | | | | |
|---|---|---|---|---|---|---|
| Q56A08 | GPKOW | G-patch domain and KOW motifs-containing protein | 50.1 | 60.2 | -1.20 | 0.0369 |
| P63056 | OLFM3 | Noelin-3 | 131.4 | 157.8 | -1.20 | 0.0080 |

FC, fold change. $|FC| > 1.20$, $p < 0.05$.

**Supplementary Table S7. Significant brain DEPs upregulated in ETP69-treated WT mice versus DMSO-injected WT mice.**

| Accession | Symbol | Description | WT ETP69 (Mean) | WT DMSO (Mean) | FC | p-value |
|---|---|---|---|---|---|---|
| Q7TPD0 | INTS3 | Integrator complex subunit 3 | 103.5 | 28.1 | 3.69 | 0.0230 |
| A0A5F8MPM4 | MPZ | Myelin protein P0 | 56.7 | 15.6 | 3.63 | 0.0026 |
| Q61646 | HP | Haptoglobin | 280.5 | 83.5 | 3.36 | 0.0448 |
| F7DBB3 | AHNAK2 | AHNAK nucleoprotein 2 (Fragment) | 1799.3 | 751.9 | 2.39 | 0.0076 |
| P15209-2 | NTRK2 | Isoform GP95-TRKB of BDNF/NT-3 growth factors receptor | 152.2 | 76.4 | 1.99 | 0.0066 |
| P13597 | ICAM1 | Intercellular adhesion molecule 1 | 18.2 | 9.3 | 1.95 | 0.00004 |
| Q91X72 | HPX | Hemopexin | 1522.1 | 793.3 | 1.92 | 0.0107 |
| P08101 | FCGR2 | Low affinity immunoglobulin gamma Fc region receptor II | 17.8 | 9.9 | 1.81 | 0.0304 |
| P04370 | MBP | Isoform 9 of Myelin basic protein | 689.4 | 389.0 | 1.77 | 0.0477 |
| Q8CA95 | PDE10A | Isoform 3 of cAMP and cAMP-inhibited cGMP 3',5'-cyclic phosphodiesterase 10A | 483.6 | 274.7 | 1.76 | 0.0129 |
| P62806 | H4C1 | Histone H4 | 13994.0 | 8668.7 | 1.61 | 0.0004 |
| P29699 | AHSG | Alpha-2-HS-glycoprotein | 158.6 | 99.4 | 1.60 | 0.0020 |
| Q60829 | PPP1R1B | Protein phosphatase 1 regulatory subunit 1B | 476.0 | 298.4 | 1.60 | 0.0013 |
| Q9D920 | BORCS5 | BLOC-1-related complex subunit 5 | 15.1 | 9.6 | 1.57 | 0.0113 |
| A2AJK6 | CHD7 | Chromodomain-helicase-DNA-binding protein 7 | 36.8 | 23.7 | 1.55 | 0.0233 |
| Q8CG46 | SMC5 | Structural maintenance of chromosomes protein 5 | 84.3 | 54.6 | 1.54 | 0.0371 |
| Q08091 | CNN1 | Calponin-1 | 57.6 | 38.3 | 1.50 | 0.0159 |
| Q9CXJ4 | ABCB8 | Mitochondrial potassium channel ATP-binding subunit | 2075.0 | 1428.1 | 1.45 | 0.0250 |
| Q91VJ2 | CAVIN3 | Caveolae-associated protein 3 | 59.3 | 41.2 | 1.44 | 0.0458 |
| Q61147 | CP | Ceruloplasmin | 264.5 | 184.8 | 1.43 | 0.00005 |
| Q9Z277 | BAZ1B | Tyrosine-protein kinase BAZ1B | 1762.1 | 1237.8 | 1.42 | 0.0059 |
| Q7TNV1 | TLCD3B | Ceramide synthase | 26.0 | 18.4 | 1.42 | 0.0095 |
| P17897 | LYZ1 | Lysozyme C-1 | 124.9 | 89.3 | 1.40 | 0.0007 |

| | | | | | | |
|---|---|---|---|---|---|---|
| Q62422 | OSTF1 | Osteoclast-stimulating factor 1 | 90.6 | 64.8 | 1.40 | 0.0333 |
| Q60709 | APLP2 | Amyloid-like protein 2 | 21.8 | 15.7 | 1.39 | 0.0084 |
| Q9JKY7 | CYP2D22 | Cytochrome P450 CYP2D22 | 47.2 | 34.2 | 1.38 | 0.0240 |
| P84244 | H3-3a | Histone H3.3 | 1107.6 | 815.5 | 1.36 | 0.0062 |
| O54983 | CRYM | Ketimine reductase mu-crystallin | 5005.1 | 3703.6 | 1.35 | 0.0278 |
| P0DOV2 | IFI204 | Interferon-activable protein 204 | 28.1 | 20.8 | 1.35 | 0.0431 |
| Q640M6 | GDPD5 | Glycerophosphodiester phosphodiesterase domain-containing protein 5 | 26.4 | 19.6 | 1.34 | 0.0121 |
| P07356 | ANXA2 | Annexin A2 | 2411.6 | 1845.5 | 1.31 | 0.0020 |
| Q8BJ05 | ZC3H14 | Zinc finger CCCH domain-containing protein 14 | 118.6 | 90.8 | 1.31 | 0.0032 |
| Q8BJ05 | DGKB | Diacylglycerol kinase beta | 2152.9 | 1652.2 | 1.30 | 0.0194 |
| Q91WG7 | DGKG | Diacylglycerol kinase gamma | 940.1 | 724.0 | 1.30 | 0.0061 |
| Q9Z321 | TOP3B | DNA topoisomerase 3-beta-1 | 30.3 | 23.8 | 1.27 | 0.0396 |
| P98086 | C1QA | Complement C1q subcomponent subunit A | 236.0 | 185.8 | 1.27 | 0.0379 |
| O70250 | PGAM2 | Phosphoglycerate mutase 2 | 120.2 | 94.6 | 1.27 | 0.0083 |
| Q80W37 | SNUPN | Snurportin-1 | 45.5 | 36.3 | 1.26 | 0.0076 |
| Q91XE4 | ACY3 | N-acyl-aromatic-L-amino acid amidohydrolase (carboxylate-forming) | 104.4 | 83.4 | 1.25 | 0.0125 |
| Q61285 | ABCD2 | ATP-binding cassette, sub-family D, member 2 | 44.4 | 35.5 | 1.25 | 0.0334 |
| E9Q236 | ABCC4 | ATP-binding cassette, sub-family C (CFTR/MRP), member 4 | 46.1 | 36.9 | 1.25 | 0.0041 |
| P35969 | FLT1 | Vascular endothelial growth factor receptor 1 | 877.7 | 704.4 | 1.25 | 0.0001 |
| Q8BYJ6 | TBC1D4 | TBC1 domain family member 4 | 24.6 | 19.8 | 1.24 | 0.0456 |
| Q8C788 | SNX18 | Sorting nexin-18 | 102.9 | 82.9 | 1.24 | 0.0063 |
| Q00519 | XDH | Xanthine dehydrogenase/oxidase | 93.4 | 75.4 | 1.24 | 0.0177 |
| Q922S4 | PDE2A | cGMP-dependent 3',5'-cyclic phosphodiesterase | 5603.8 | 4534.3 | 1.24 | 0.0240 |
| P28667 | MARCKSL1 | MARCKS-related protein | 2276.7 | 1858.4 | 1.23 | 0.0340 |

| | | | | | | |
|---|---|---|---|---|---|---|
| Q6P1B3 | PIANP | PILR alpha-associated neural protein | 71.2 | 58.1 | 1.22 | 0.0364 |
| Q9DCB4 | ARPP21 | Isoform 3 of cAMP-regulated phosphoprotein 21 | 113.2 | 92.4 | 1.22 | 0.0198 |
| A2TJV2 | PALM3 | Paralemmin-3 | 69.2 | 56.8 | 1.22 | 0.0169 |
| Q9DCJ9 | NPL | N-acetylneuraminate lyase | 397.6 | 326.5 | 1.22 | 0.0055 |
| Q9D1Q4 | DPM3 | Dolichol-phosphate mannosyltransferase subunit 3 | 56.6 | 46.5 | 1.22 | 0.0121 |
| Q8K215 | LYRM4 | LYR motif-containing protein 4 | 114.3 | 94.2 | 1.21 | 0.0497 |
| P06909 | CFH | Complement factor H | 118.0 | 97.3 | 1.21 | 0.0242 |
| P56380 | NUDT2 | Bis(5'-nucleosyl)-tetraphosphatase [asymmetrical] | 53.5 | 44.1 | 1.21 | 0.0197 |
| P14106 | C1QB | Complement C1q subcomponent subunit B | 596.7 | 494.2 | 1.21 | 0.0293 |
| E9QA16 | CALD1 | Caldesmon 1 | 128.0 | 106.2 | 1.21 | 0.0262 |
| Q9D379 | EPHX1 | Epoxide hydrolase 1 | 990.1 | 822.9 | 1.20 | 0.0014 |
| Q9CY18 | SNX7 | Sorting nexin-7 | 89.8 | 74.8 | 1.20 | 0.0380 |

FC, fold change. FC > 1.20, $p < 0.05$.

**Supplementary Table S8. Significant brain DEPs downregulated in ETP69-treated WT mice versus DMSO-injected WT mice.**

| Accession | Symbol | Description | WT ETP69 (Mean) | WT DMSO (Mean) | FC | p-value |
|---|---|---|---|---|---|---|
| Q6PFX9 | TNKS | Poly [ADP-ribose] polymerase tankyrase-1 | 10.9 | 16.8 | -1.54 | 0.0013 |
| P00397 | MTCO1 | Cytochrome c oxidase subunit 1 | 166.1 | 249.3 | -1.50 | 0.0278 |
| Q02111 | PRKCQ | Protein kinase C theta type | 73.8 | 104.0 | -1.41 | 0.0344 |
| O88428 | PAPSS2 | Bifunctional 3'-phosphoadenosine 5'-phosphosulfate synthase 2 | 13.4 | 18.1 | -1.36 | 0.0490 |
| Q9QXV0 | PCSK1N | ProSAAS | 417.8 | 561.1 | -1.34 | 0.0253 |
| B0F2B4 | NLGN4L | Neuroligin 4-like | 67.3 | 90.4 | -1.34 | 0.0500 |
| Q6ZPR4 | KCNT1 | Potassium channel subfamily T member 1 | 22.0 | 29.0 | -1.32 | 0.0197 |
| Q8BZ94 | ZMAT4 | Zinc finger matrin-type protein 4 | 113.7 | 149.9 | -1.32 | 0.0031 |
| Q8BLE7 | SLC17A6 | Vesicular glutamate transporter 2 | 1193.2 | 1571.1 | -1.32 | 0.0085 |
| Q8C3X4 | GUF1 | Translation factor Guf1, mitochondrial | 53.4 | 70.0 | -1.31 | 0.0015 |
| Q9CWZ3 | RBM8A | RNA-binding protein 8A | 95.1 | 121.5 | -1.28 | 0.0451 |
| Q8VD73 | KCNAB3 | Potassium voltage-gated channel, shaker-related subfamily, beta member 3 | 368.5 | 468.5 | -1.27 | 0.0011 |
| Q8R242 | CTBS | Di-N-acetylchitobiase | 21.3 | 26.9 | -1.27 | 0.0069 |
| Q80U93 | NUP214 | Nuclear pore complex protein Nup214 | 31.3 | 39.4 | -1.26 | 0.0472 |
| Q9CW07 | PPP1R3G | Protein phosphatase 1 regulatory subunit 3G | 38.0 | 47.2 | -1.24 | 0.0451 |
| Q3TVA9 | CCDC136 | Coiled-coil domain-containing protein 136 | 300.8 | 371.9 | -1.24 | 0.0191 |
| B1AUH5 | CASK | Peripheral plasma membrane CASK (Fragment) | 50.6 | 62.2 | -1.23 | 0.0003 |
| Q5SQY2 | BOD1 | Biorientation of chromosomes in cell division protein 1 | 60.7 | 74.5 | -1.23 | 0.0299 |
| Q9DCP2 | SLC38A3 | Sodium-coupled neutral amino acid transporter 3 | 63.9 | 78.3 | -1.23 | 0.0033 |
| Q5XG69 | FAM169A | Soluble lamin-associated protein of 75 kDa | 408.8 | 500.8 | -1.22 | 0.0046 |
| Q8BUL6 | PLEKHA1 | Pleckstrin homology domain-containing family A member 1 | 30.0 | 36.7 | -1.22 | 0.0115 |

| | | | | | | |
|---|---|---|---|---|---|---|
| Q8K296 | MTMR3 | Myotubularin-related protein 3 | 30.2 | 36.8 | -1.22 | 0.0290 |
| Q9D0D3 | MTPAP | Poly(A) RNA polymerase, mitochondrial | 39.5 | 47.9 | -1.21 | 0.0453 |
| A0A140LIW3 | FRMPD3 | FERM and PDZ domain-containing 3 | 64.3 | 78.1 | -1.21 | 0.0429 |
| P60755 | MDGA2 | MAM domain-containing glycosylphosphatidylinositol anchor protein 2 | 148.9 | 180.5 | -1.21 | 0.0002 |

FC, fold change. $|FC| > 1.20$, $p < 0.05$
